# Mutation count and mutation profile analyses of single-nucleotide variants in single human colon crypts

**DOI:** 10.64898/2026.06.11.731472

**Authors:** Zarko Manojlovic, Cindy Okitsu, Timothy Okitsu, Jordan Wlodarczyk, Michael R. Lieber, Yong Hwee Eddie Loh, Chih-Lin Hsieh

## Abstract

Genetic heterogeneity due to the accumulation of mutations in normal tissues can increase cancer risk and be an important factor in many age-related degenerative and chronic diseases. Somatic mutations can arise from DNA damage or replication errors and accumulate in normal tissues with age. We obtained high depth whole genome sequencing data to comprehensively profile somatic mutations in 106 single human colon crypts with matched bulk controls from 21 individuals age 10 months to 90 years old. Our analysis reveals that about half of the human crypts are polyclonal (multi-lineage) instead of entirely monoclonal as conventionally construed. Consequently, the DNA mutation count would be inflated, while the variant allele frequency for each variant would be reduced in the cell population of a multi-lineage colon crypt. Therefore, our mutation count analysis includes using single stem cell lineage colon crypts exclusively to establish the somatic mutation rate for each of the 96 trinucleotide mutation categories. In addition, the mutation profile, representing the relative presence of the 96 trinucleotide mutation categories, in each colon crypt is analyzed. Unlike mutation count, the mutation profile is not affected by crypt clonality and is very similar across all ages in individuals with no chemotherapy or radiation treatment. Importantly, combined results from mutation count and mutation profile analyses of individuals with chemotherapy or radiation suggest an intriguing impact of these treatments on cell survival. The baseline mutation rate and the normal mutation profile established in our study provide a framework for a genomic standard to assess biological age and deviations associated with factors such as lifestyle, age-related degenerative and chronic diseases, and potentially cancer treatment outcome. Future studies similar to this current study on other tissues can provide further insight into how and why different tissues age differently in humans.

## INTRODUCTION

The human intestine is lined by millions of colon crypts, and human intestinal stem cells (hISCs) reside in a stem cell niche at the base of each colon crypt. These hISCs constantly regenerate the cell lining of each colon crypt, which is replaced as often as every three to five days (1, 2, 3). hISCs divide to renew the stem cells (SCs) in the niche and to populate the transit-amplifying (TA) cell pool to form the lining of each crypt. TA cells proliferate, differentiate as they migrate upward, and eventually shed into the colon lumen after reaching the orifice of the crypt (4, 5). It has been suggested that each colon crypt, which consists of ∼1,500 cells, is populated by multiple stem cells in early development and becomes a single stem cell lineage monoclone in early life due to neutral drift both in mouse and human (2, 6, 7, 8, 9, 10). Based on this assumption, human colon crypts have been used as a model mini-organ system, representing single stem cells, to identify somatic mutations (11, 12, 13). We successfully generated 106 single human colon crypt libraries using our novel whole genome sequencing (WGS) library preparation method. High-quality genomic sequences with ≥30X depth post-alignment were produced from these libraries without ex *vivo* tissue culture, whole genome amplification, or DNA purification (14). Somatic DNA mutations identified in these single crypts include mutations in the stem cell that gives rise to each crypt and mutations that occur during the first few proliferative divisions of the TA cells forming the crypt lining. Our dataset is the first of its kind in providing high quality WGS data for a comprehensive count and profile of mutations in multiple colon crypts from each of the 21 individuals spanning the human lifespan from 10-month to 90-year-old. Our study provides new insights into hISC biology as well as naturally occurring somatic mutations and their repair mechanisms (12, 15, 16).

If each colon crypt were monoclonal, as inferred in previous studies using single genetic markers for lineage tracing by others (for a review, see 5), somatic mutations from the original SC progenitor should be present in all cells in the crypt lining. Therefore, the variant allele frequency (VAF) of these somatic mutations should be 0.5, given that a somatic mutation is typically present only on one allele in a diploid genome. The rationale and data previously shown by our group and others indicate that the VAF distribution profile representing thousands of variants in a single SC lineage crypt can be used to determine whether a cell population is monoclonal or polyclonal (12,17). Unexpectedly, VAF analyses of the 106 single colon crypts in our study showed that only 52 crypts are of single stem cell (1SC) lineage and 54 crypts are of multi-stem cell (multi-SC) lineage (12). Mutation count is inflated in a multi-SC colon crypt because the different somatic mutations harbored by each stem cell lineage are additive, leading to a larger sum of the count than if it were a 1SC colon crypt. Our critical observation on colon crypt clonality clearly indicates that the rate of mutation accumulation with age is most accurately estimated from monoclonal cell populations, such as colon crypts of 1SC lineage. Mutation count increases predominantly linearly with age and can be used to assess biological age once an accurate mutation rate is established. Having an accurate biological age assessment would facilitate future studies on aging and how factors, such as lifestyle, chronic diseases, and medical treatments, influence aging.

The mutation profile, which represents the relative presence (in %) of the 96 mutation categories in each sample, is a qualitative illustration with quantitative measurement of somatic mutation types. Mutation profile analysis of the 106 colon crypts in our study clearly demonstrates that the mutation profile is consistent in different individuals of different ages, unlike the mutation count. The mutation profile can be used to evaluate the impact of extrinsic factors on somatic mutation and to provide insight into the biological relevance of that impact. Here, we report the mutation rate for each of the conventional 96 trinucleotide mutation categories established from 1SC lineage colon crypts of different ages. We also establish a baseline mutation profile from normal colon crypts. Importantly, we demonstrate that both mutation count (which quantifies the accumulation of mutation at a given age) and mutation profile (which measures the balance of mutation categories) are essential for the understanding of aging and eventual insight into biological processes. We present intriguing suggestions of the impact of cancer treatment on mutation categories and the potential biological effect of these treatments from our analysis of individuals with chemotherapy or chemo/radiation combination treatment.

## MATERIALS AND METHODS

### Tissue collection, crypt isolation, sequencing library construction, and whole genome sequencing

Collection of tissue from normal portions of colorectal samples, crypt isolation, and sequencing library construction have been described in detail previously (12, 14). A total of 106 crypt libraries and 21 bulk libraries are constructed from 21 individuals of age 10-month to 90-year-old. These sequencing libraries are pooled after quality control and then sequenced on seven S4 flow cells using NovaSeq 6000 (Illumina, San Diego) S4 300 cycles reagent kit (v1.5) in the Keck Genomics Platform (KGP) Core facility at USC.

### Variant calling

The sequencing reads are assessed and processed using a validated in-house pipeline from the KGP based on GATK v4.2 best practices as described previously (12, 14). The sequencing reads are also processed using the DRAGEN Somatic v 4.2.4 pipeline (Illumina Inc) as described previously (12, 14). PCR duplicate reads were identified and excluded from downstream variant-calling steps. Two germline callers, HaplotypeCaller and FreeBayes (v1.2.0), and two somatic variant callers, Mutect2 and Strelka (v2.9.0), are used in the KGP pipeline. Somatic variant calls in the crypt samples are made with the matched bulk tissue sample from the same individual as the control. The shared somatic variant calls from Mutect2 and Strelka (named 2Caller) through the KGP pipeline and the DRAGEN pipeline have a high inter-caller concordance as previously reported (15). The variant calls from 2Caller are used to increase variant call confidence for all downstream analyses. The variant calls from Illumina’s DRAGEN somatic pipeline are only used for internal validation of the analyses.

### Workflow for variant call filtering

To increase the confidence of variant calls from 2Caller, the 413,323 total SNV calls by the 2Caller are further filtered for downstream analyses of SNV. The non-autosomal variants are excluded because the presentation of sex chromosomes differs between males and females, the mitochondrial DNA is exposed to different mutational conditions than nuclear DNA, and the variants at unknown genomic locations are less clear to interpret. Only autosomal SNV variants supported by ≥15 total sequencing reads are included. Somatic mutations are unlikely to be shared by different cells unless the two cells share the same progenitor. A unique SNV call list is generated by removing variants that appear in more than a single crypt (apparent sharing) either in a single individual or in multiple individuals. We previously found that unique variants in the 2Caller variant call list can still be shared by other crypts or individuals but do not appear on the call list (hidden inter-sharing; 15, 16). These hidden inter-sharing events are more likely artifacts rather than true mutations and must be removed. The final list of SNVs is generated by removing the hidden inter-sharing events after using a script we developed previously to identify them from FreeBayes VCFs (https://github.com/twewyttst777/VariantDuplicateCheck) and as described previously (15, 16).

### Mutation count analysis

Mutation count is essential for evaluating the association of DNA mutations with human aging. The final SNV list is converted into 96-channel single-base substitution trinucleotide-context matrices using SigProfilerMatrixGenerator with the GRCh38 reference genome. Matrix generation is performed in genome mode without exome or BED-based restriction (18, 19). Each SNV is classified into one of the conventional 96 trinucleotide mutation categories, defined by the six pyrimidine-normalized base-substitution classes and the immediate 5′ and 3′ sequence context as described above. The resulting 96 trinucleotide mutation count matrix is used to obtain the mutation count for each trinucleotide category in each crypt.

Average and standard deviation of the mutation count for total as well as for each of the 96 trinucleotide categories are calculated from the 5 or 6 crypts for each individual. The average mutation count for each individual and the average mutation count in each mutation category across different groups of individuals are performed. In addition to analyzing the mutation count across all 21 individuals in our colon crypt cohort, groups of individuals or crypts with specific features that may potentially affect the mutation count in the colon crypts are analyzed separately and then compared. Our analyses of mutation count are comprehensively presented in four groups: Group 1) all crypts from all individuals (All; 21 individuals, 106 crypts); Group 2) all crypts from all individuals with no treatment (no treatment; 14 individuals, 70 crypts); Group 3) all single stem cell lineage crypts from all individuals (1SC crypts;16 individuals, 52 crypts); and Group 4) all single stem cell lineage crypts from individuals with no treatment (1SC no treatment, 11 individuals, 36 crypts). It is important to note that some individuals have no single stem cell lineage crypts.

The slope (mutation rate per year) and R^2^ are calculated from a linear regression model for the average mutation count, either for the total or for each mutation category against age. A t-test of the regression slope is used to determine whether the slope is zero, based on the resulting p-value. The results of the mutation count analysis from the four Groups are further compared to decipher biological indications.

The expected age-appropriate mutation count for each mutation category can be calculated using the intercept and slope derived from Group 4 (1SC crypts, no treatment). The expected mutation count for each treated 1SC crypt is calculated for each mutation category as: age-appropriate count = y-intercept + (slope × age). The difference in mutation count is then calculated by subtracting the expected age-appropriate count from the observed count (observed – age-appropriate). This difference can be used to evaluate category-specific deviations from the expected normal count in each crypt.

### Mutation profile analyses

Mutation profile represents the relative presentation (in %) of each mutation type among the 96 trinucleotide mutation categories in a single crypt. The mutation profile is generated by dividing the mutation count of each mutation category by the total mutation count in each colon crypt. Therefore, the percentage for each mutation category from each individual is analyzed and presented. The mutation profile analyses include the analysis of all mutations and of each mutation category. Again, individuals and crypts with specific features that may affect the mutation profile are analyzed separately across nine Groups, which are then compared.

The nine Groups analyzed and presented are: Group A) all crypts from all individuals (all 21 individuals; 106 crypts); Group B) all crypts from individuals with no treatment (14 individuals; 70 crypts); Group C) 1SC lineage crypts from adults with no treatment (5 individuals; 12 crypts); Group D) adults with no treatment (7 individuals; 35 crypts); Group E) multiple-lineage SC crypts from adults with no treatment (6 individuals; 23 crypts); Group F) young children 0.8 to 4 years old (5 individuals, 25 crypts); Group G) 1SC lineage crypts from young children (4 individuals; 17 crypts); Group H) adults with treatment (7 individuals, 36 crypts); and Group I) 1SC lineage crypts from adults with treatment (5 individuals, 16 crypts).

The slope and R^2^ are calculated, and a t-test is used to evaluate the linear relationship between age and the representation of each mutation category. The results of the mutation profile analysis from the nine Groups are further compared to evaluate the biological significance of the mutation profile.

## RESULTS

### Strategy for quantitative mutation count analyses

The goal of this study is to analyze new somatic mutations occurring in single human colon crypts to assess the mutation accumulation with human aging and to measure the mutation rate in human epithelial cells. A total of 413,323 single nucleotide variants (SNVs) are called by both Mutect2 and Strelka (designated as 2Caller, see Materials and Methods) in 106 human colon crypts from 21 individuals of ages 10 months to 90 years old. After filtering for autosomal SNV supported by ≥15 total sequencing reads, variants shared by two or more colon crypts either in a single individual or in multiple individuals are removed to reduce mutations already present in the stem cells prior to colon crypt formation and to remove false positive variants. A list of 336,210 unique autosomal SNVs is obtained. It is important to further remove hidden inter-sharing variants, which are false positive variant calls that exist in different individuals but only appear in a single crypt on the unique variant list, prior to the downstream analysis (see Methods; 15, 16). After removing variants with hidden inter-sharing from the 336,210 unique variant list, 262,802 SNVs were included in the final analysis presented here.

Among the 21 individuals, four adults received chemotherapy and three adults received a chemotherapy and radiation combination treatment (chemo/radiation) prior to the tissue collection. It is known that these therapies can potentially damage DNA in normal cells, leading to additional mutations, even though the colon crypts were isolated from the normal portion of the colon distant from the neoplasm. Contrary to conventional perception and as discussed in the Introduction, we discovered that colon crypts are frequently polyclonal (multi-SC lineage) as determined by the variant allele frequency analysis of these colon crypts in our previous study (12). Potentially, the number of SNVs identified in a polyclonal colon crypt may be inflated because different mutations exist and new mutations can arise in each of the multiple stem cells in the crypt; importantly, these mutations are additive in the mutation count tabulation of the crypt. However, some of the new mutations in a polyclonal colon crypt may not meet the variant call criteria due to low alternate allele read count. These complexities lead to uncertainty when the mutation count is the focus of the analysis. Our analyses of mutation count are comprehensively presented in four Groups as described in Materials and Methods above.

Mutation count in single colon crypts of single 1SC lineage from individuals of different ages provides the least confounded estimate of the somatic mutation rate in human colon epithelium in this dataset. As a convention, 96 possible trinucleotide mutation categories with the middle base being the mutation site is used for qualitative single base substitution mutation analysis involving different mechanisms underlying DNA damage and repair. By analyzing mutation count across the 96 mutation categories in individuals of different ages, we provide detailed quantitative information on the contribution of each category to the overall mutation rate in human colon crypts. This detailed analysis is also presented in the four groups as described above. The goal of these analyses is to determine whether mutation accumulation is linear with human age and to establish a normal baseline of mutation with age.

### Somatic single nucleotide mutations accumulate with age in normal human colon crypts

A total of 262,802 autosomal SNVs with ≥15 read depth are included in the analysis after removal of both apparent and hidden sharing variants (as mentioned above and in Supplementary Table S1). A linear trend (R^2^ = 0.7771) of increased SNV count with age, with an estimated mutation rate of 54.49 SNV per year, is observed when all 106 crypts are included (Group 1) in the analysis (Fig. 1A, upper panel graph). All six individuals with average SNV counts above the trend line are those with either chemotherapy or chemo/radiation treatment (Fig. 1A, green and orange dots in the upper panel graph). As mentioned above, both chemotherapy and radiation can potentially increase DNA damage and lead to somatic mutations. After removing the seven individuals with treatment from the analysis, the remaining 70 colon crypts from the 14 individuals with no treatment (Group 2) showed a better fitting linear trend of mutation accumulation with age (R^2^ increased to 0.9597) with an estimated mutation rate of 46.80 SNVs per year (Fig. 1A, lower panel graph).

**Figure 1.**
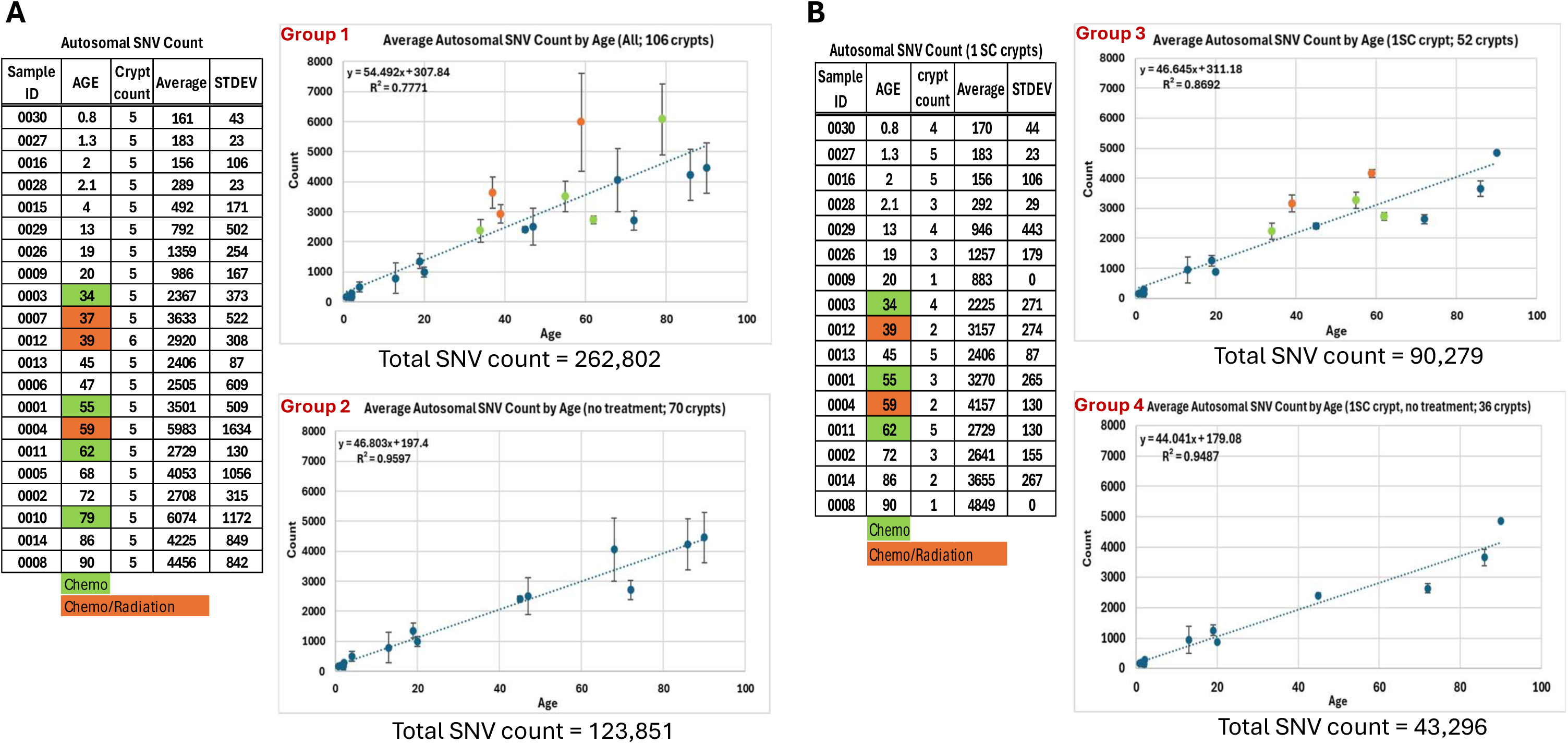
SNV count increase in human colon crypts with age. A) Average of autosomal SNV count in all colon crypts of each individual and standard deviation (STDEV) are summarized and graphed. The top panel graph includes all 106 crypts from 21 individuals. The bottom panel graph includes 70 crypts from 14 individuals with no treatment. B) Average of autosomal SNV count in all 1SC crypts from each individual and standard deviation (STDEV) are summarized and graphed. The top panel graph includes 52 1SC crypts from 16 individuals. The bottom panel graph only includes 36 1SC crypts from 11 individuals with no treatment. The individuals with chemotherapy are marked by green highlight and the individuals with chemo/radiation combination treatment are marked by orange highlight as indicated below the Summary table. Please refer to Supplementary Figure S1 For the same graphs using SNV count of each crypt instead of average SNV count.

As discussed above, human colon crypts are not always monoclonal as conventionally inferred previously (12). Stem cells from different lineages can acquire new somatic mutations over time and complicate the variant calls and the variant counts. The SNV counts in the 52 1SC crypts from 16 individuals (Group 3) showed a slightly better fit for a linear trend (R^2^ = 0.8692) than data including all 106 crypts (where R^2^ = 0.7771) with an estimated mutation rate of 46.64 SNVs per year (Fig. 1B, upper panel graph). It is noteworthy that the deviation of the SNV counts among the 1SC crypts compared to all crypts within the same individual is much reduced in most of the individuals. Four of the five individuals with higher than trend line average SNV counts are individuals with chemotherapy or chemo/radiation treatments. The remaining individual with a higher SNV count with no treatment only has a single 1SC crypt (the remaining crypts sequenced from this individual are multi-SC crypts); therefore, the higher count may not be representative for that individual.

Overall, the higher average SNV counts observed in some individuals is not solely due to the crypt clonality from these individuals. Instead, the chemotherapy and/or radiation treatment contributed to the increase in mutation count in these individuals. About one-third of all the colon crypts (36 of 106 crypts) are 1SC crypts from 11 individuals with no treatment (Group 4). The R^2^ based on the SNV counts from these 36 crypts (R^2^ = 0.9487) showed strong support for a linear mutation accumulation at an estimated rate of 44.04 SNVs per year (Fig. 1B, lower panel of the plots). The same analyses based on autosomal SNV counts of each crypt, instead of average autosomal SNV count of each individual, showed similar results with lower estimated mutation rates for Groups 3 and 4 than based on the average count of each individual (Supplementary Figure S1).

Taken together, the clonality of the crypts and medical treatment can influence the estimated mutation rate. All six individuals with average SNV counts higher than the trend line are individuals with chemotherapy or chemo/radiation treatment. A mutation rate of >54 SNVs per year estimated from all 106 colon crypts (Group 1) is clearly inflated by individuals with chemotherapy and chemo/radiation treatment, as well as crypt clonality. The mutation rate estimated based on average SNV counts in the 70 crypts from individuals with no treatment (Group 2) and in the 52 1SC crypts from 16 individuals (Group 3) are nearly the same at ∼47 mutations per year. The most accurate mutation rate should be based on the SNV counts in the 36 1SC crypts from 11 individuals with no treatment (Group 4) at 44 SNV mutations per year. The mutation rates estimated based on autosomal SNV counts of each crypt (Supplementary Fig. S1) are similar to the mutation rates estimated based on the average SNV count from each individual in all four groups (Fig. 1). Potential causes of large variation among the 5 crypts within each of five specific individuals with no treatment (Fig. 1A Group 2, sample IDs 0005, 0006, 0008, 0014; and 0029) are 1) a likely event of colon crypt fission (sample ID 0029); and 2) clonality of the crypt (sample IDs 0005, 0006, 0008, and 0014).

### C to T change in the 5’-CpG configuration has the highest mutation rate

To further dissect the association of somatic mutation with age, the count of SNV calls from the 2Caller are analyzed for each of the 96 mutation categories (Supplementary Table S2). The mean SNV count for each of the 96 mutation categories from each individual is plotted relative to the age of each individual and a linear regression model is used to calculate the slope (Supplementary Fig. S2). The slope calculated for each of the 96 plots is the estimated per-year mutation rate for the specific mutation category. The contribution of each mutation category to the accumulation of mutations with age can be determined from the slope.

As mentioned above, medical treatment and colon crypt clonality can influence the mutations included in the mutation analysis. Therefore, four groups of analyses are presented: all 106 crypts; 70 crypts from individuals with no treatment; 52 1SC crypts; and 36 1SC crypts from individuals with no treatment (Fig. 2). Based on a t-test of the slope in the linear regression model, a few mutation categories showed no age-associated increase when all individuals, including those with treatment, are included in the analysis (Groups 1 and 3, Fig. 2A and 2C, p-value ≥0.05 highlighted in orange color). When only individuals with treatment are excluded in the analysis (Groups 2 and 4, Fig. 2B and 2D), all 96 mutation categories, except one with a marginal p-value of 0.051, showed significant linear increases of mutation count. These findings indicate that counts of all mutation categories increase linearly with age at different rates in normal individuals with only one exception with a borderline p-value of 0.051.

**Figure 2.**
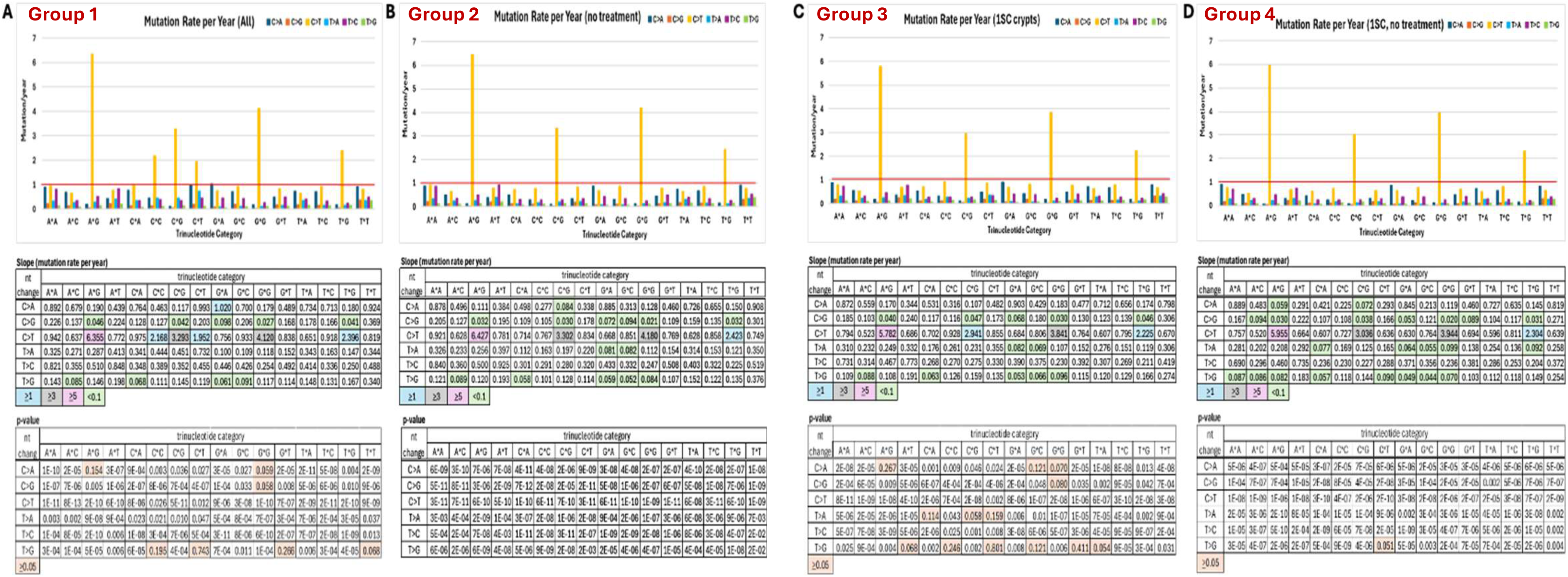
Mutation rate of nucleotides in 96 mutation trinucleotide categories. The estimated mutation rate for nucleotide changes from the slope of the linear regression model based on the mean SNV counts in each individual are graphed and summarized as groups of: A) all 106 crypts from 21 individuals in the study; B) 70 crypts from 14 individuals with no treatment; C) 52 single stem cell lineage crypts from 16 individuals; D) 36 single stem cell lineage crypts from 11 individuals with no treatment. The graph at the top of each panel shows the mean SNV counts in each of the 96 trinucleotide categories. The * in the X-axis denotes the location of the SNV with the nucleotide changes indicated in the top right of the graph by different colors. The y-axis indicates the mutation rate based on the slope calculated from the mean SNV counts in each of 96 trinucleotide categories based on a linear regression model. The red horizontal line indicates mutation rate of 1 per year in the graphs. The slopes and the p-values from t-test on the slope of the regression line are summarized below the graph. The keys for the highlighted numbers are below each summary when applicable.

It is clear that the C as the nucleotide in the middle of NCG (N = any nucleotide) configuration changing to T has the highest mutation rate among the 96 categories in all four groups of analyses (Fig.2, histogram bars above red horizontal lines in the graph at the top of each panel and pink, blue, and grey highlights in the Slope summary below the graph in each panel). This C>T preference is consistent with findings in many previous studies. We also provide further details of the contribution from each type of nucleotide changes to the overall mutation rate (Table 1). While most of the 96 trinucleotide categories of mutation showed a linear trend of increased SNV counts with age, some categories of mutation have a nearly negligible mutation rate of <0.1 per year (Fig. 2, green highlights).

**Table 1.** Contribution of nucleotide changes to the overall mutation rate.

| nt<br>change | % of total |  |  |  |
| --- | --- | --- | --- | --- |
|  | Group 1 | Group 2 | Group 3 | Group 4 |
| **C>A** | 17.4% | 15.6% | 17.2% | 16.1% |
| **C>G** | 4.4% | 4.1% | 4.6% | 4.2% |
| **C>T** | 52.4% | 54.5% | 52.7% | 55.8% |
| **T>A** | 8.7% | 7.0% | 7.6% | 6.3% |
| **T>C** | 13.2% | 14.6% | 13.5% | 13.5% |
| **T>G** | 4.0% | 4.3% | 4.5% | 4.2% |

Group1 - 106 crypts from 21 individuals
Group2 - 70 crypts from 14 individuals w/ no treatment
Group3 - 52 single SClineage crypts from 16 individuals
Group4 - 36 single SClineage crypts from 11 individuals w/ no treatment

### Strategy for qualitative mutation profile analyses

The mutation profile, with the convention of presenting the percentage of mutation in each mutation category, illustrates the relative presence of the 96 mutation categories in each sample. The mutation profile has been used to assess the impact of abnormal conditions, such as neoplastic change or exogenous (environmental or therapeutic) mutagenesis. The mutation count will always increase as an individual ages, assuming no significant number of backward mutations occurs at the existing mutation sites. However, the mutation profile may not change without marked influence of any extrinsic factor that alters the mutation rate of one or more mutation categories. Mutation count analysis and mutation profile analysis provide different but complementary information about the effects of intrinsic aging and extrinsic factors on somatic mutations. Both mutation count analysis and the mutation profile analysis are necessary to assess whether deviation from normal aging occurs and what is the effect of the abnormal factor. If a factor affects the mutation rate of many mutation categories, the mutation profile of the individual may show no change despite the mutation count being higher than the age-appropriate mutation count. Similarly, an individual with an age-appropriate total mutation count may have an abnormal mutation profile due to extrinsic factors affecting specific mutation categories. Comparison of mutation counts in a specific sample to a baseline standard established from normal individuals of different ages would allow further assessment of deviation from normal aging.

The mutation profile reflects the relative presentation among the mutation categories within a single sample. It is likely that the mutation profile does not vary with age, since the mechanisms underlying mutation are not expected to differ when no abnormal or extrinsic factor is present. We analyzed mutation profiles from individuals of different ages, with and without medical treatment, and colon crypts of different clonalities to determine whether the mutation profile changes with age. For various important comparisons, the mutation profile analysis is presented here as nine different Groups as described in the Materials and Methods.

### Mutation profile from individuals of various ages reveals limited changes with age

Mutation profile was generated for each individual (Supplementary Figure S3). Overall, mutation profile is visually similar across all individuals with more notable changes in individuals with chemotherapy or with a chemo/radiation combination treatment (sample ID 0001, 0003, 0004, 0007, 0010, 0011, and 0012). The most remarkable and consistent difference is the lower presence of C>T mutations at the NCG sites (N = any nucleotide) of individuals with treatment. It is worth noting that variation within an individual appears to be higher in young children (sample ID 0015, 0016, 0027, 0028, and 0030) as indicated by larger standard deviation among the multiple crypts in many mutation categories from the same individual. However, this may be the effect of a small number of mutations for percentage calculation, leading to higher fluctuation.

To further assess the association of mutation profile with age, the average percentage of SNV in each mutation category from each individual is plotted by age and a linear regression model is used to calculate the slope, which represents the change of mutation profile per year (Supplementary Fig. S4). When all 106 crypts from 21 individuals of various ages (Group A) are included, most of the 96 mutation categories had no clearly discernible increase in percentage of SNV with age and many mutation categories even show a decrease with age (Supplementary Fig. S4). Likewise, very little change in mutation profile with age was observed in individuals with no treatment (Group B; Supplementary Fig. S4). The p-value of t-test on the slope in most of the 96 mutation categories showed no support for mutation profile change with age (Fig. 3), in contrast to the mutation count analysis (Fig. 2). The mutation profile and the mutation count analyses clearly illustrate mutation count increases linearly in all the 96 mutation categories with age while the mutation profile distribution among the 96 mutation categories remains similar with age.

**Figure 3.**
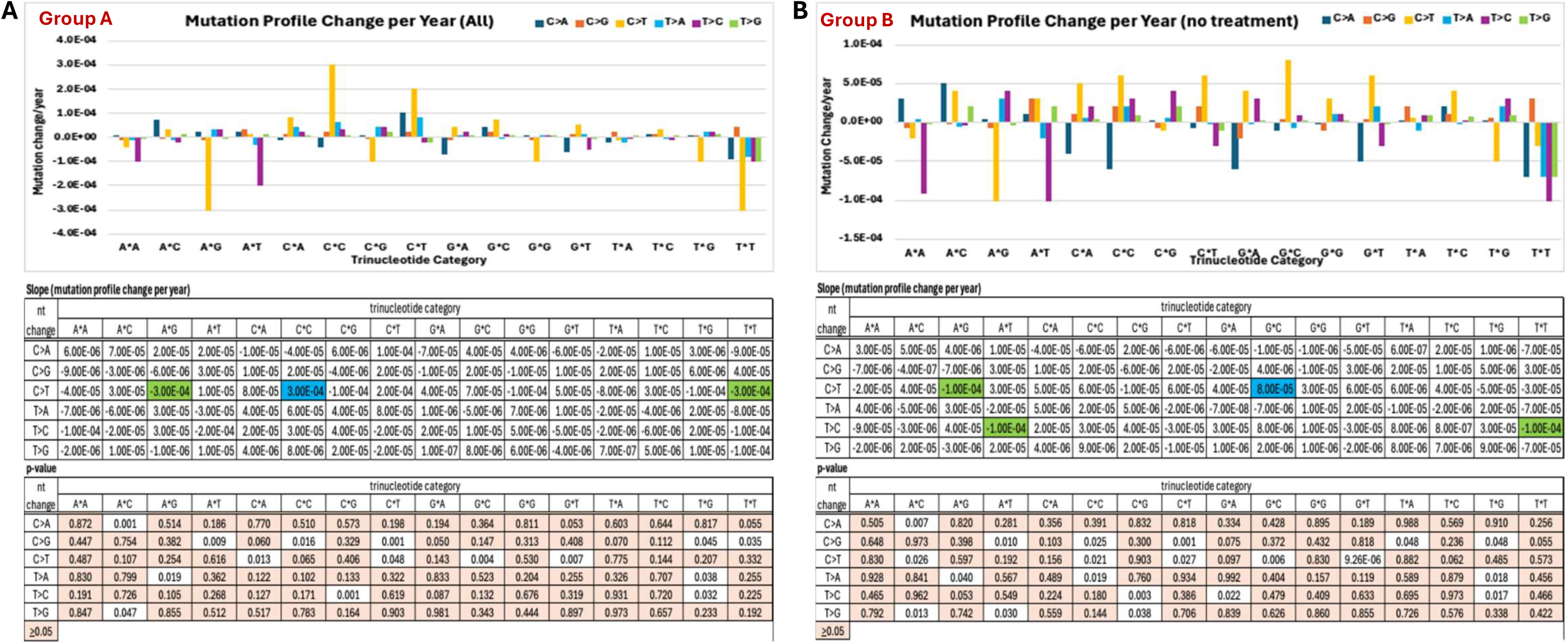
Summary of mutation profile change in 96 mutation trinucleotide categories. The estimated mutation profile changes from the slope of the linear regression model based on the mean SNV distribution of 96 mutation categories in each individual are graphed and summarized as two groups: A) all 106 crypts from 21 individuals in the study; B) 70 crypts from 14 individuals with no treatment. The * in the X-axis indicates the location of the SNV with the nucleotide changes indicated in the top right of the graph by different colors. In the graph on the top of each panel, the y-axis indicates the change of distribution per year based on the slope calculated from the mean SNV percentage in each of 96 trinucleotide categories based on a linear regression model (see Supplementary Fig. S4). The slopes and the p-values from t-test on the slope of the regression line are summarized below the graph. The blue highlight indicates the largest slope and the green highlight indicates the smallest slope in the slope summary. The p-value ≥0.05 is highlighted in orange.

### Stem cell clonality of the colon crypts does not affect the mutation profile

Stem cell clonality affects mutation count as described and demonstrated above. However, the number of stem cells in a colon crypt is not expected to affect the mutation profile because the mutation rate of each mutation category should be the same across stem cells residing in the same colon crypt. To examine whether stem cell clonality impacts the mutation profile, we first compared the mutation profiles of 1SC colon crypts from adults with no treatment (Group C) and that of all colon crypts from adults with no treatment (Group D). There is no visually discernible difference between the two groups (Fig. 4A and Fig. 4B). The fold difference of each mutation category between the two groups is calculated by dividing the percentage of mutation in 1SC colon crypts by the percentage of mutation in all colon crypts from adults with no treatment (Fig. 4C). All 96 categories differ less than 2-fold, and the difference is more likely due to random fluctuation and the relatively small sample size. We then further compared the 1SC crypts (Group C) to the multi-SC crypts (Group E) from adults with no treatment (Fig. 5). Similar to the findings from the first comparison, there is less than 2-fold difference between the 1SC crypts and multi-SC crypts from adults with no treatment in all 96 mutation categories (Fig. 5B). These analyses clearly indicate the clonality of the colon crypts does not change the mutation profile even though the mutation count is clearly affected. These findings also affirm the hypothesis that stem cell clonality affects the quantity of the detected mutation but does not alter the mutation profile.

**Figure 4.**
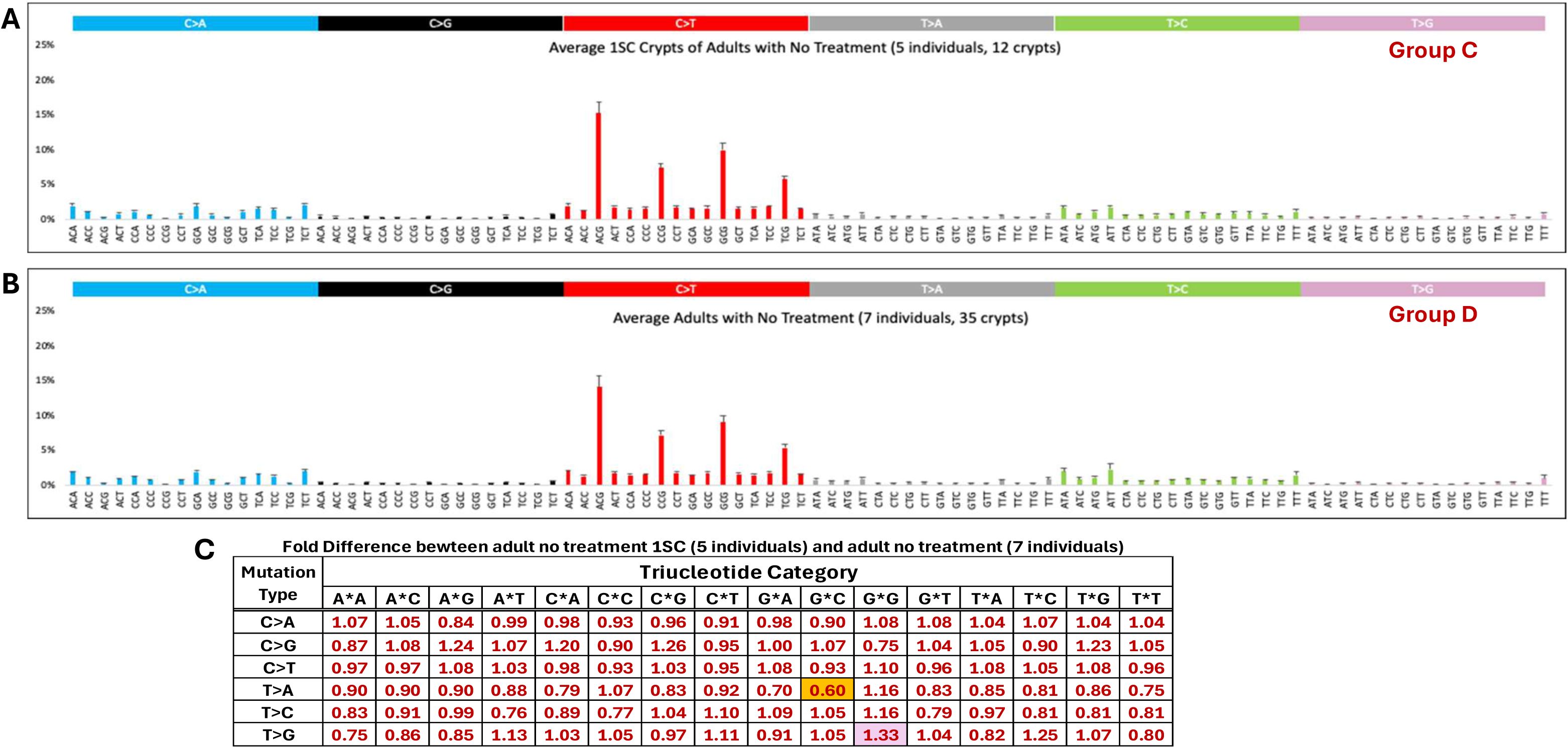
Mutation profile comparison between single stem cell (1SC) colon crypts and all colon crypts from adults with no treatment. A) Mutation profile of average mutation percentage of each of the 96 mutation categories from 5 adults (12 1SC crypts) with no treatment is presented with standard deviation. B) Mutation profile of average mutation percentage of each of the 96 mutation categories from 7 adults (35 crypts) with no treatment is presented with standard deviation. C) The fold difference is calculated by dividing the percentage of mutation in each category from A) to that from B). The highest fold difference is highlighted in pink and the lowest fold difference is highlighted in orange.

**Figure 5.**
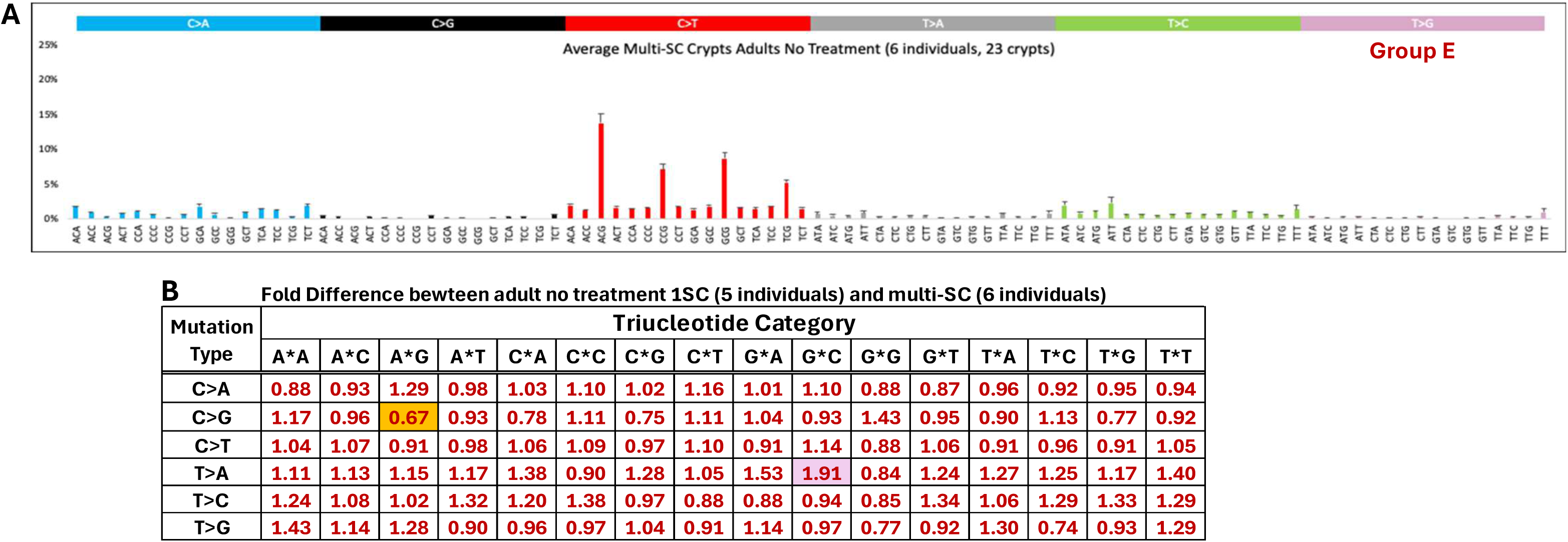
Mutation profile comparison between 1SC colon crypts and multi-SC colon crypts from adults with no treatment. A) Mutation profile of average mutation percentage of each of the 96 mutation categories from 6 adults with no treatment (23 multi-SC crypts) is presented with standard deviation. B) The fold difference is calculated by dividing the percentage of mutation in each category from Fig. 4A to that from Fig.5A. The highest fold difference is highlighted in pink and the lowest fold difference is highlighted in orange.

### Mutation profile of young children shows very limited difference from adults with no treatment

To further examine whether mutation profile of very young children is different from the mutation profile of adults, those profiles are compared. First, the average of 25 crypts from five very young children of ages 0.8 to 4 years old (Group F) is compared with the average of 35 crypts from seven adults with no treatment (Group D) (Fig. 6A and Fig. 4B). A total of 11 mutation categories showed >2-fold difference between the two groups (Fig. 6C).

**Figure 6.**
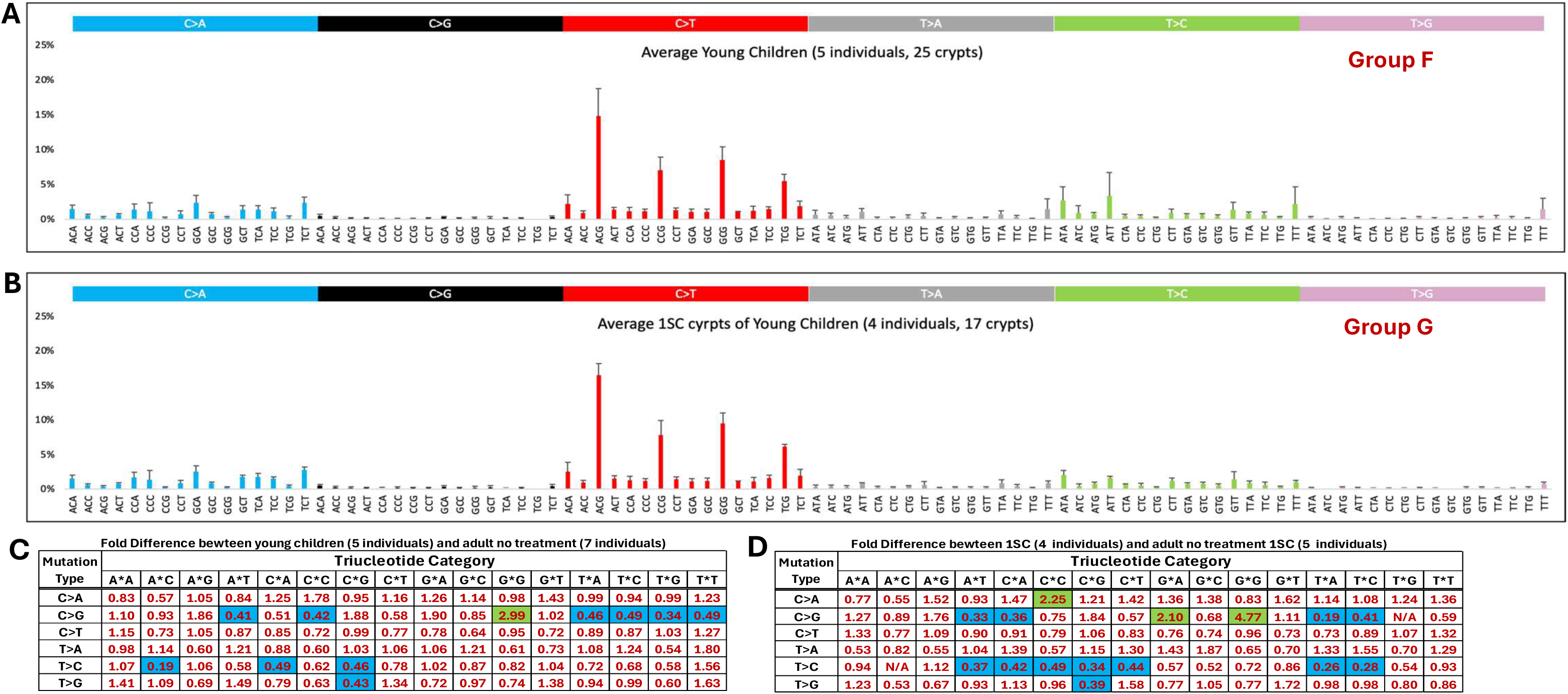
Mutation profile comparison between young children and adults with no treatment. A) Mutation profile of average mutation percentage of each of the 96 mutation categories from 5 young children (25 crypts) is presented with standard deviation. B) Mutation profile of average mutation percentage of each of the 96 mutation categories from 1SC crypts of 4 young children (17 crypts) is presented with standard deviation. C) The fold difference is calculated by dividing the percentage of mutation in each category from Fig. 6A to that from Fig. 4B. D) The fold difference is calculated by dividing the percentage of mutation in each category from Fig. 6B to that from Fig. 4A. In both C) and D), >2-fold difference is highlighted in green and <0.5-fold difference is highlighted in blue.

Second, the average of 17 1SC lineage crypts from four very young children of 0.8 to 2.1 years old (Group G; the 4 years old individual does not have any 1SC lineage crypt) is compared with the average of 12 1SC lineage crypts from five adults with no treatment (Group C) (Fig. 6B and Fig. 4A). A total of 15 mutation categories showed >2-fold difference between the two (Fig. 6D). The very young children showed a 2.99-fold and a 4.77-fold increase from adults with no treatment in C>G change of the GCG category in both the first and the second comparison, respectively. Also, the young children showed a 5.26-fold lower percentage (0.19-fold of adults) of T>C change in the ATC category compared with adults with no treatment in the first comparison. There is no SNV with a T>C change in the ATC category in any of the 1SC lineage crypts from young children (N/A is entered in Fig. 6D). It is important to note that all the >2-fold differences between the young children and adults with no treatment are all in the categories with very low percentage distribution in both groups. It is more likely that the large fold difference is due to fluctuations in the very small numbers observed. Therefore, the comparisons here suggest that the mutation profile is consistent throughout life under normal conditions.

### Chemotherapy and chemo/radiation combination treatment are associated with changes in specific mutation categories

Total mutation count analysis shows a clear increase in somatic mutations in individuals treated with chemotherapy or a chemo/radiation combination treatment, as presented above (Fig. 1A and 1B). To determine whether the increased mutation count is mutation category-specific versus non-specific, a thorough mutation profile comparison is needed. When the mutation profile of adults with treatment (Group H; Fig. 7A) and that of adults with no treatment (Group D; Fig. 4B) are compared, 11 mutation categories with >2-fold difference were identified (Fig. 7C, green highlights). Similarly, when the mutation profile of 1SC crypts from adults with treatment (Group I; Fig. 7B) and that of 1SC crypts from adults with no treatment (Fig. 4A) are compared, 16 of the 96 mutation categories showed >2-fold difference (Fig. 7D, green highlights). Furthermore, clonality of the crypts in individuals with treatment also showed no >2-fold difference clearly beyond potential experimental fluctuation (Fig. 7E). There are 10 mutation profile changes with chemotherapy or chemo/radiation treatment, if considering the overlaps of categories with >2-fold increase after treatment in all crypts and 1SC crypts (Fig. 7B, marked by gold and red arrowheads). It is noteworthy that the percentage of C>T changes at NCG sites (which have the highest mutation rates among the 96 mutation categories) in adults with treatment is consistently about 60% (0.62- to 0.65-fold) of that in adults with no treatment (Fig. 7C and 7D). Taken together, these findings show a clear change in the mutation profile with treatment, including higher relative incidence of the 10 mutation categories and lower relative incidence of the four C>T changes at NCG sites.

**Figure 7.**
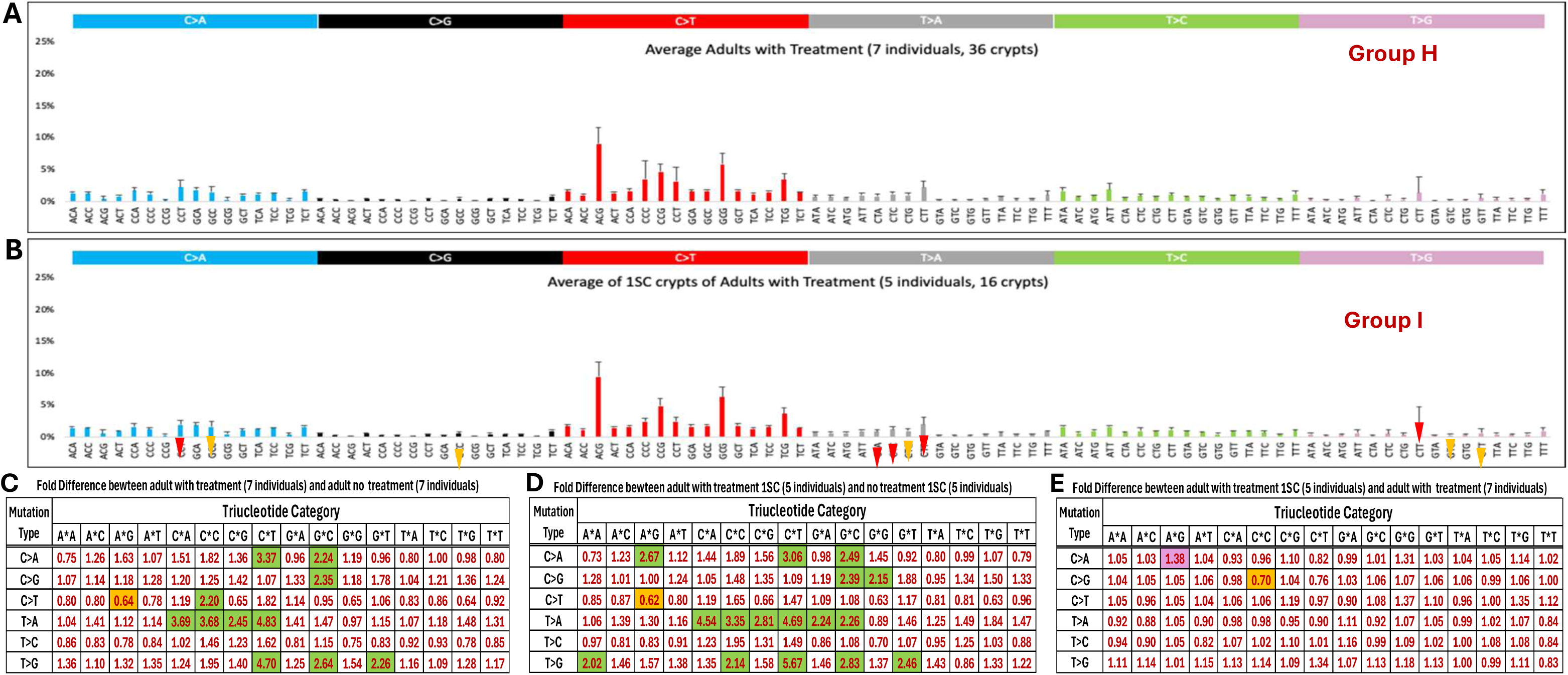
Mutation profile comparison between adults with treatment and no treatment. A) Mutation profile of average mutation percentage of each of the 96 mutation categories from 7 adults (36 crypts) with chemotherapy or chemo/radiation treatment is presented with standard deviation. B) Mutation profile of average mutation percentage of each of the 96 mutation categories from 1SC crypts of 5 adults (16 crypts) with chemotherapy or chemo/radiation treatment is presented with standard deviation. The gold arrowhead indicates >2-fold change and red arrowhead indicates >3-fold difference observed in both comparisons C and D. C) The fold difference is calculated by dividing the percentage of mutation in each category from Fig. 7A to that from Fig. 4B. D) The fold difference is calculated by dividing the percentage of mutation in each category from Fig. 7B to that from Fig. 4A. In both C) and D), >2-fold difference is highlighted in green and the lowest fold difference is highlighted in orange. E) The fold difference is calculated by dividing the percentage of mutation in each category from Fig. 7B to that from Fig. 7A. The highest fold difference in E) is highlighted in pink and the lowest fold difference is highlighted in orange.

The combined increase in mutations across the other 92 categories could reduce the relative presence of C>T changes at NCG sites in individuals receiving treatment, since chemotherapy and radiation are known to cause DNA damage. This possibility can be evaluated by comparing the observed mutation count with the expected age-appropriate mutation count derived from individuals with no treatment. As described in Materials and Methods, the mutation count difference between treated 1SC crypts and the expected normal count was calculated by subtracting the age-appropriate expected count from the observed count. When averaged across treated 1SC crypts, six mutation categories showed lower-than-age-appropriate mutation counts, ranging from 2 to 45 mutations below the expected count (Fig. 8; Supplementary Table S3, pink highlight in the top table). The four C>T changes at NCG sites showed the largest decreases relative to their expected counts among all mutation categories. It is important to note that the mutation count should not decrease beyond normal biological fluctuations in any category, because somatic mutations are not expected to disappear from a surviving lineage except through rare back mutation or loss of the mutated lineage. The non-trivial decreases from expected mutation counts in C>T mutations at NCG sites are observed in nearly all crypts from individuals with treatment (Figure 8 and Supplementary Table S3). This pattern is consistent with the possibility that treatment selects against stem-cell lineages with higher pre-existing age-associated mutation burden.

**Figure 8.**
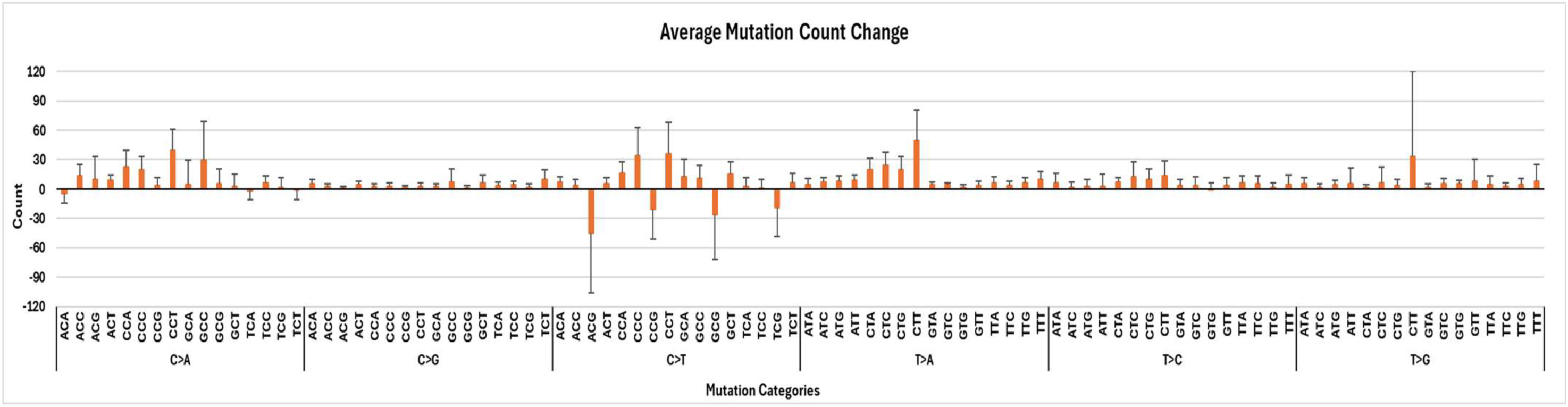
Average difference between observed mutation count and expected age-appropriate mutation count in 1SC lineage crypts from individuals with treatment. The graph shows the average observed minus expected mutation count for each mutation category across 1SC lineage crypts from five individuals with treatment. The numerical summary is presented in Supplementary Table S3, and the mutation count for each mutation category of all crypts can be found in Supplementary Table S2.

## DISCUSSION

Somatic mutation occurs in cells in a manner that is both dependent and independent of DNA replication. Each new mutation continues to exist in all the descendant cells. Therefore, two daughter cells from the same progenitor would harbor all the mutations present in the progenitor cell while acquiring different new mutations as they become two independent cells and thereafter. This cycle of sharing the old and acquiring the new continues as cells divide and remain in the human body. These somatic mutations cause increasing heterogeneity in the human body with age. The somatic mutations accumulated in a clonal cell population derived recently from a progenitor, such as tissue specific stem cells, can be obtained by whole genome sequencing (WGS) of the clonal cell population.

As mentioned earlier, each colon crypt has been conventionally considered a single stem cell lineage monoclone, based on several elegant studies that examined one or very few genetic markers. Based on this prior inference, we generated a dataset by sequencing five or six single colon crypts and a bulk DNA control from each of the 21 individuals of different ages using our novel WGS method (12, 14). We analyzed the autosomal SNVs from this dataset both quantitatively and qualitatively to determine the rate of mutation increase, which is a representation of aging, and the pattern of somatic mutation in human colon crypt stem cells. However, our previous variant allele frequency distribution analysis found that human colon crypts are as often polyclonal as monoclonal (12). In this aging rate analysis, we thoroughly analyzed the autosomal SNVs from this dataset, both quantitatively and qualitatively, with crypt clonality in mind.

Our findings in this study include the following: 1) somatic mutation count increases with age in colon crypt stem cells; 2) we determined an accurate rate of mutation per year for each of the 96 trinucleotide mutation categories in normal 1SC lineage (monoclonal) colon crypts; 3) clonality of a colon crypt influences somatic mutation count (quantitatively) in the crypt but does not qualitatively change the mutation profile of the crypt; 4) mutation profile does not change with age under normal conditions; 5) chemotherapy and chemo/radiation treatment can impact specific mutation categories and change mutation profile.

It is important to note that only 49% of the colon crypts sequenced are monoclonal and do not allow adequate statistical power for the analysis. However, this is the only study in which multiple primary cell clones from each individual in a cohort of 21 individuals age 10 months to 90 years old are sequenced by WGS to high depth with matched bulk DNA control for each individual. The mutation count can only be accurately determined when a cell population is a monoclone. Furthermore, we took stringent filtering steps and removed high false positive variants with tools we developed for a high-quality analysis. In addition, there is no known germline pathological mutation among the 160 DNA damage response and repair loci examined in these individuals, as reported in an independent analysis (20). This study provides the first estimate of mutation accumulation with age in monoclonal normal human colon crypts, with reasonably high confidence, although further confirmation is needed.

The estimated total mutation rate is 44 per year based on the average mutation count of 1SC lineage crypts from each of the 11 individuals with no chemotherapy or chemo/radiation treatment (Fig. 1B). The estimated total mutation rate is 41 per year when each 1SC lineage crypt is considered in the analysis (Supplementary Fig. S1). The baseline mutation count at age 0 is 179 when extrapolated from the individual average analysis (Fig. 1B), and it is 225 when extrapolated from each single crypt analysis (Supplementary Fig. S1). This discrepancy is most likely caused by the small sample size. A previous study estimated a mutation rate of ∼40 mutations per year (an interval between 26.9 and 50.6) in human colon by sequencing 21 clonal organoid cultures from six individuals younger than 25 or between 50 and 70 years old (13). Another study estimated a rate of 43.6 mutations per year in a low depth WGS dataset of colon crypts without matched controls (11). While our estimate and previous estimates of the mutation rate are within a relatively narrow range, the small difference per year and the initial baseline mutation at birth can lead to a notable difference in older individuals. Difference in the extrapolated baseline mutation count at birth (179 vs. 225) can account for an appreciable bias in the very young individuals since the average mutation count in each clonal cell population for children less than four years old is less than 500 (Fig. 1). Therefore, analyzing somatic mutations from a larger sample of 1SC lineage colon crypts should be considered before the age dependent mutation count can be used as a standard for studying conditions or factors that cause deviations from normal.

We also examine the mutation rate for each mutation category and evaluate whether it increases linearly over the human lifespan. We find only a single mutation category, T>G change at CTT sites, did not meet nominal P<0.05 threshold for a non-zero regression slope, with p = 0.051 when 1SC lineage crypts from individuals with no treatment are considered (Fig. 2D). The mutation categories with the highest mutation rate are C>T changes at NCG sites (N is any nucleotide) in mutation count analysis of all four Groups (Fig. 2). These four mutation categories are clearly the drivers of mutation accumulation, as expected due to 5-methylcytosine deamination, at a rate of 5.8, 2.9, 3.8, and 2.2 per year at ACG, CCG, GCG, and TCG sites, respectively, in the 1SC lineage crypts from individuals with no treatment. Each of the other 92 mutation categories contributes much less to mutation accumulation individually than each of the NCG mutations, but they account for ∼60% of total mutations per year. Overall, more than half of the mutations involve C>T changes and C>G, T>A, and T>G changes are the least abundant (Table 1). Using our estimates of the specific mutation rate for each mutation category, an age-dependent estimate of mutation counts can be derived to evaluate deviations and measure the biological age for any study subject. For example, as expected, the colon crypts from individuals with treatment in our study showed higher mutation counts. It is possible that different tissues can have different mutation rates due to exposure to environmental or therapeutic factors. It would be interesting to compare our findings in 1SC lineage colon crypts to clonal cell populations from other tissue types in the future.

The mutation profile represents the relationship among the 96 mutation categories in a sample as mentioned above. Comparing mutation profiles across groups reveals that they do not change with age. This is an important finding that supports the validity of cross comparison of a sample with a standard normal mutation profile of the same tissue. Different tissues have been shown to have different mutation profiles (13), but establishing a comprehensive mutation count and mutation profile with a large number of clonal cell populations from different tissues is essential in establishing a normal standard for mutation studies. Furthermore, comparison between 1SC lineage crypts and multi-SC lineage crypts clearly indicates that clonality of the crypt does not affect its mutation profile. This finding demonstrates that the mutation profile does not vary appreciably in different cells from the same tissue. Therefore, the mutation profile of a non-clonal normal cell population can be considered representative of the specific tissue studied.

The mutation count analysis suggests that most of the individuals with chemotherapy and chemo/radiation treatment have a higher total number of mutations than the linear trendline indicates for the cohort (Fig. 1A). The mutation profile analysis shows a lower percentage of C>T changes at NCG sites than normal. Mutagenic therapeutic agents are expected to increase mutations. The C>T changes are mostly replication independent and more time dependent than other mutations. The change in the mutation profile in individuals with treatment may be due to more mutagen generated mutations in other mutation categories than at NCG sites. However, the number of C>T changes at NCG sites that already exist in the stem cells prior to treatment should at least remain unchanged. Yet, upon examination of the counts of C>T changes at NCG sites in individuals with treatment, a clear decrease is evident (Fig. 8; Supplementary Table S3). While there is an expected number of mutations in each individual based on age, the mutation count from each cell or cell clone is distributed within a range. It is possible that mutagens from chemotherapy and radiation are more lethal to cells harboring a higher number of mutations accumulated with age prior to treatment. Not all cells in the same tissue would have exactly the same number of mutations; and the cells with more mutations, which are biologically older, may not survive the treatment as well as the biologically younger cells. Therefore, the surviving stem cells after treatment show a young profile with fewer age dependent mutations such as C>T changes at NCG sites, suggesting the older cells may be selected against by the treatment. This interpretation is speculative but might be hypothesis-generating because the current study was not designed to investigate this hypothesis. It will require an appropriately designed larger study to verify and further explore. However, this comparison illustrates the importance of conducting both mutation count and mutation profile analyses to assess the impact of extrinsic factors on DNA mutation and the biological age of somatic cells. The mutation profile is not affected by clonality and can be established from polyclonal cell populations. It is not always possible to obtain clonal cell populations from primary human tissues for sequencing and directly interpretable mutation count analysis. However, an expected mutation count can be calculated based on the age of the subject when a normal standard for the specific tissue is available.

In conclusion, we have provided a mutation rate for each of the 96 mutation categories and a normal mutation profile for human colon epithelium. We also present a framework for removing potential false positive variants for a high-quality analysis. Although further study with adequate statistical power is needed to confirm our findings, what we presented here lays the foundation and provides guidance for future studies.

## ACKNOWLEDGEMENT

We would like to thank the Norris Comprehensive Cancer Center Translational Pathology Core for the sample collection. We would like to acknowledge the Norris Comprehensive Cancer Center Molecular Genomics Core and the Keck Genomics Platform at the University of Southern California for the sequencing work. This work is supported by funds from NIA R01 AG 067615 and the Catherine and Joseph Aresty Endowment to CLH. MRL is supported by NIGMS R35118009.

---

**Supplementary Figure S1.**
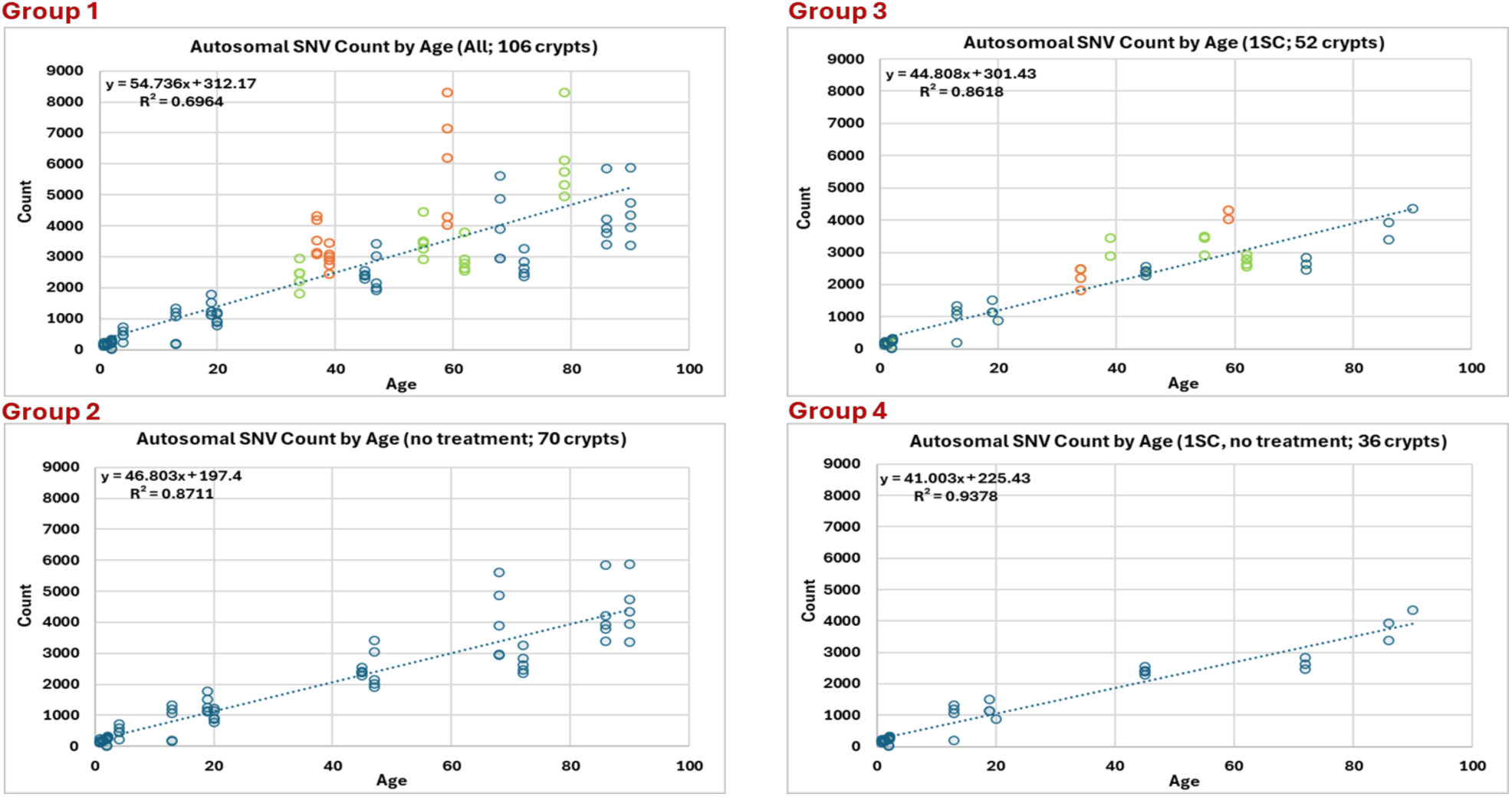
SNV count increase in human colon crypts with age. Autosomal SNV count of each colon crypt is plotted by age of the individual. The top left panel includes all 106 crypts from 21 individuals. The bottom left panel includes 70 crypts from 14 individuals with no treatment. The top right panel includes the 52 single stem cell lineage colon crypts from 16 individuals. The bottom right panel only includes 36 single stem cell lineage crypts from 11 individuals with no treatment. Each circle, filled with 90% transparency, represents the autosomal SNV count of a single crypt. The individuals with chemotherapy are marked in green and the individuals with chemo/radiation combination treatment are marked in orange.

**Supplementary Figure S2.**
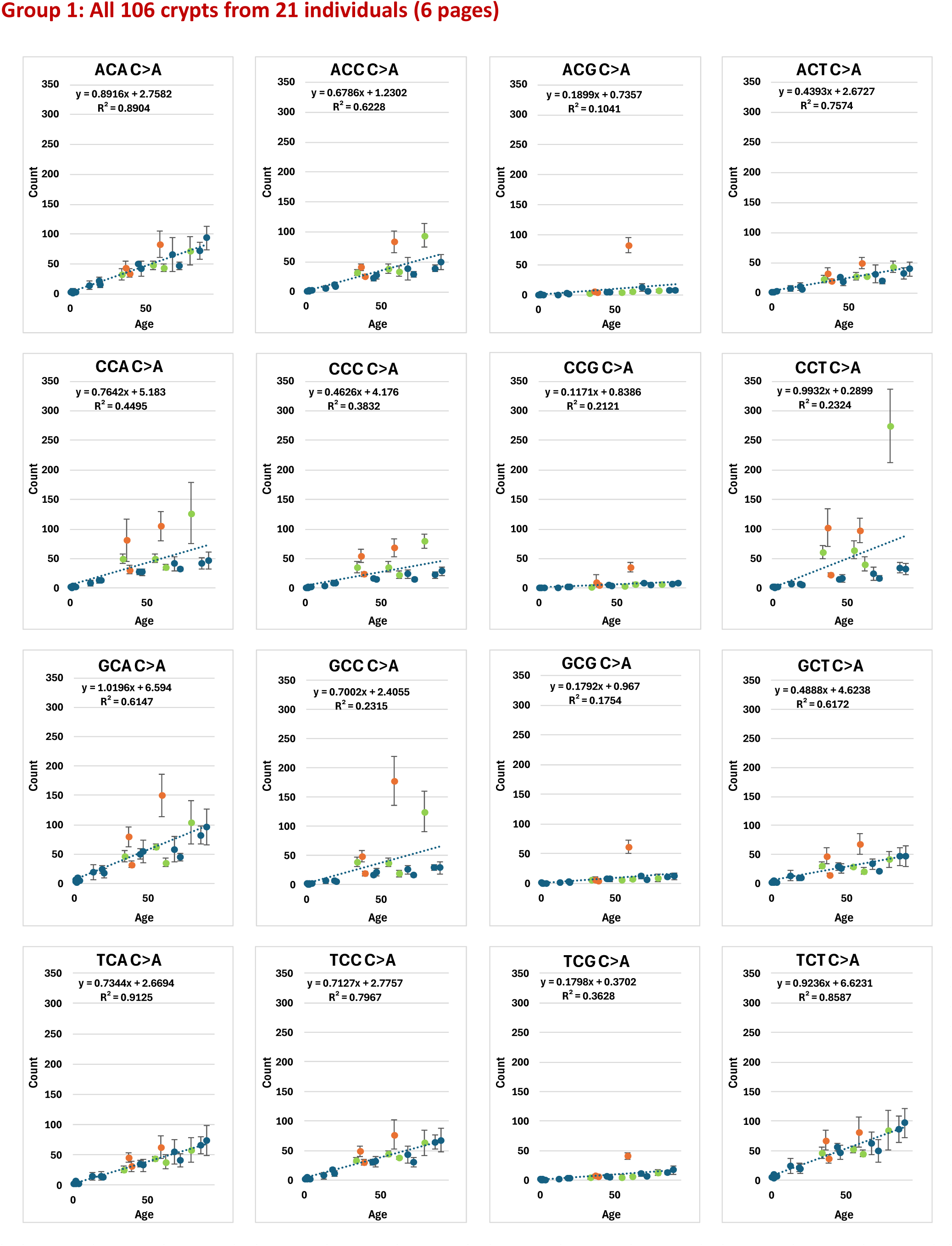
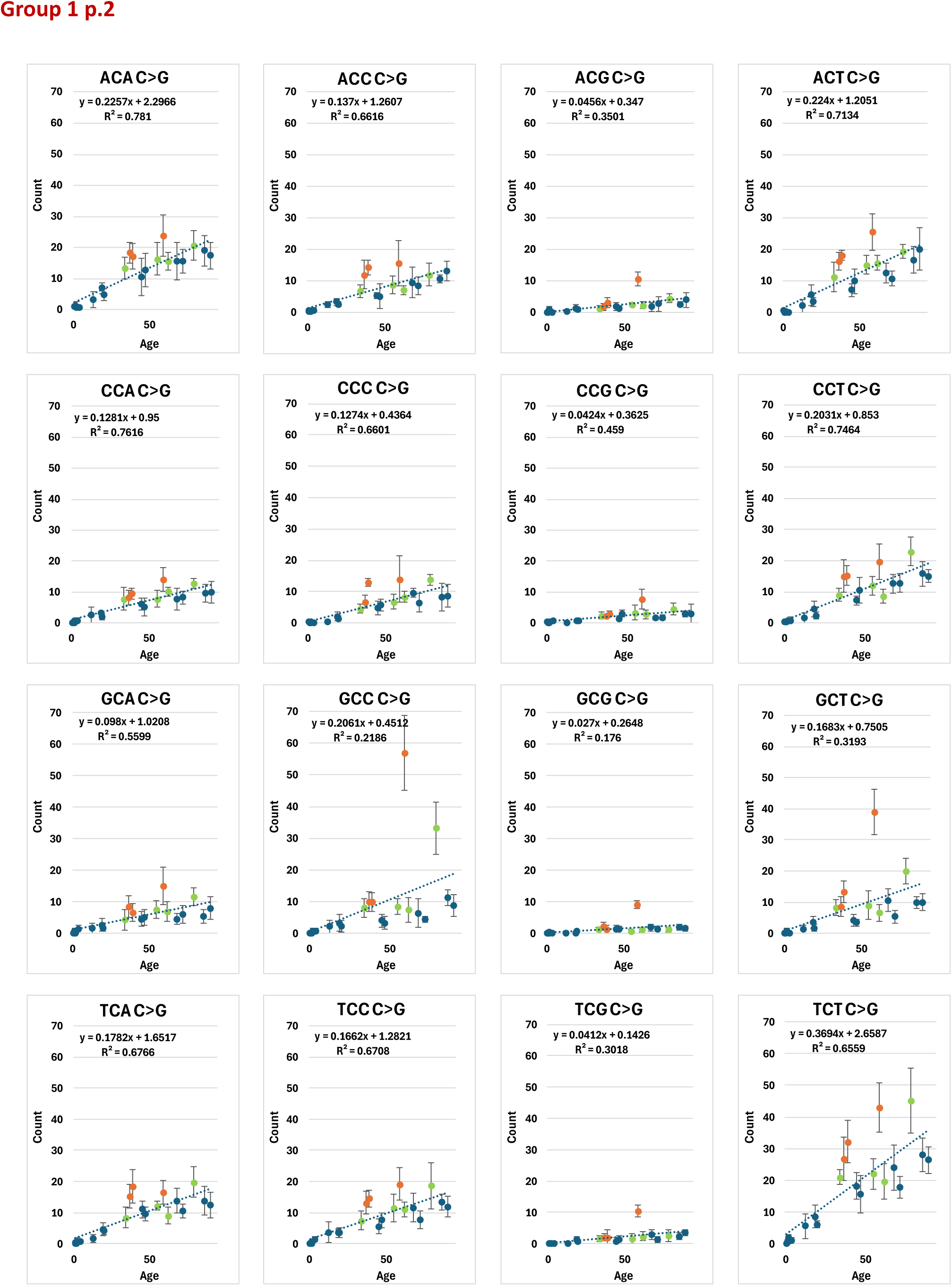
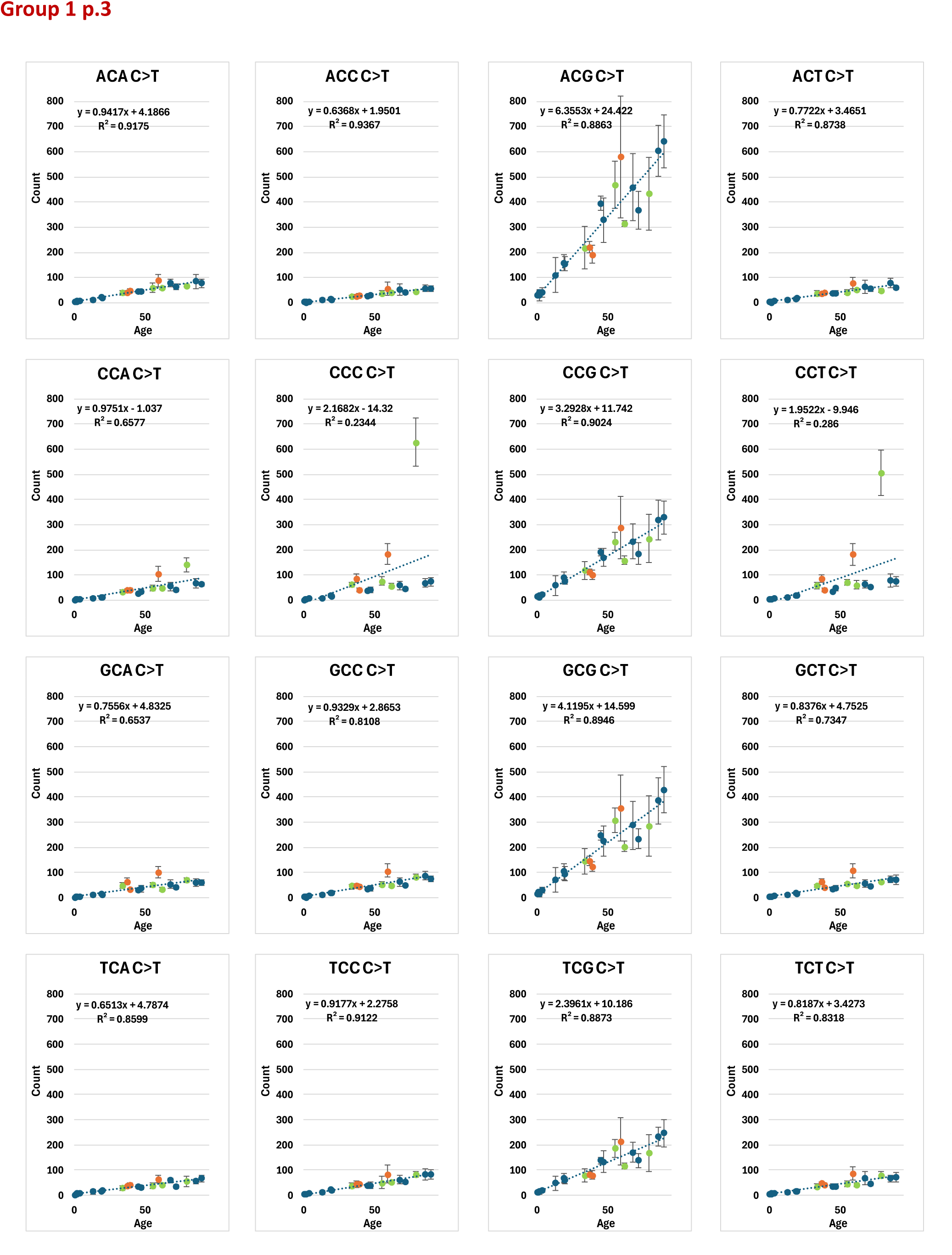
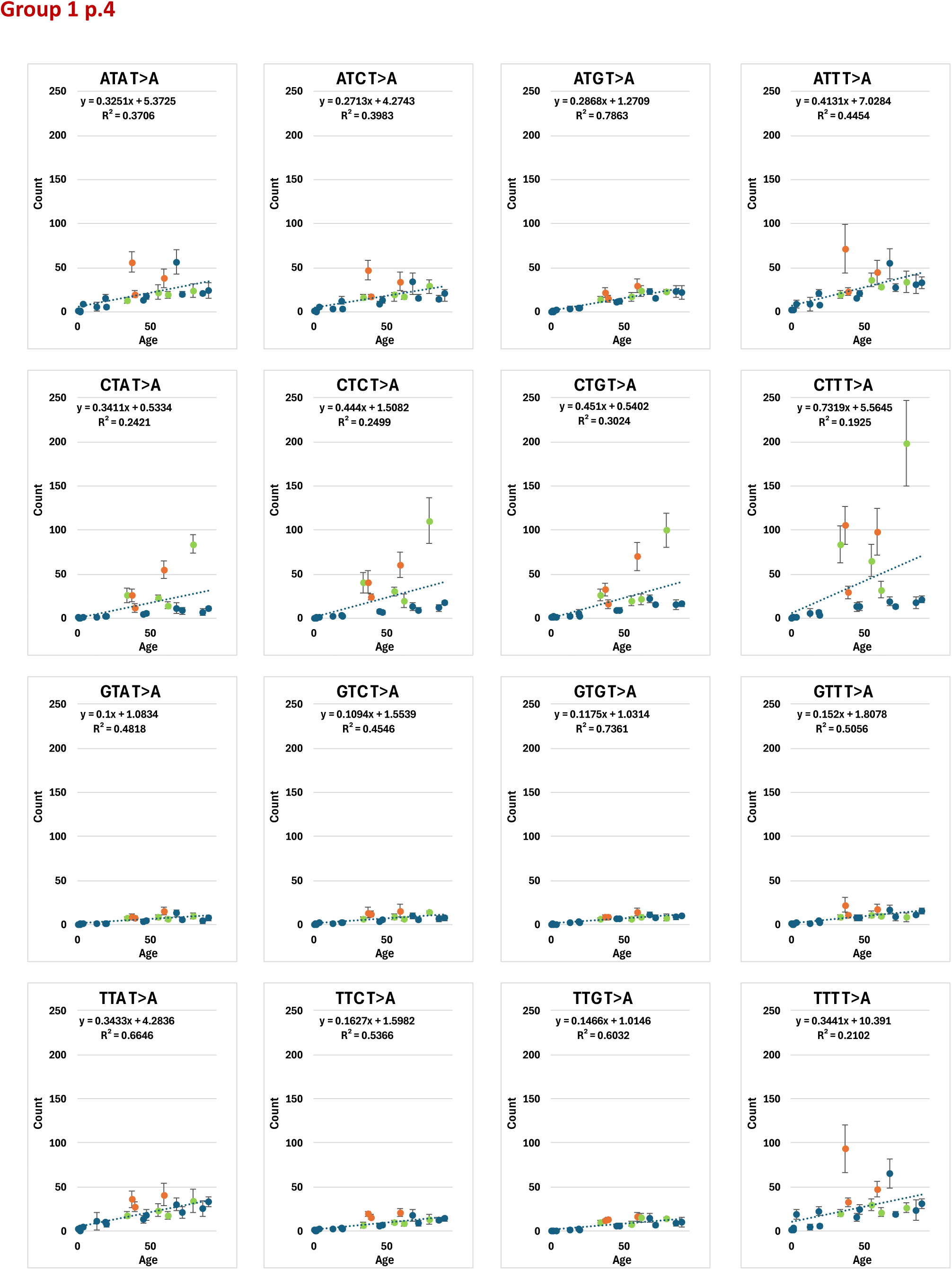
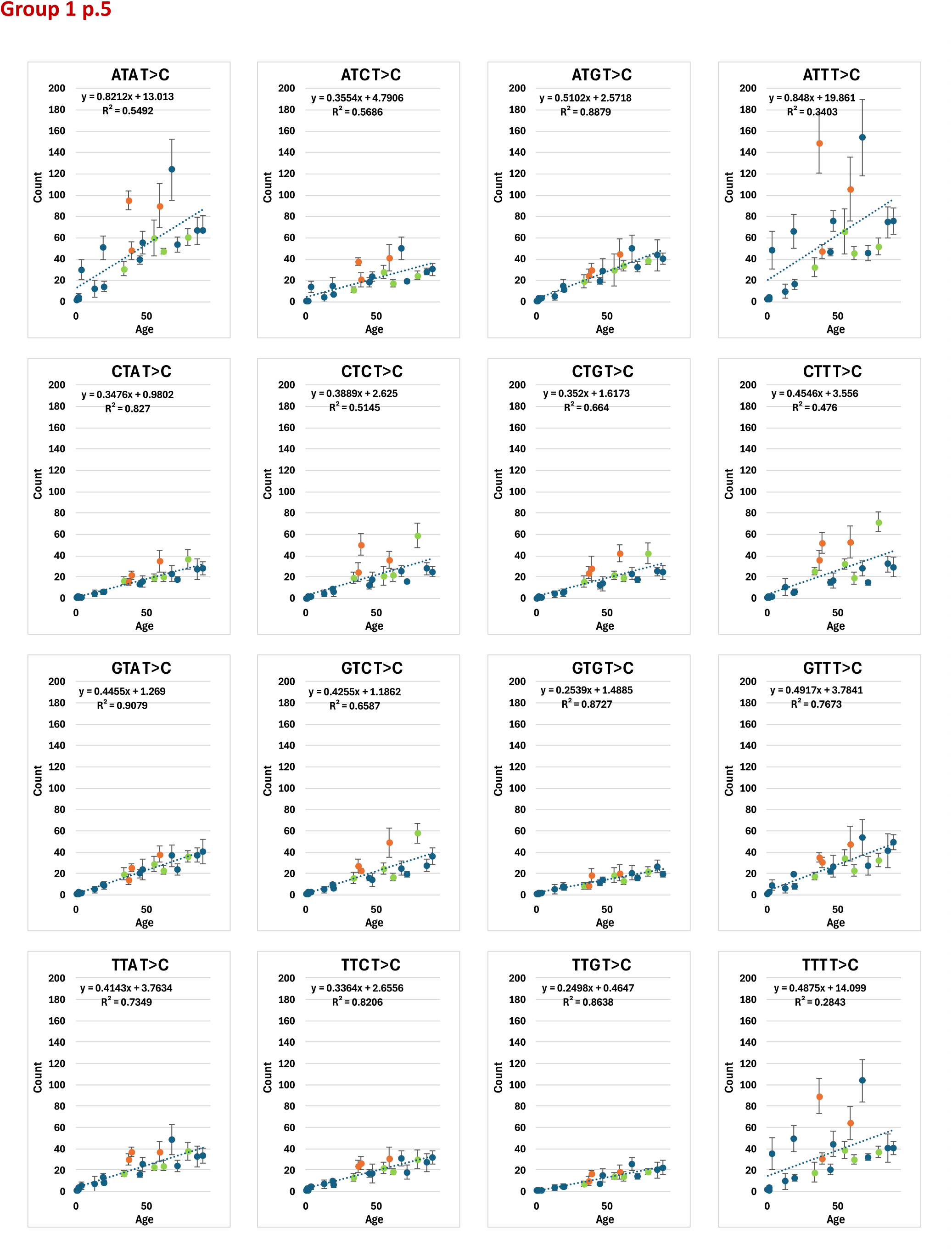
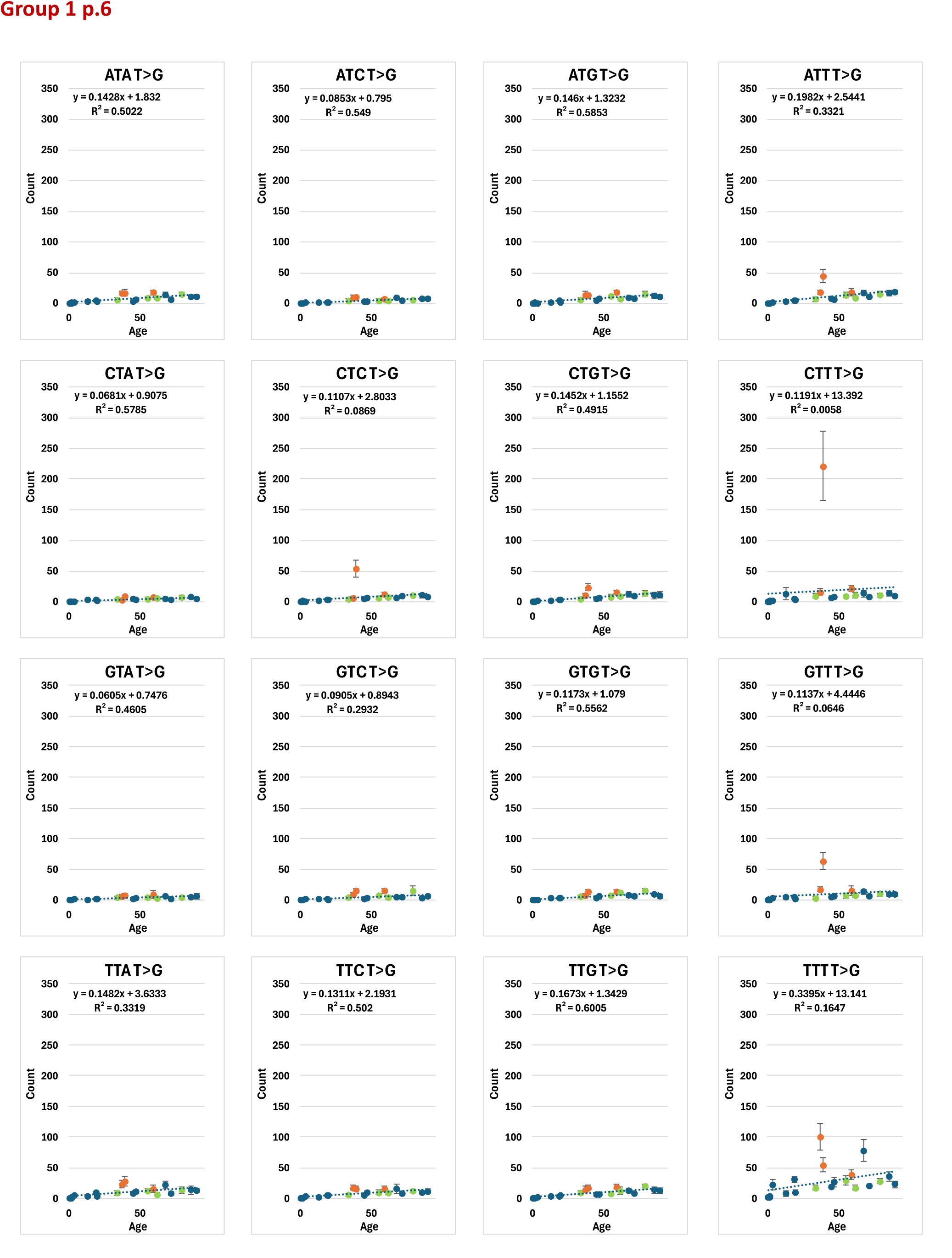
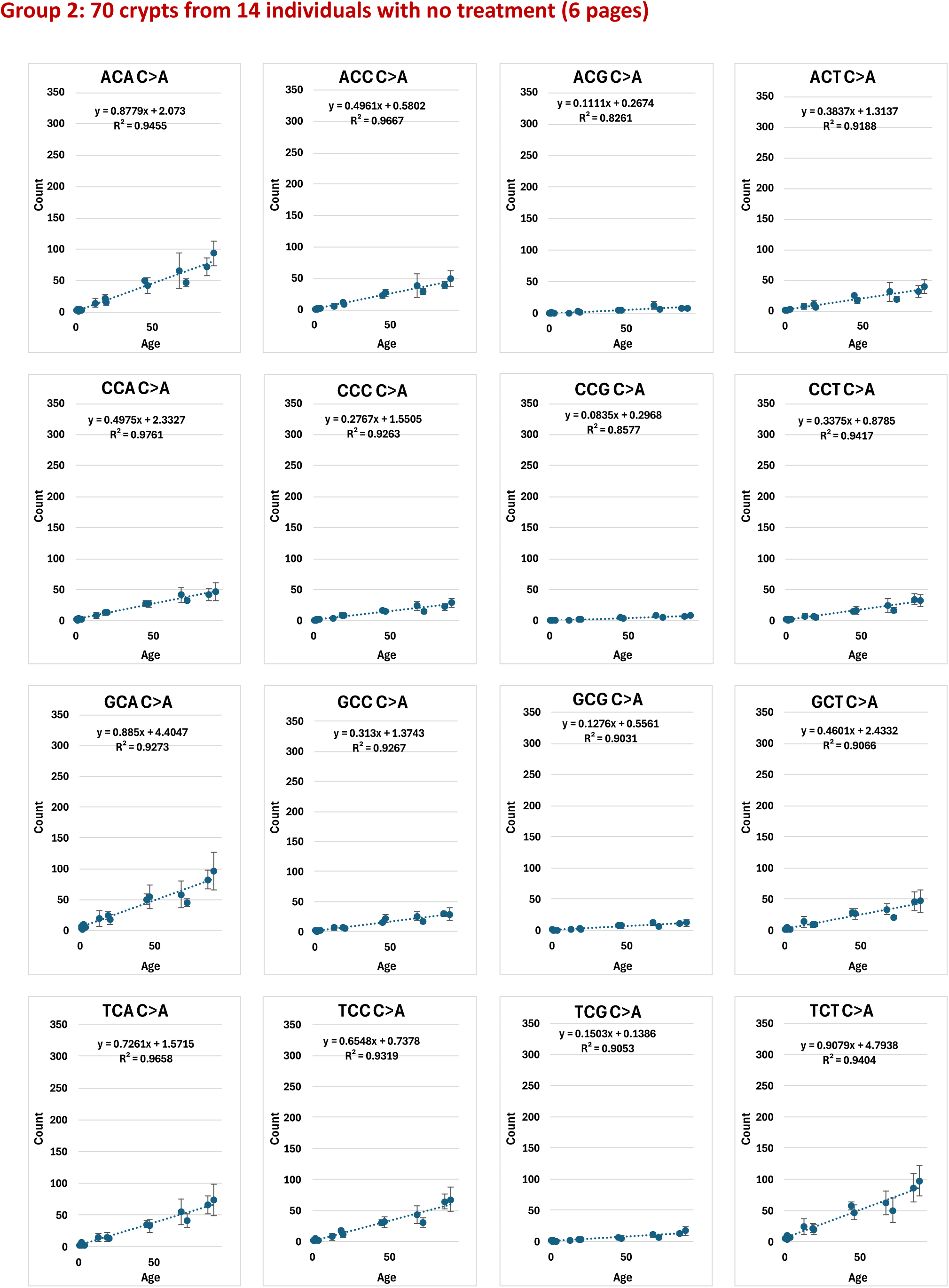
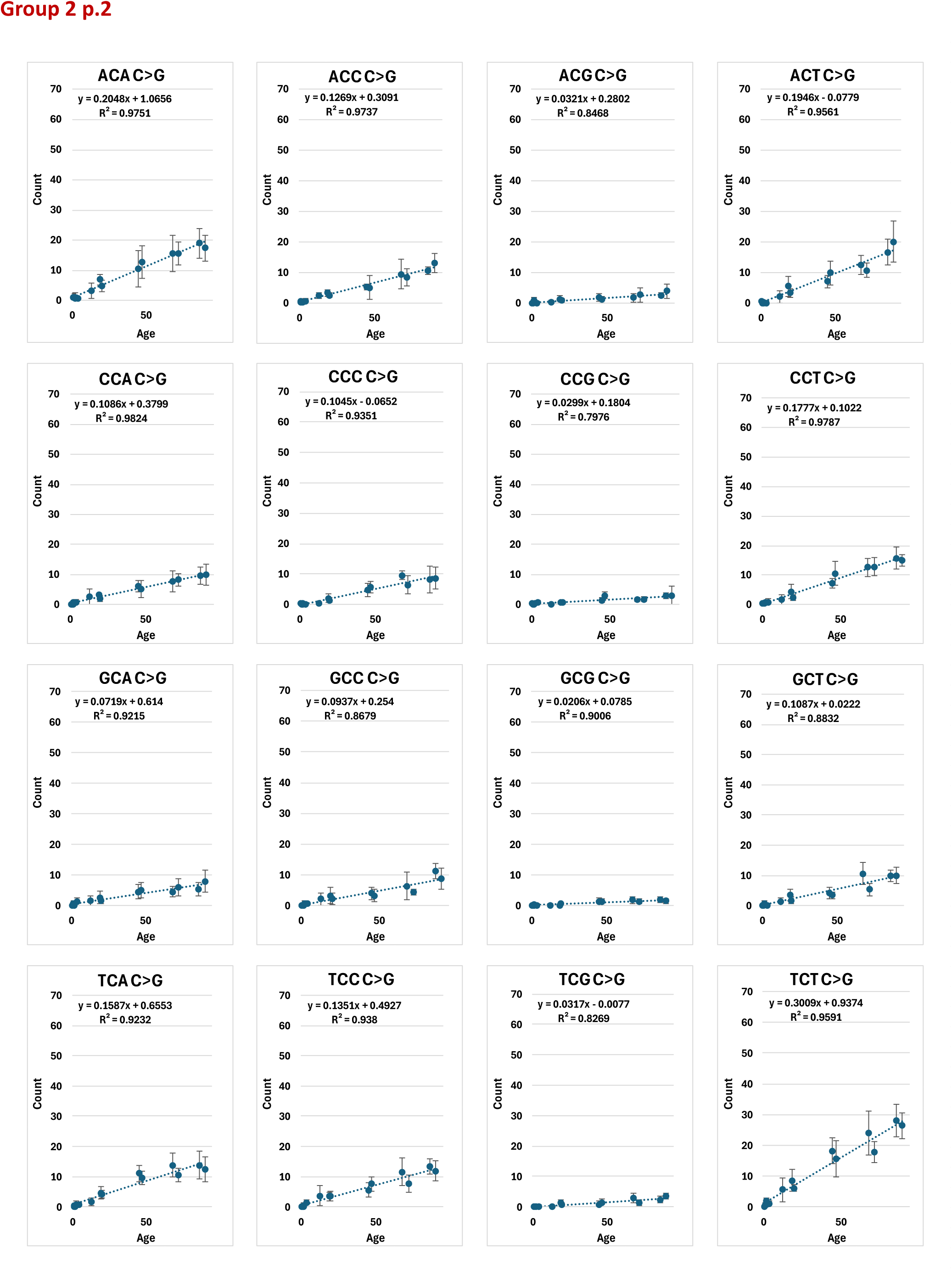
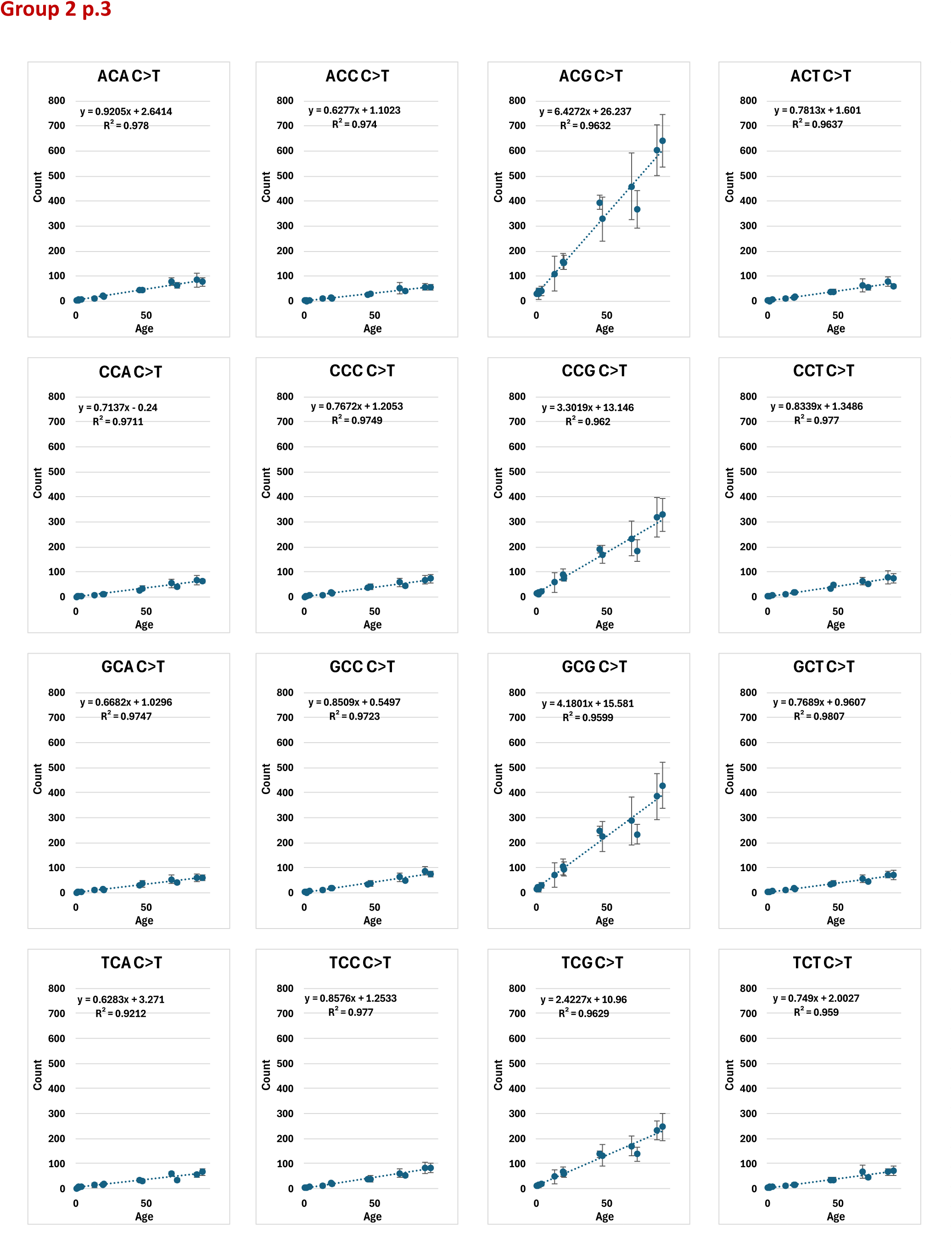
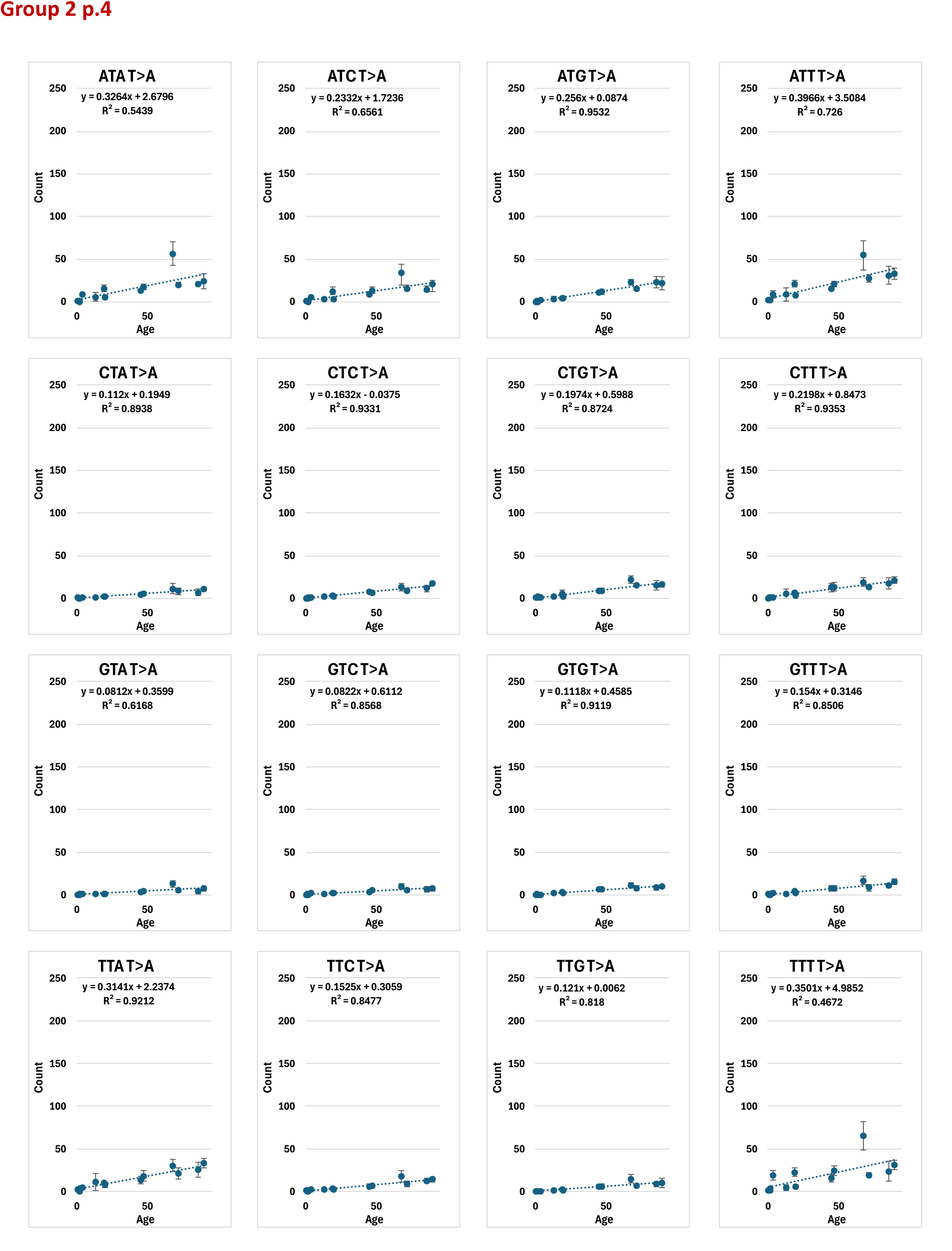
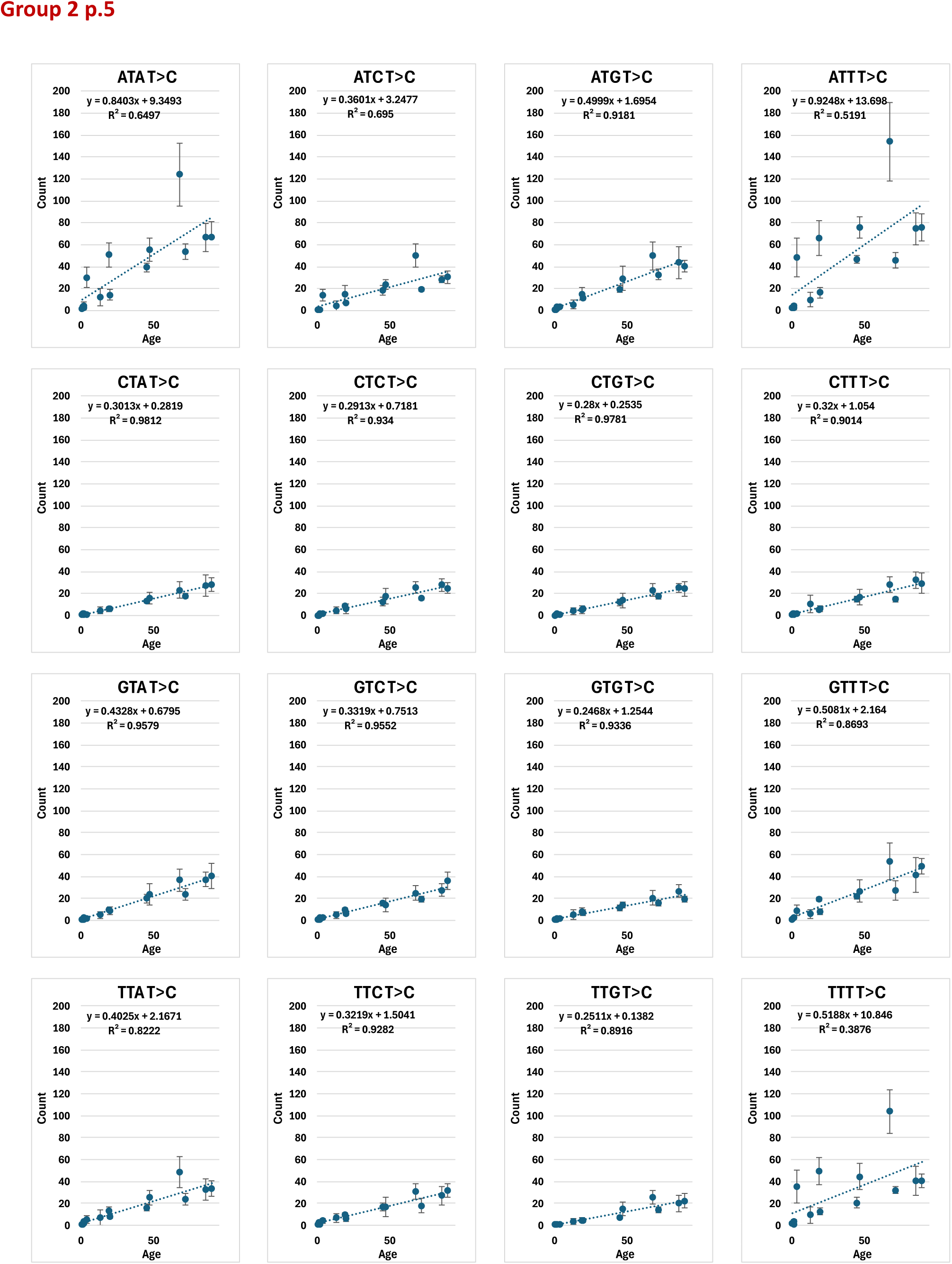
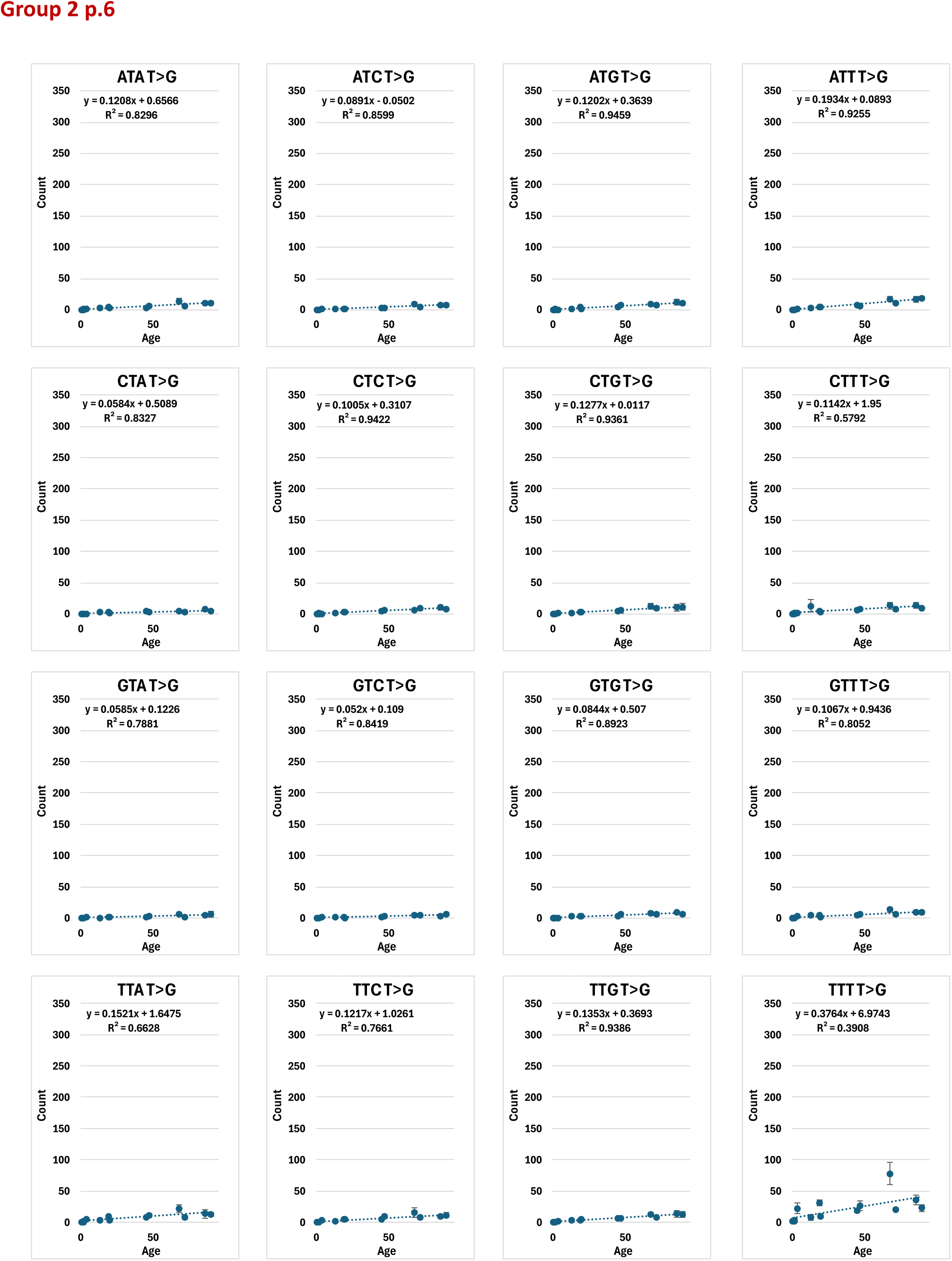
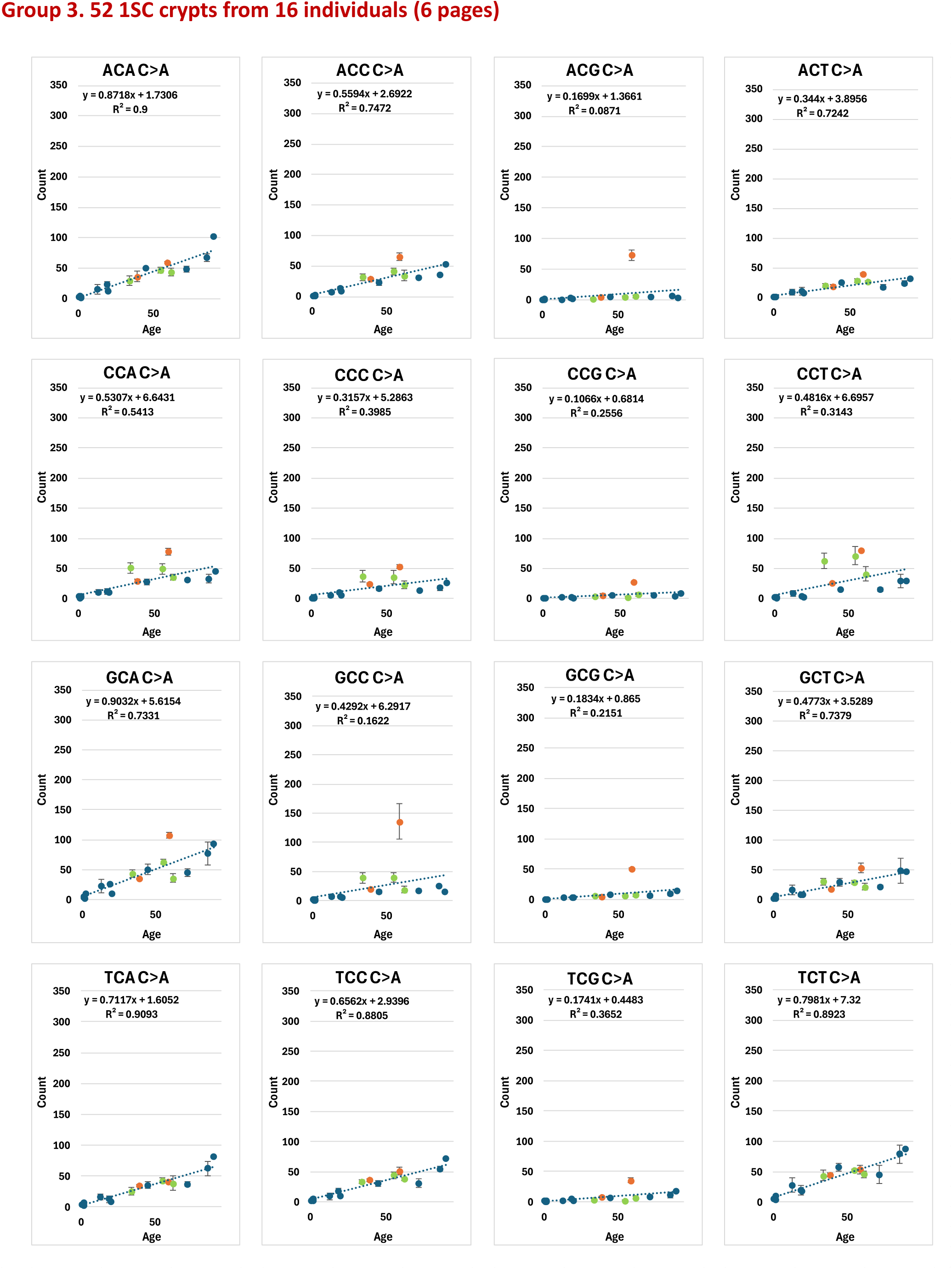
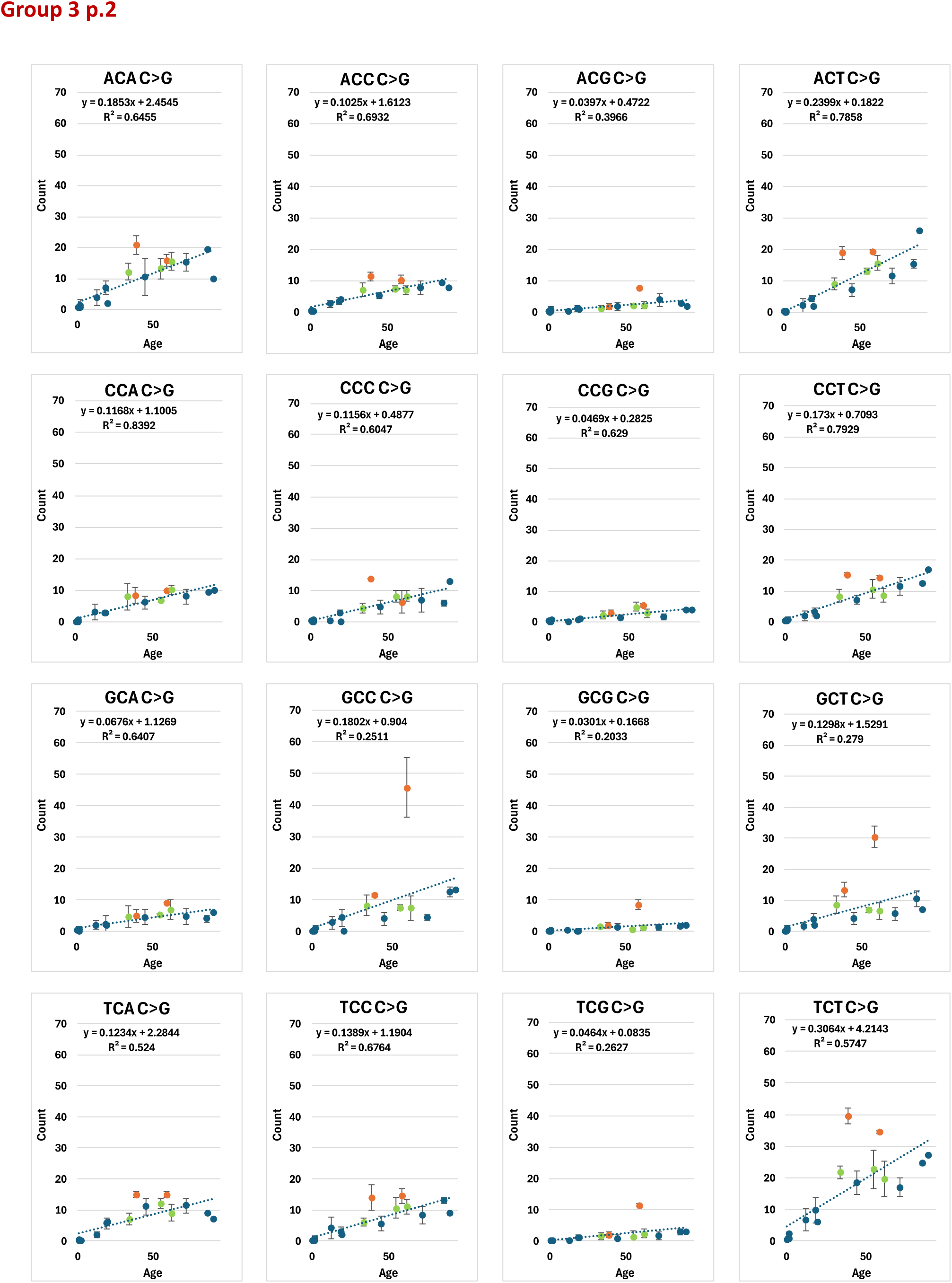
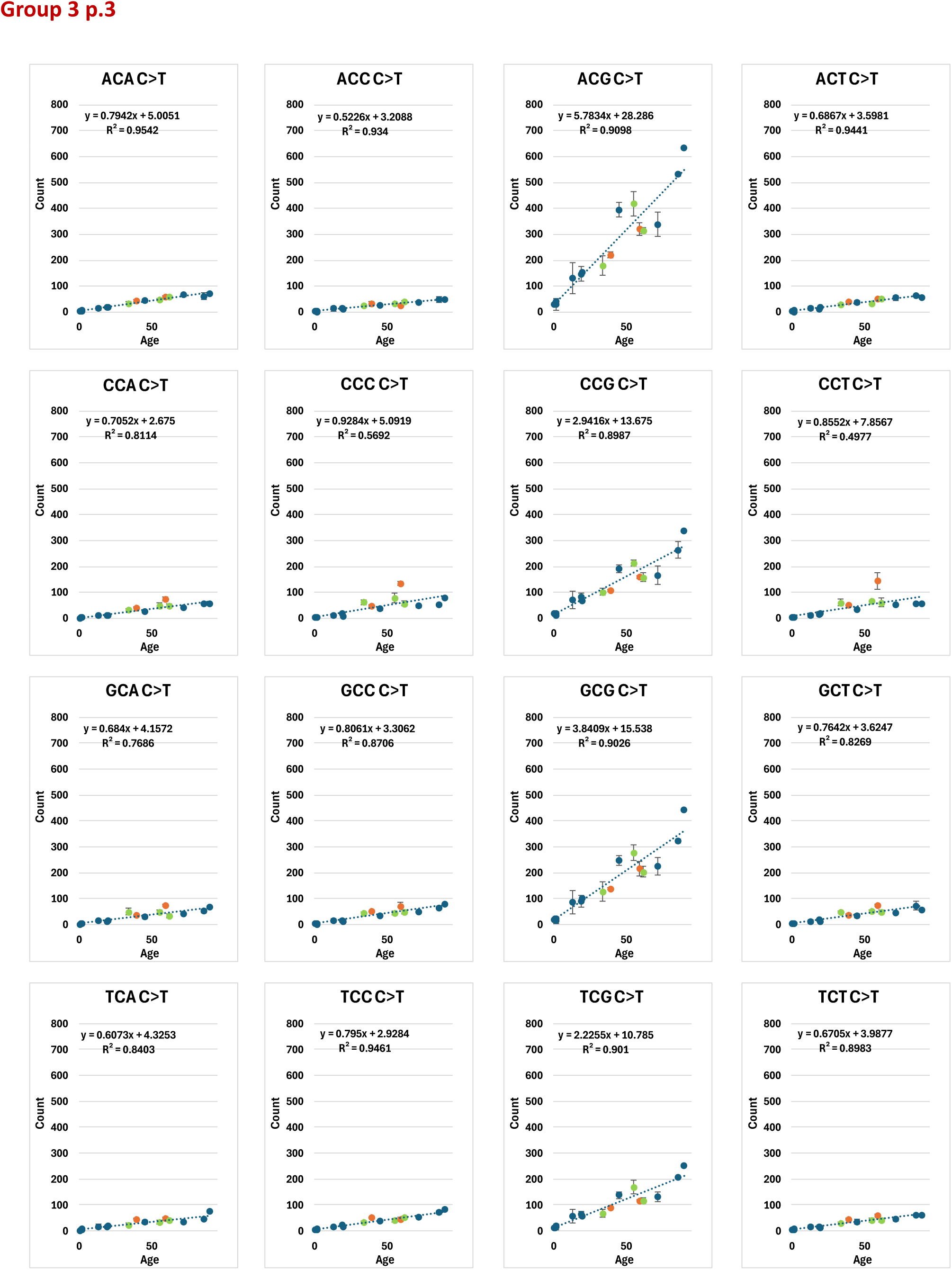
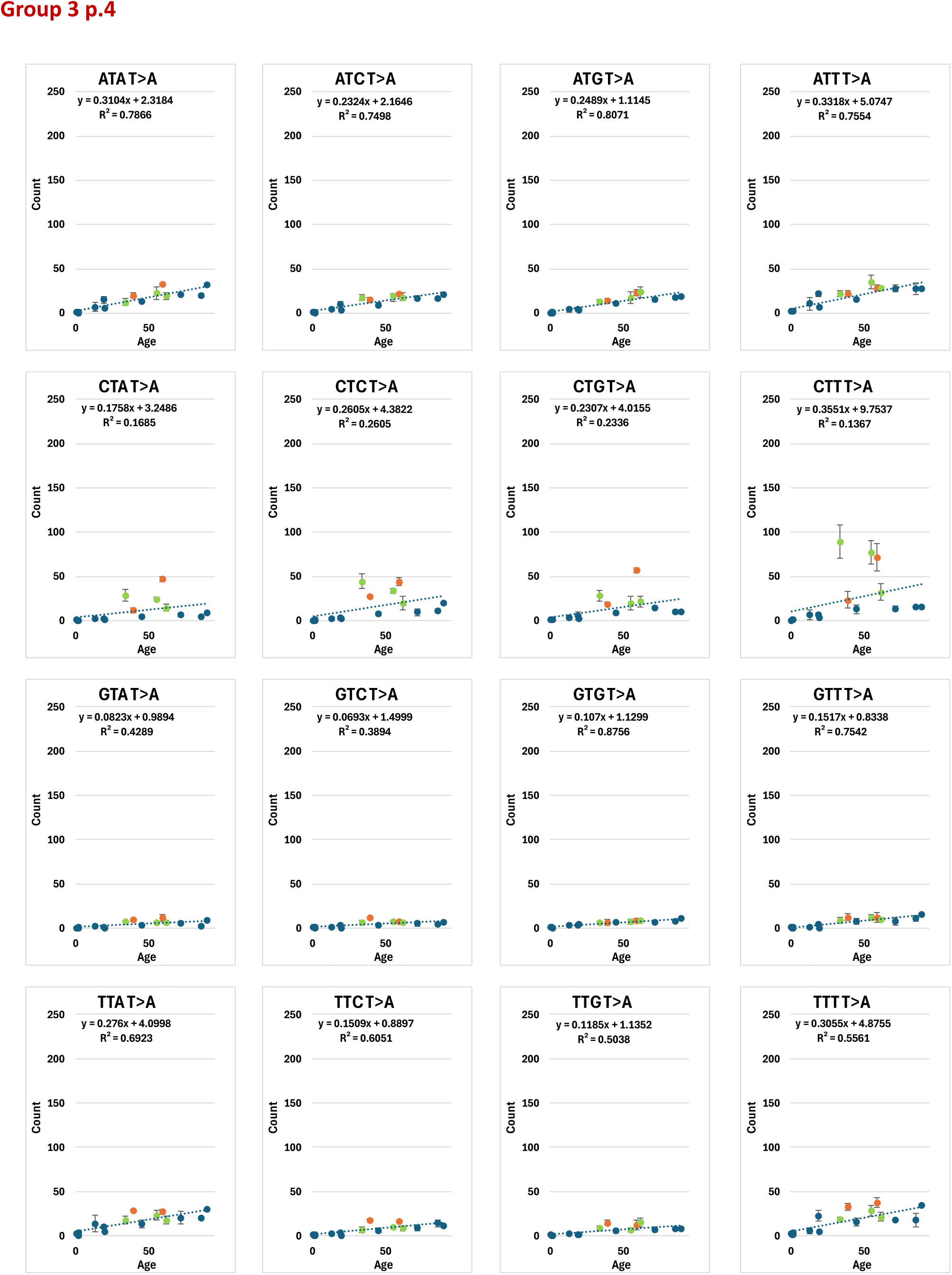
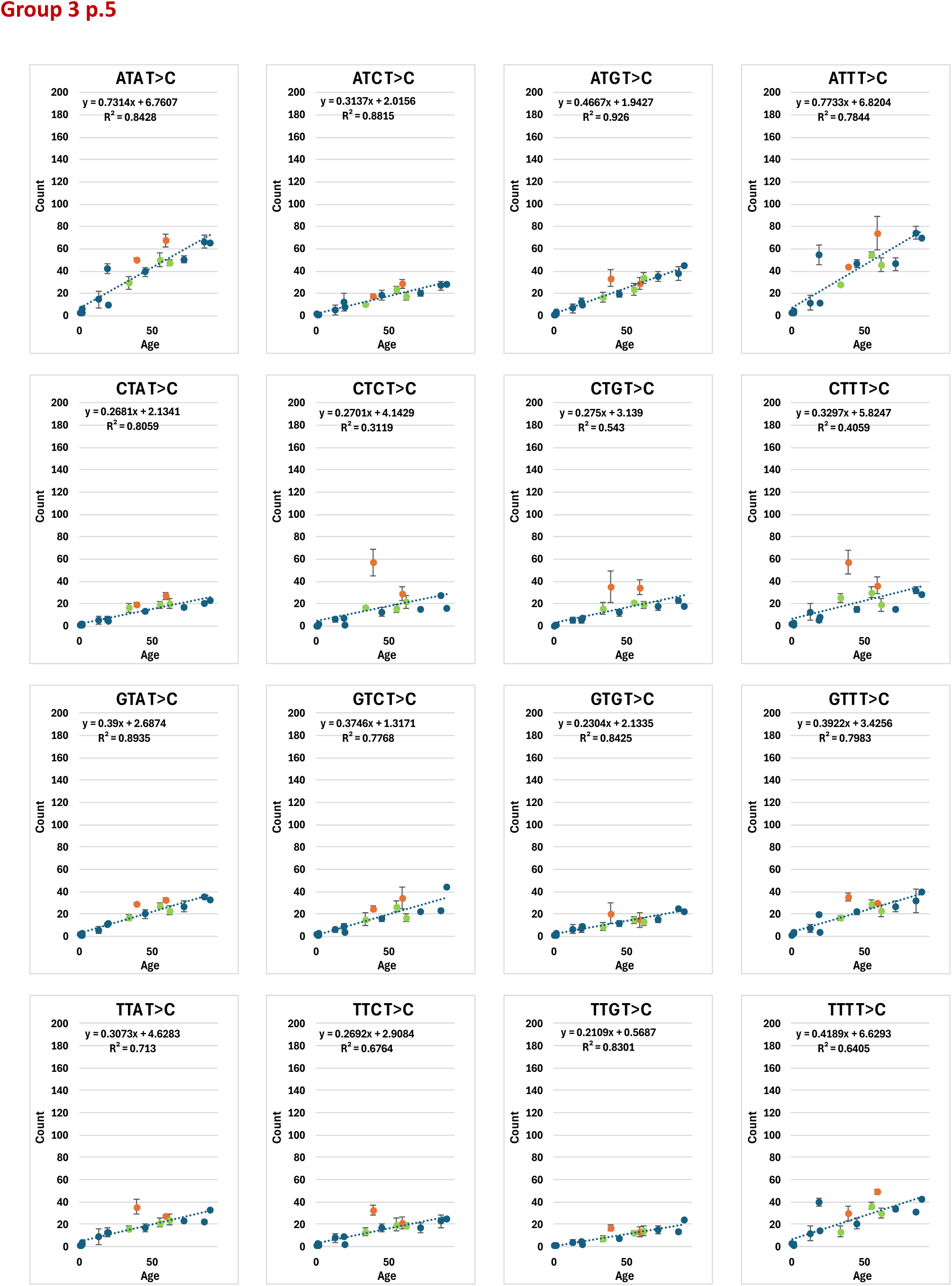
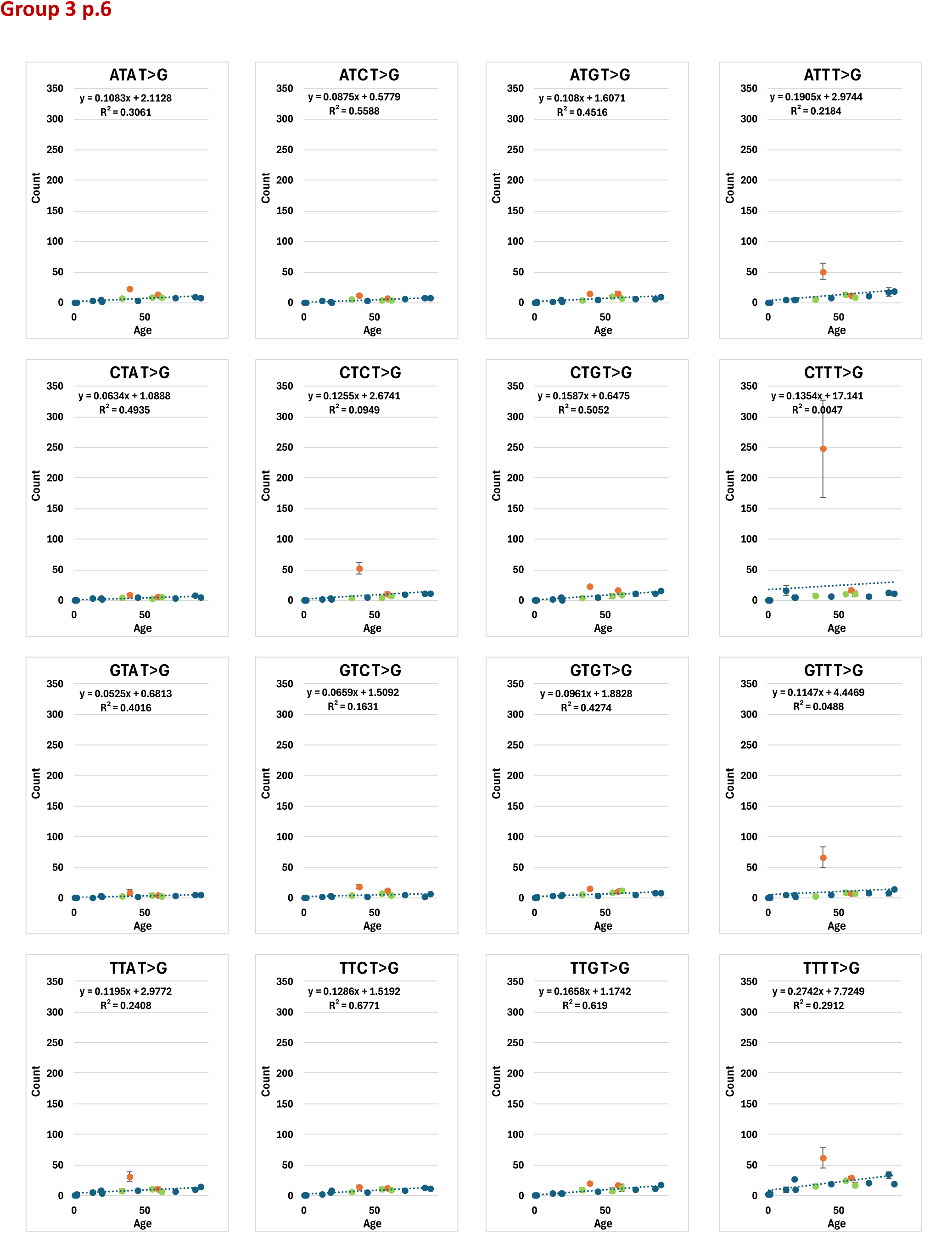
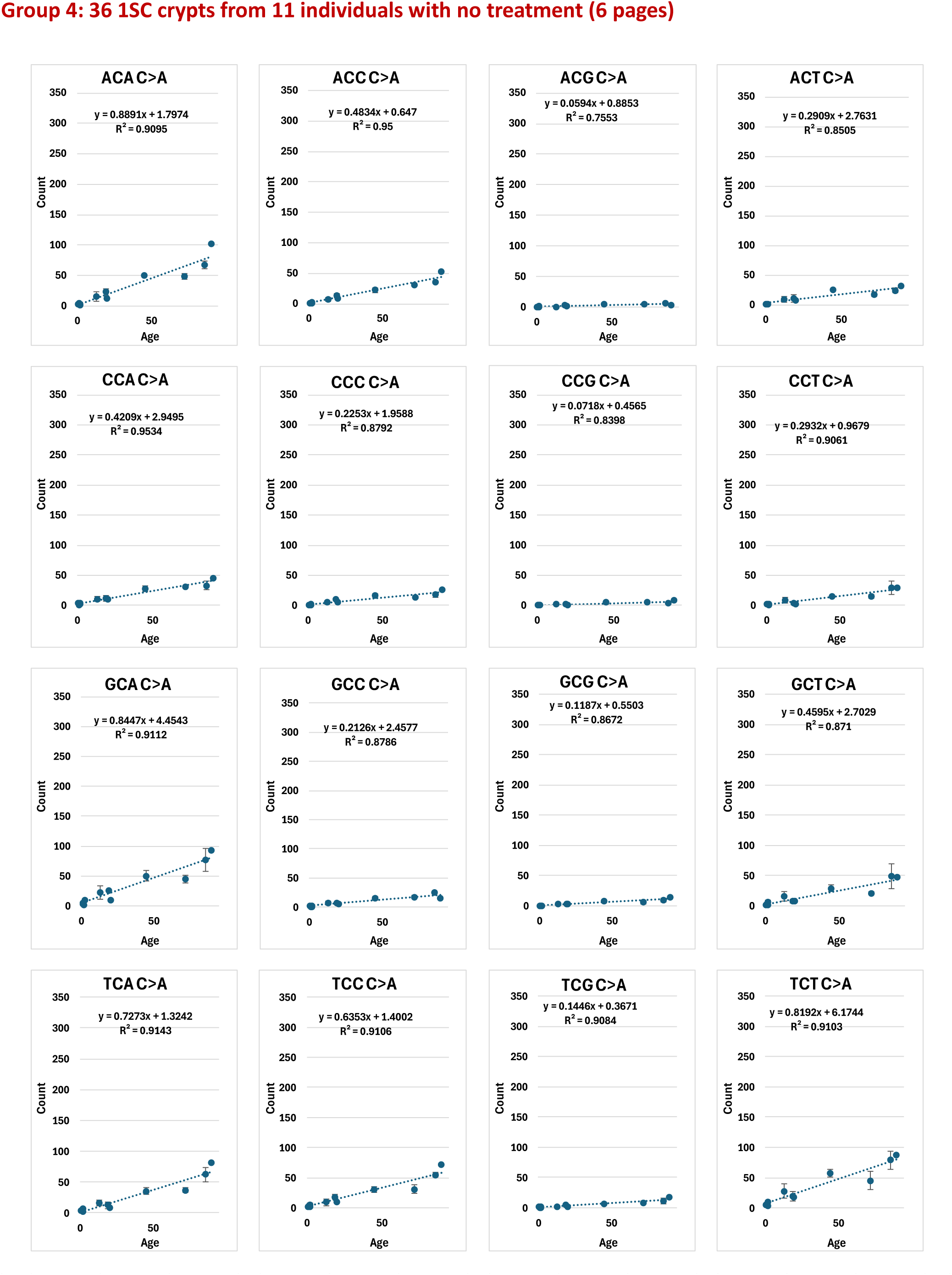
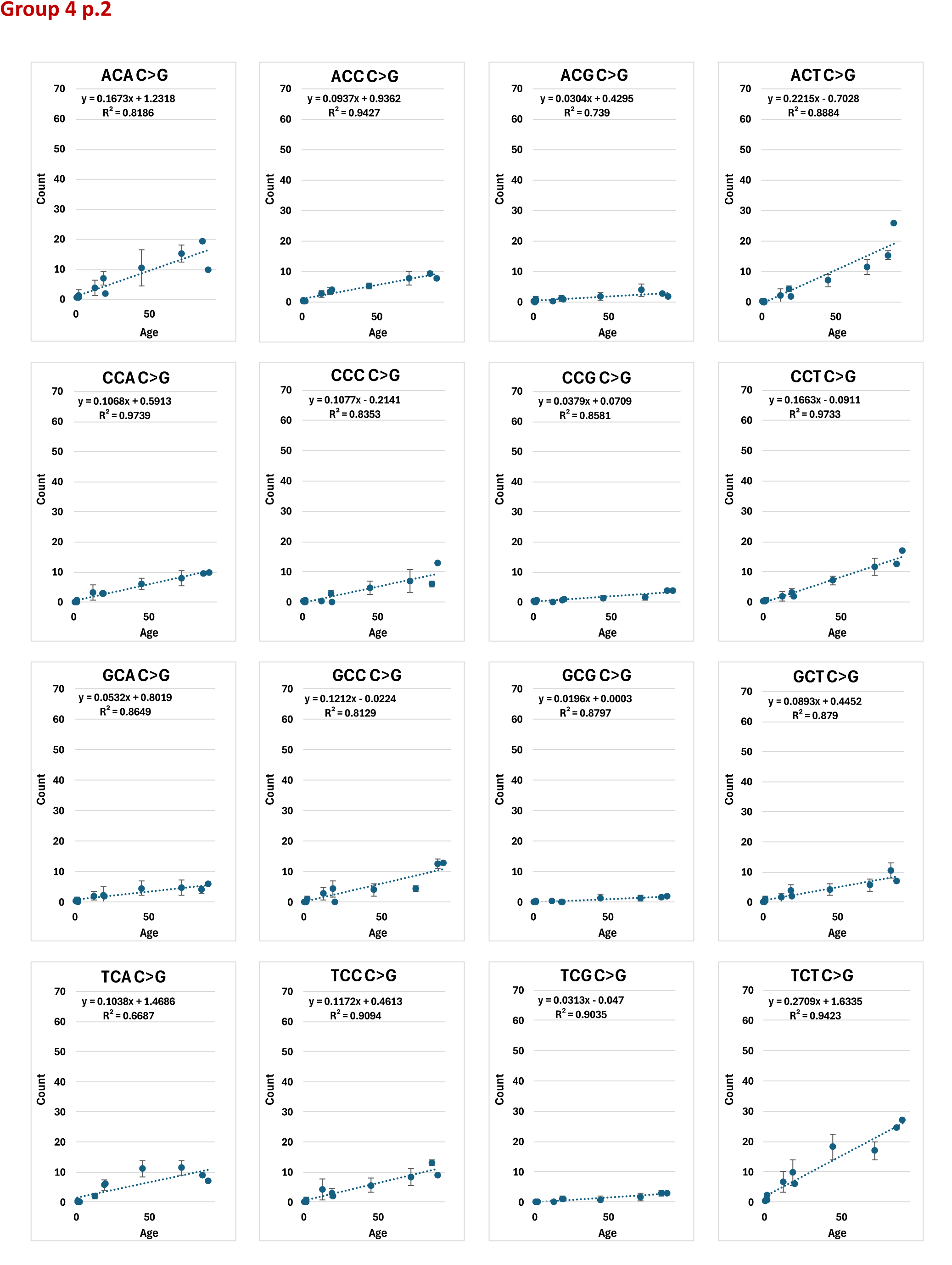
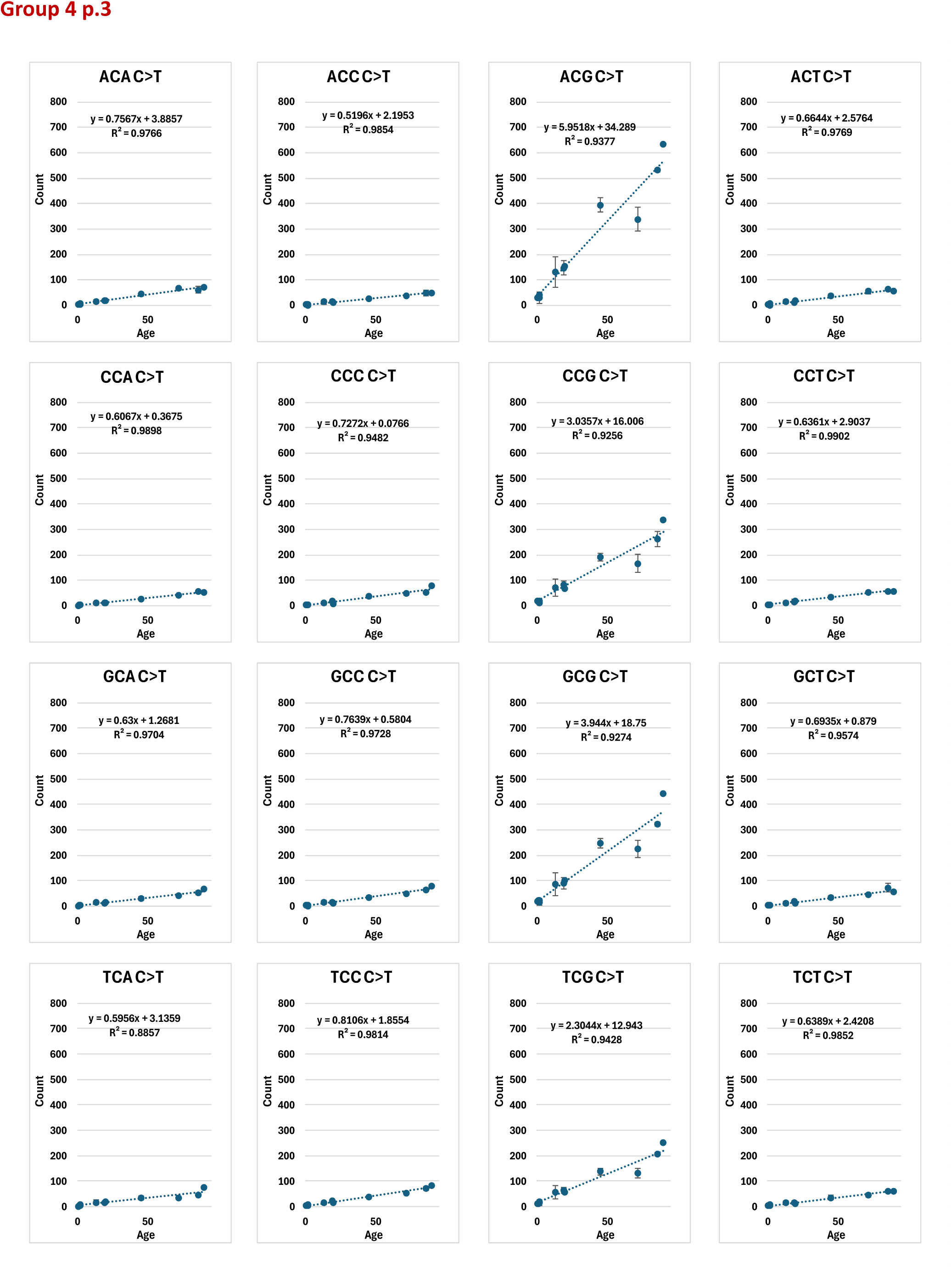
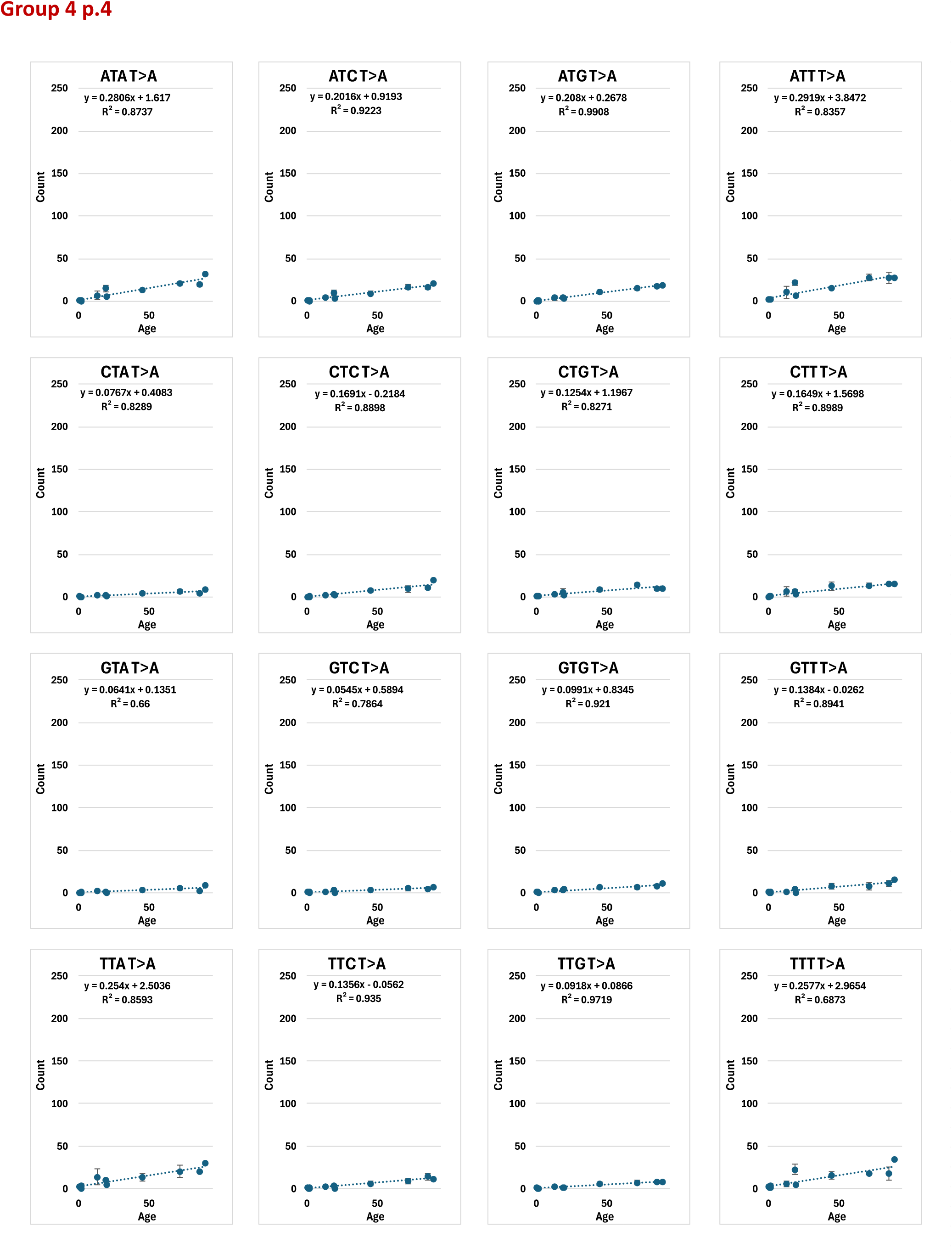
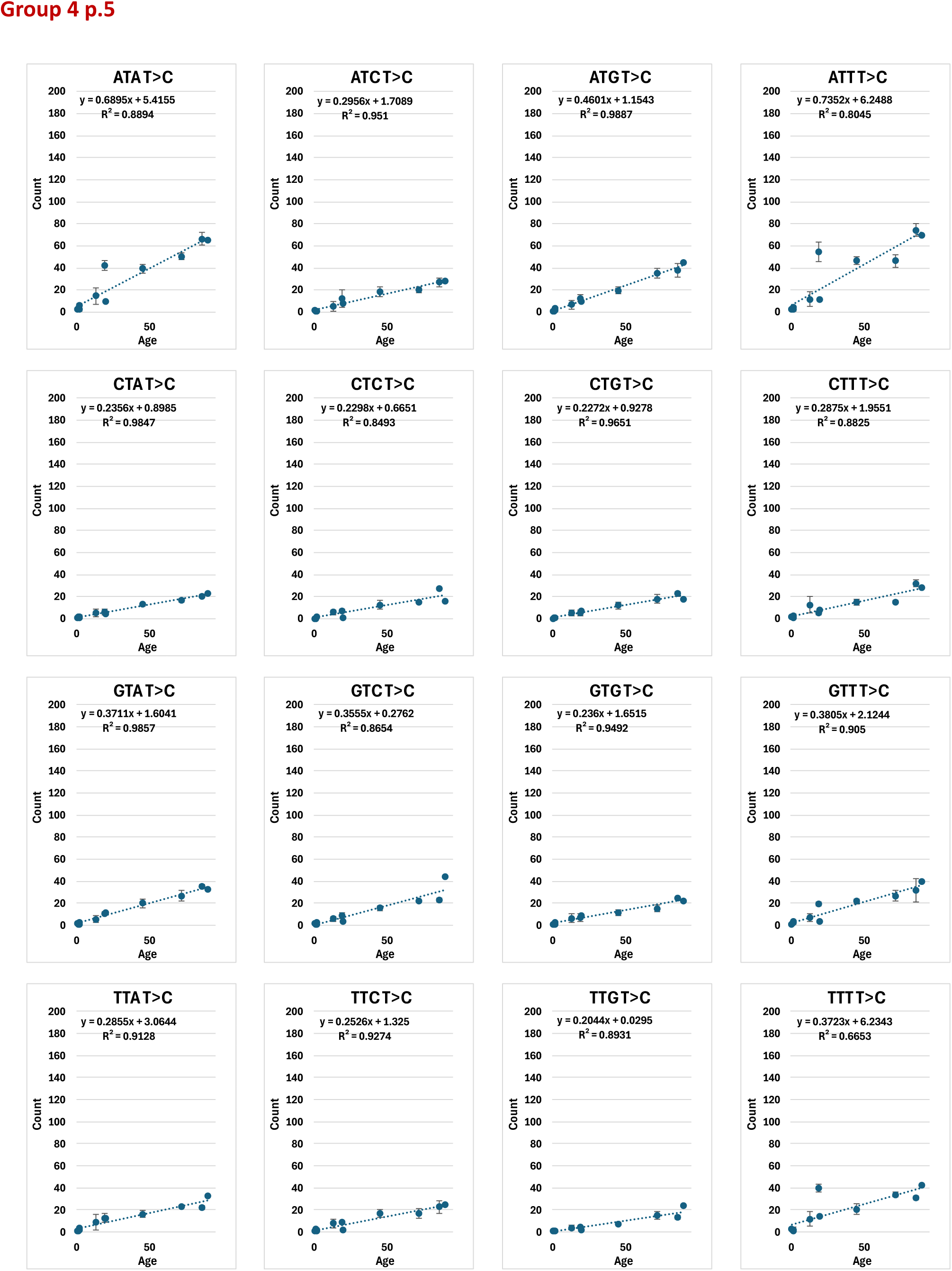
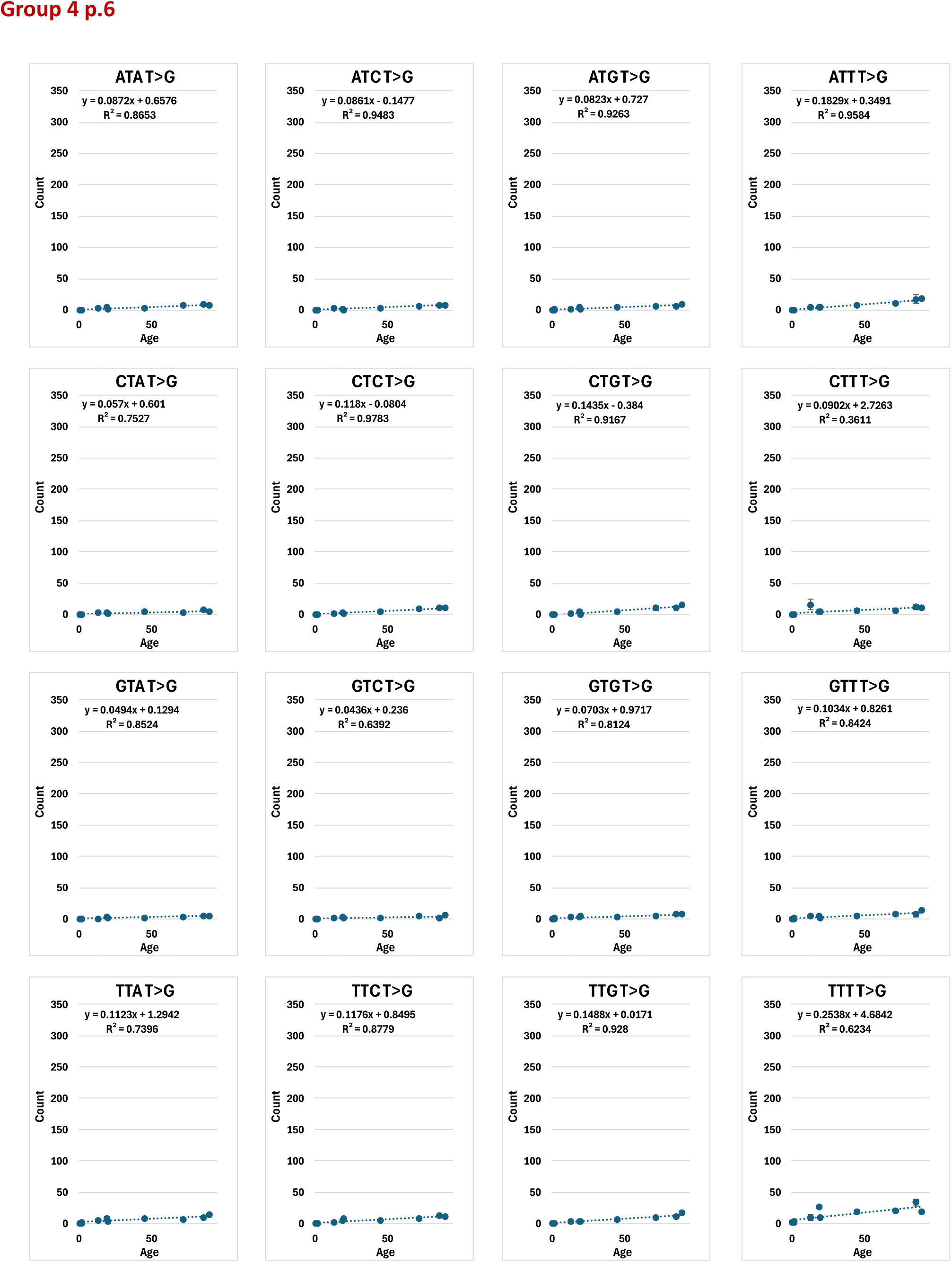
Mutation count by age for each of the 96 tri-nucleotide categories.

**Supplementary Figure S3.**
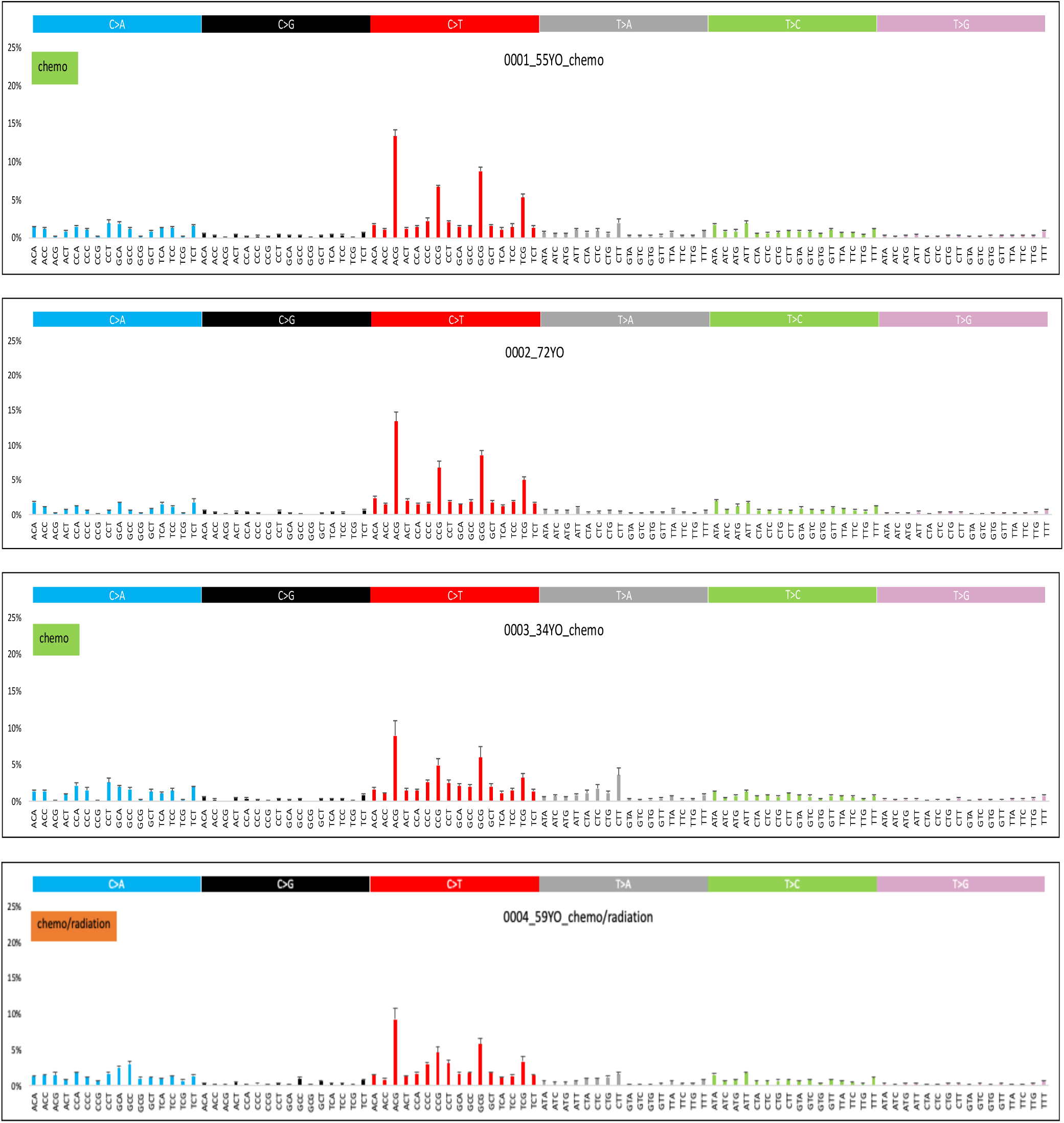
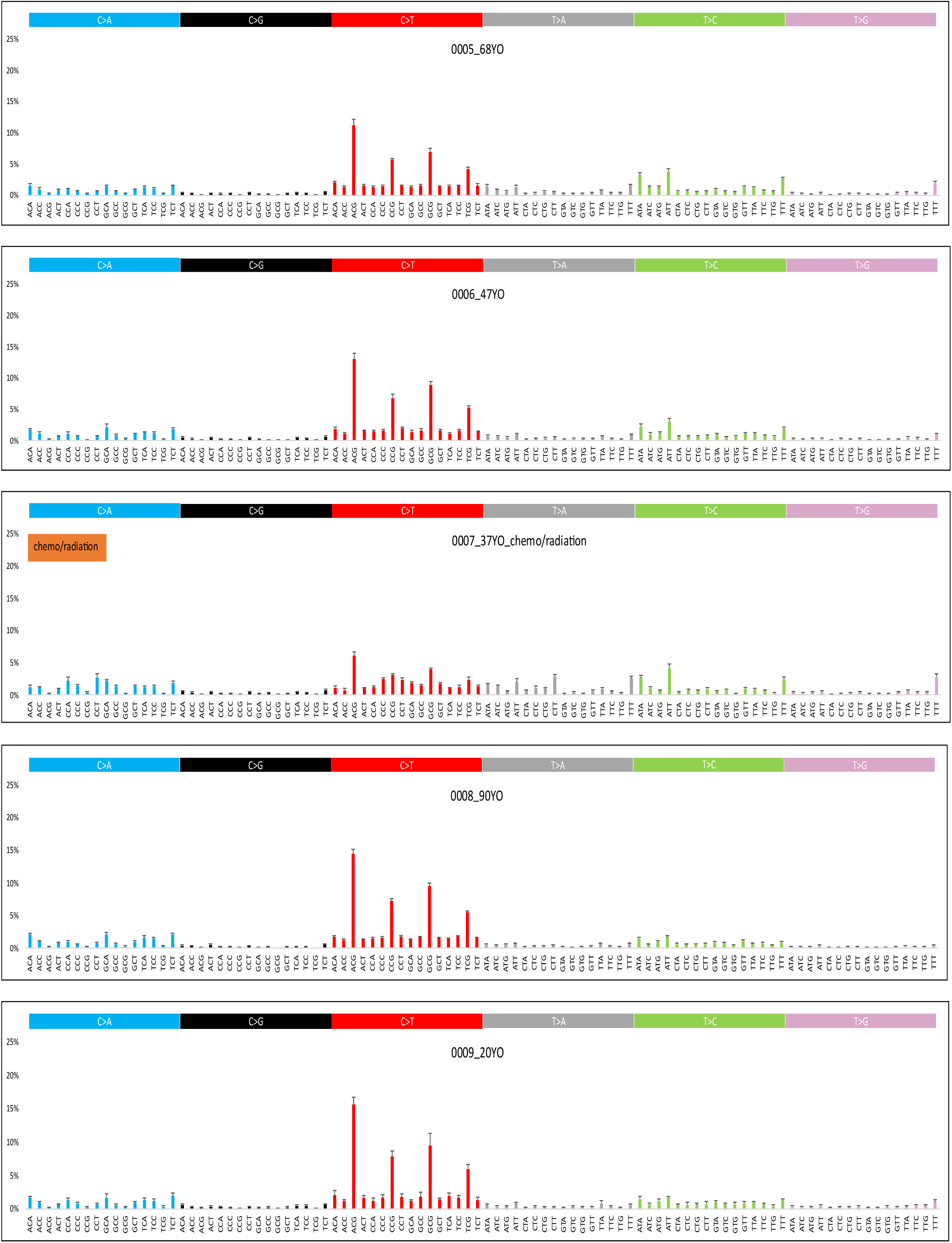
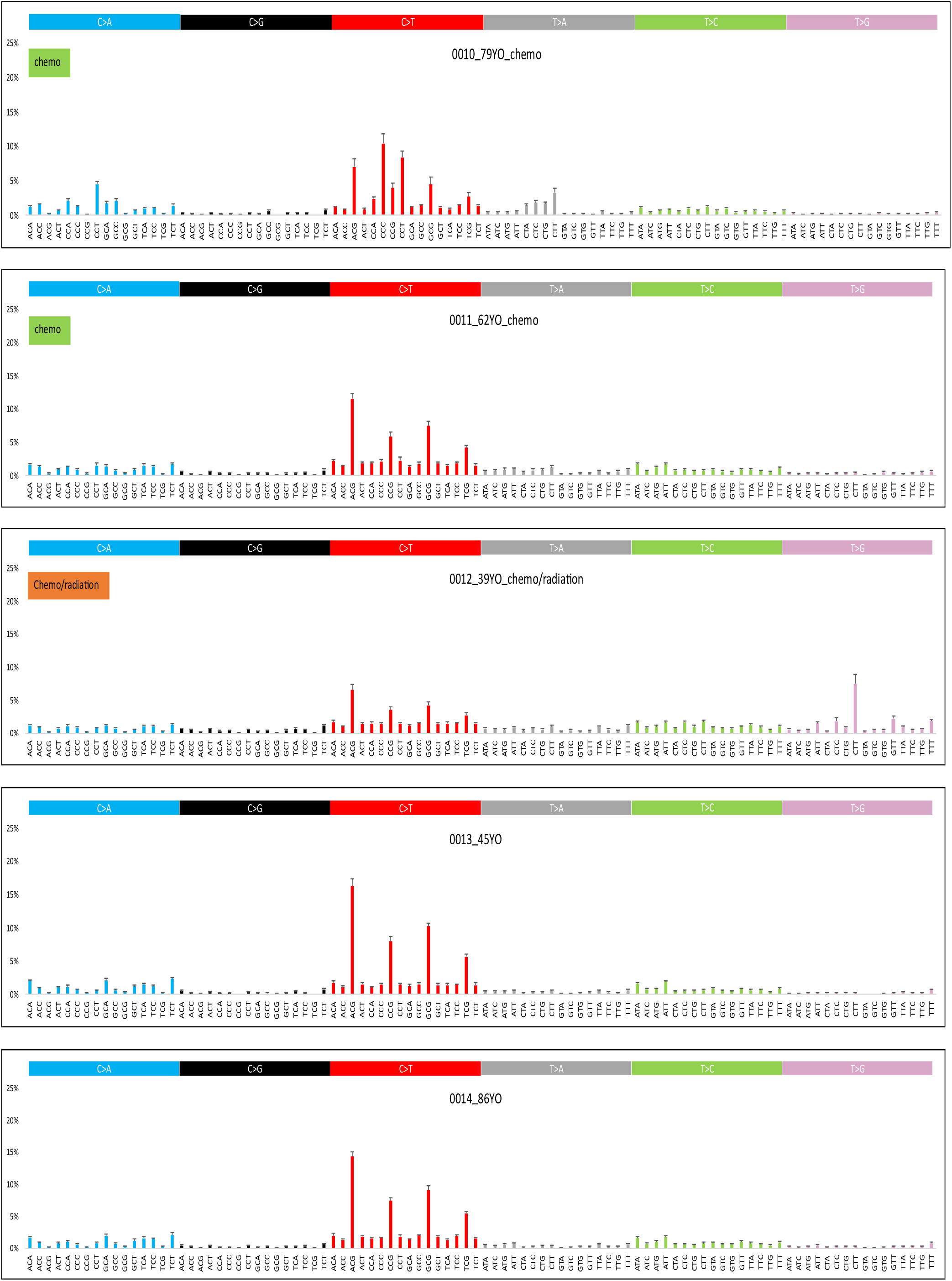
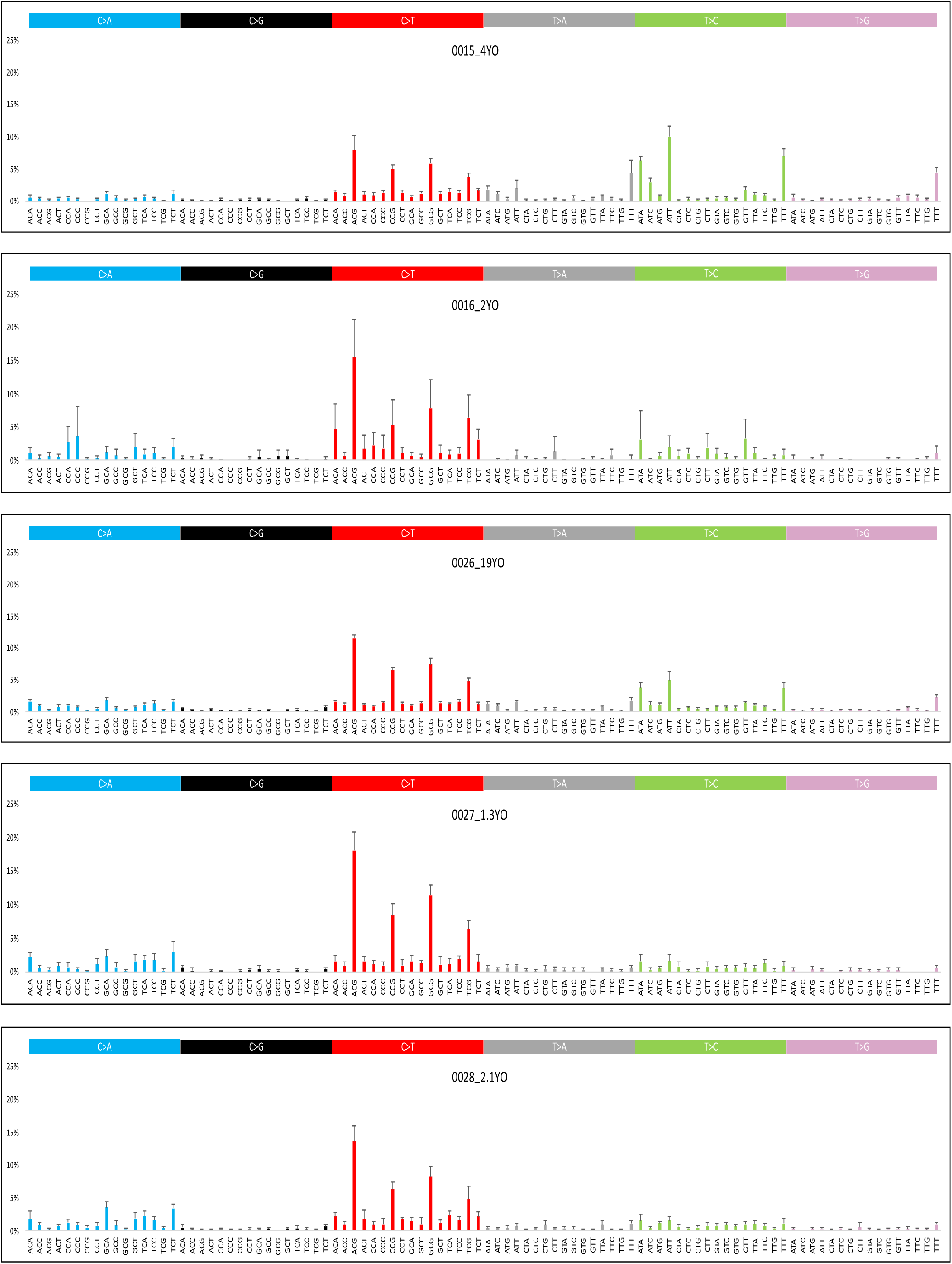
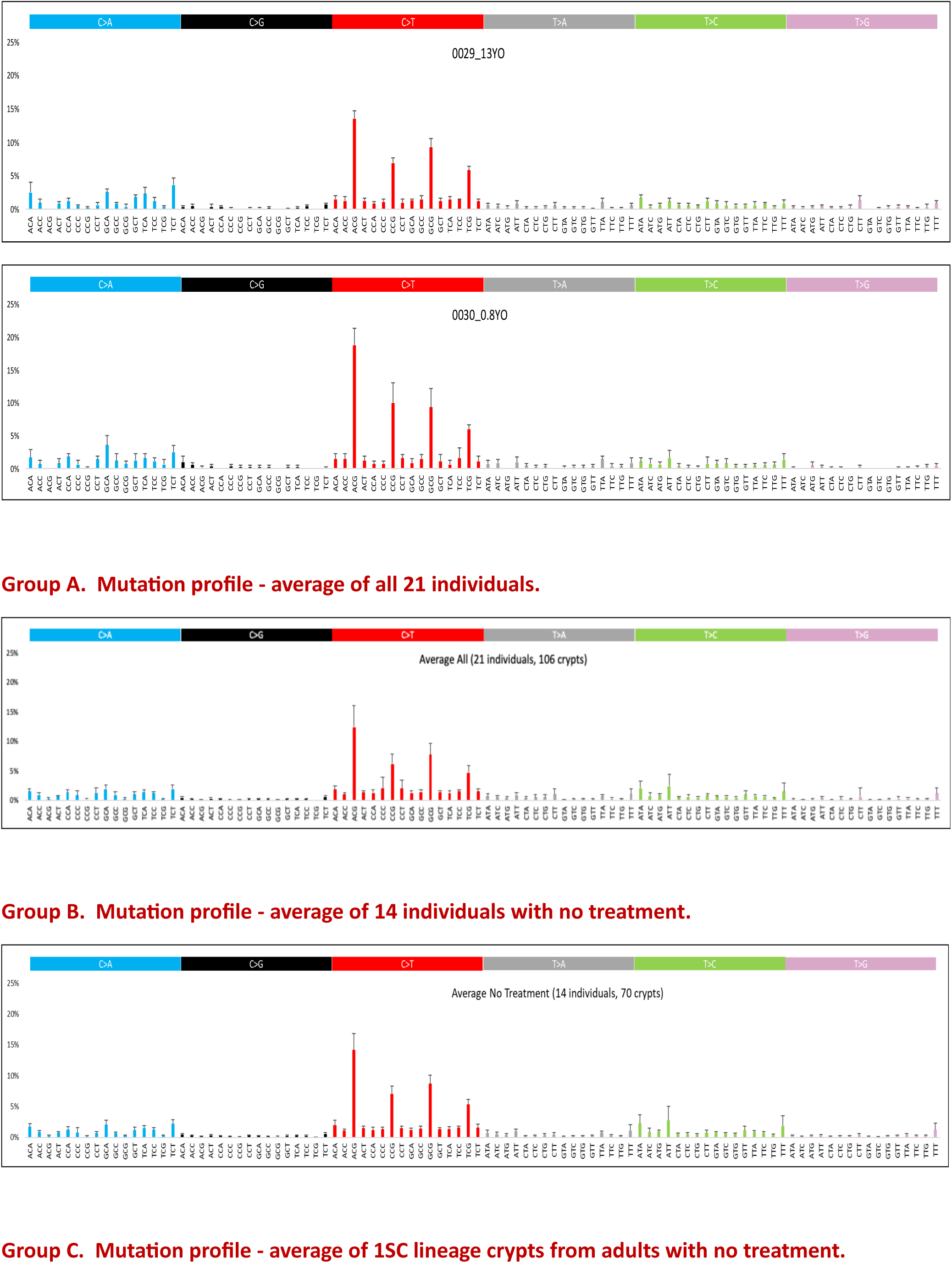
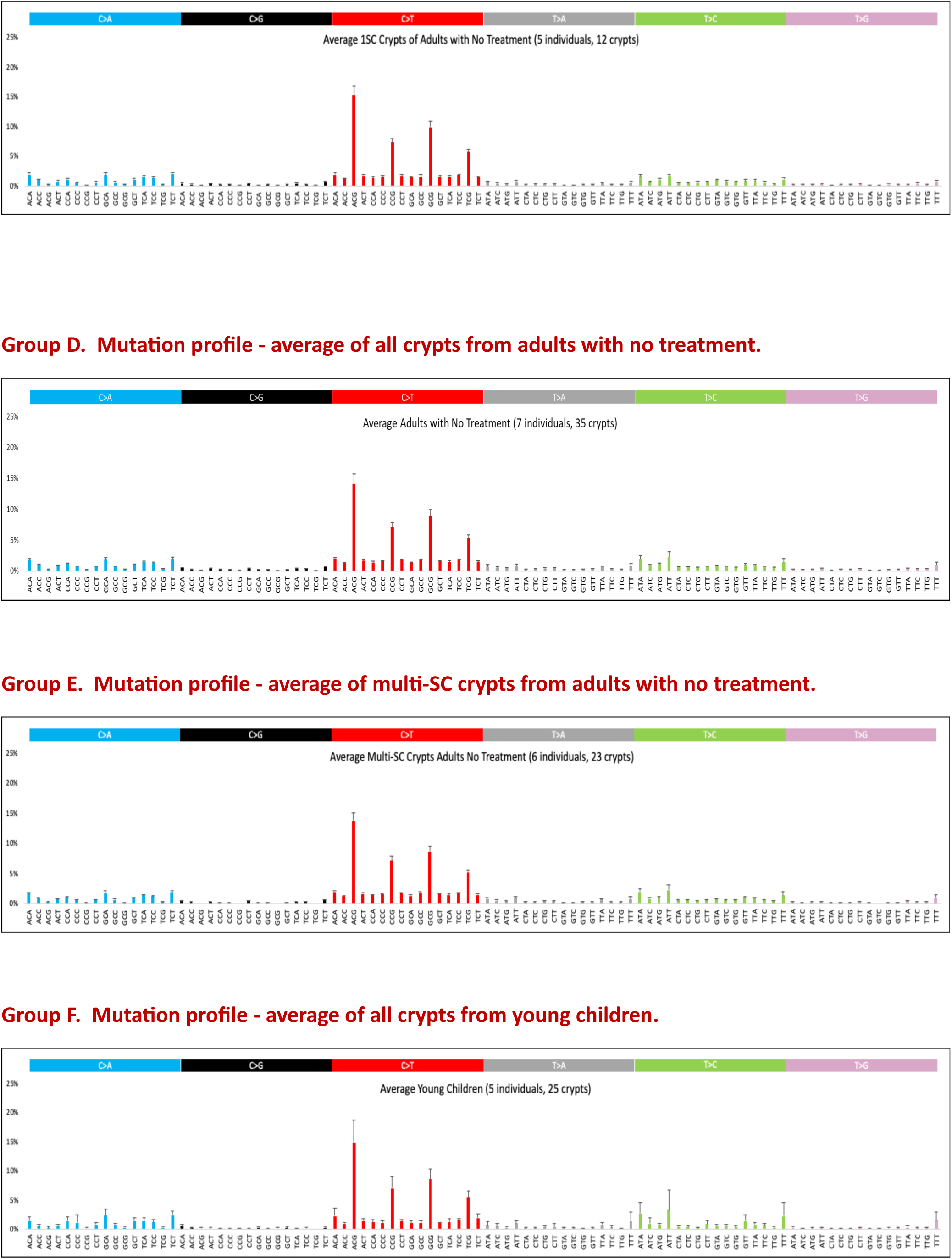
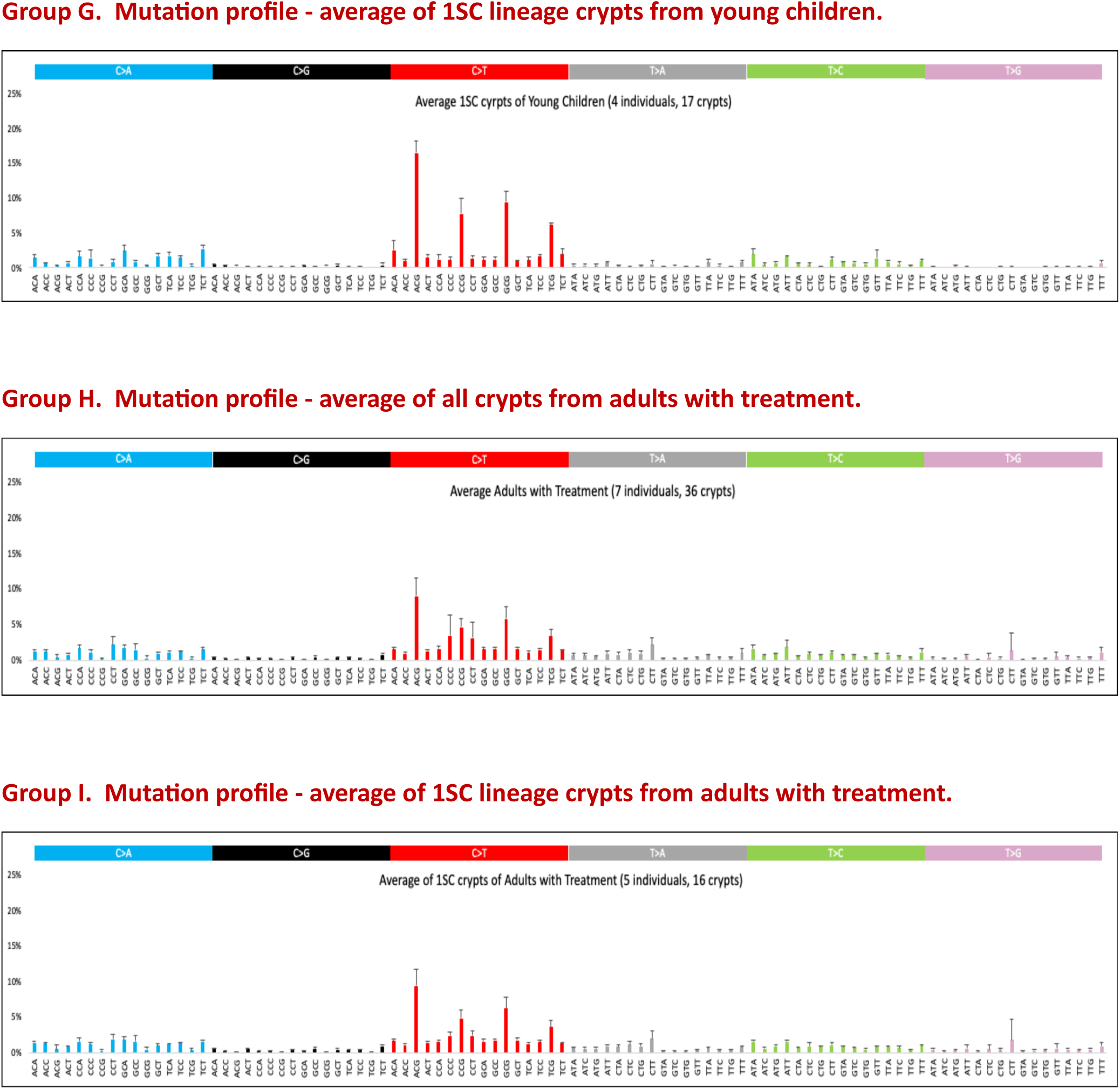
Mutation profile for each of the 21 individuals presented as 96 trinucleotide mutation categories (individuals received treatment as marked), and mutation profiles from average of indivisuals in Groups A to I.

**Figure S4.**
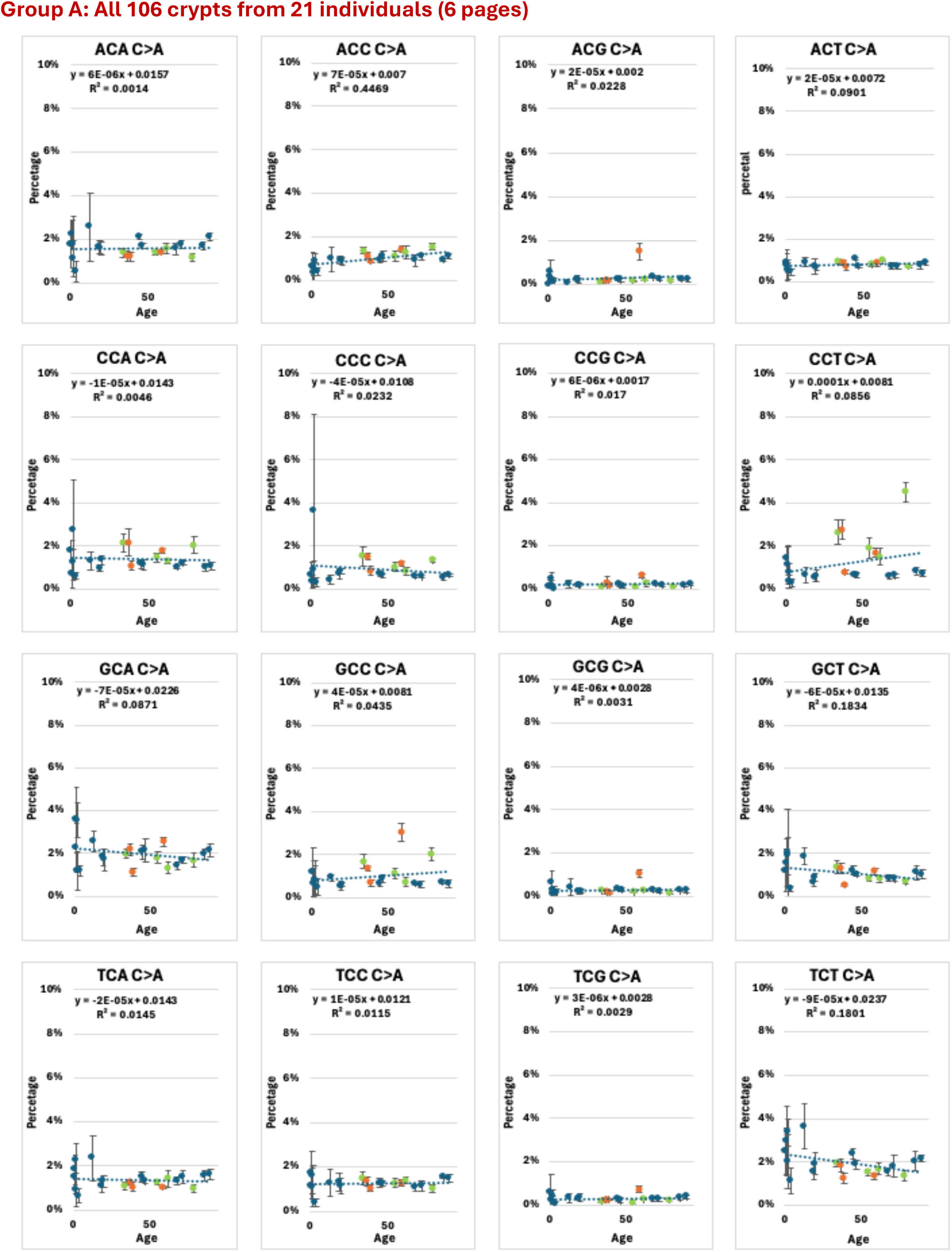
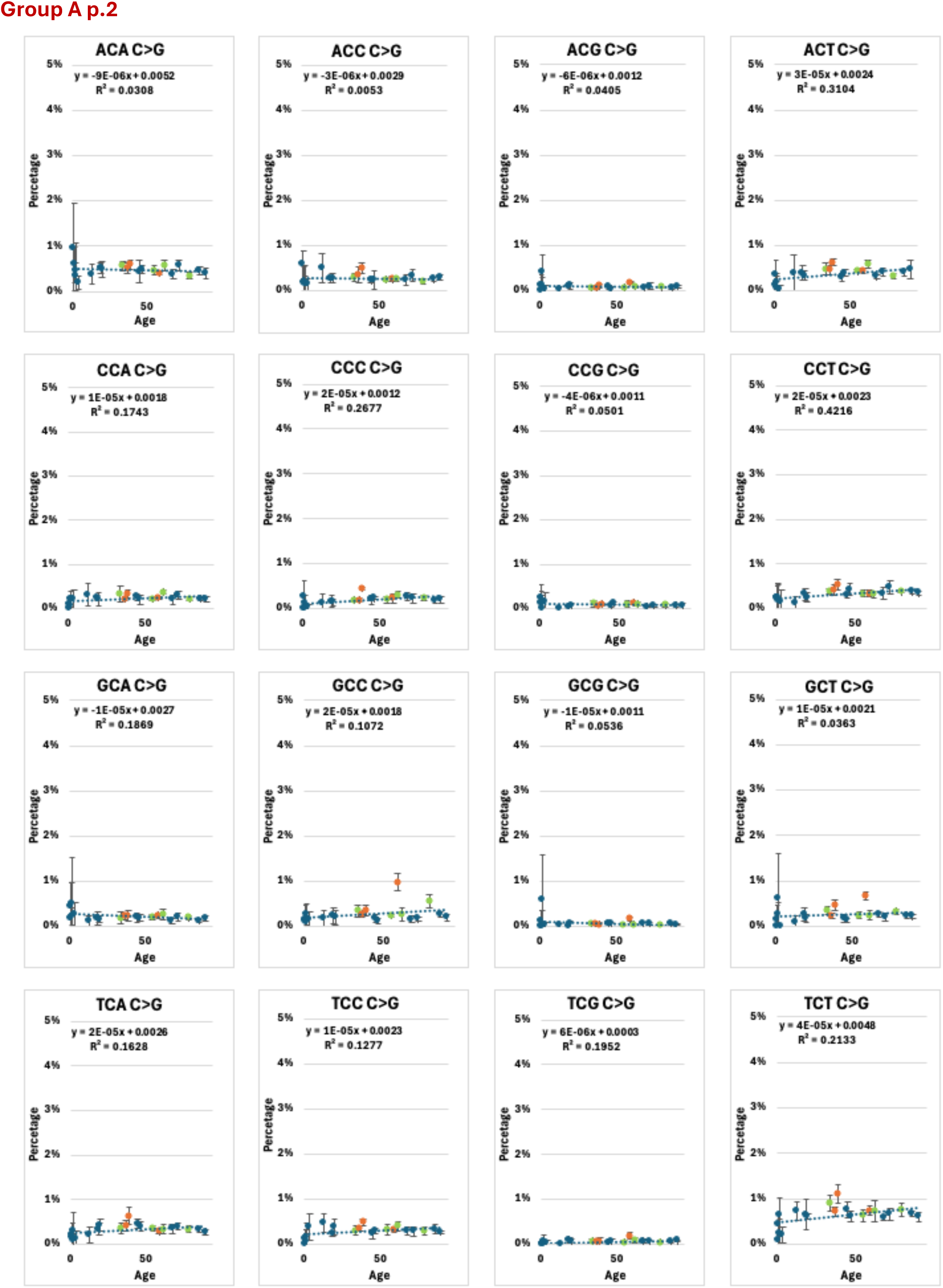
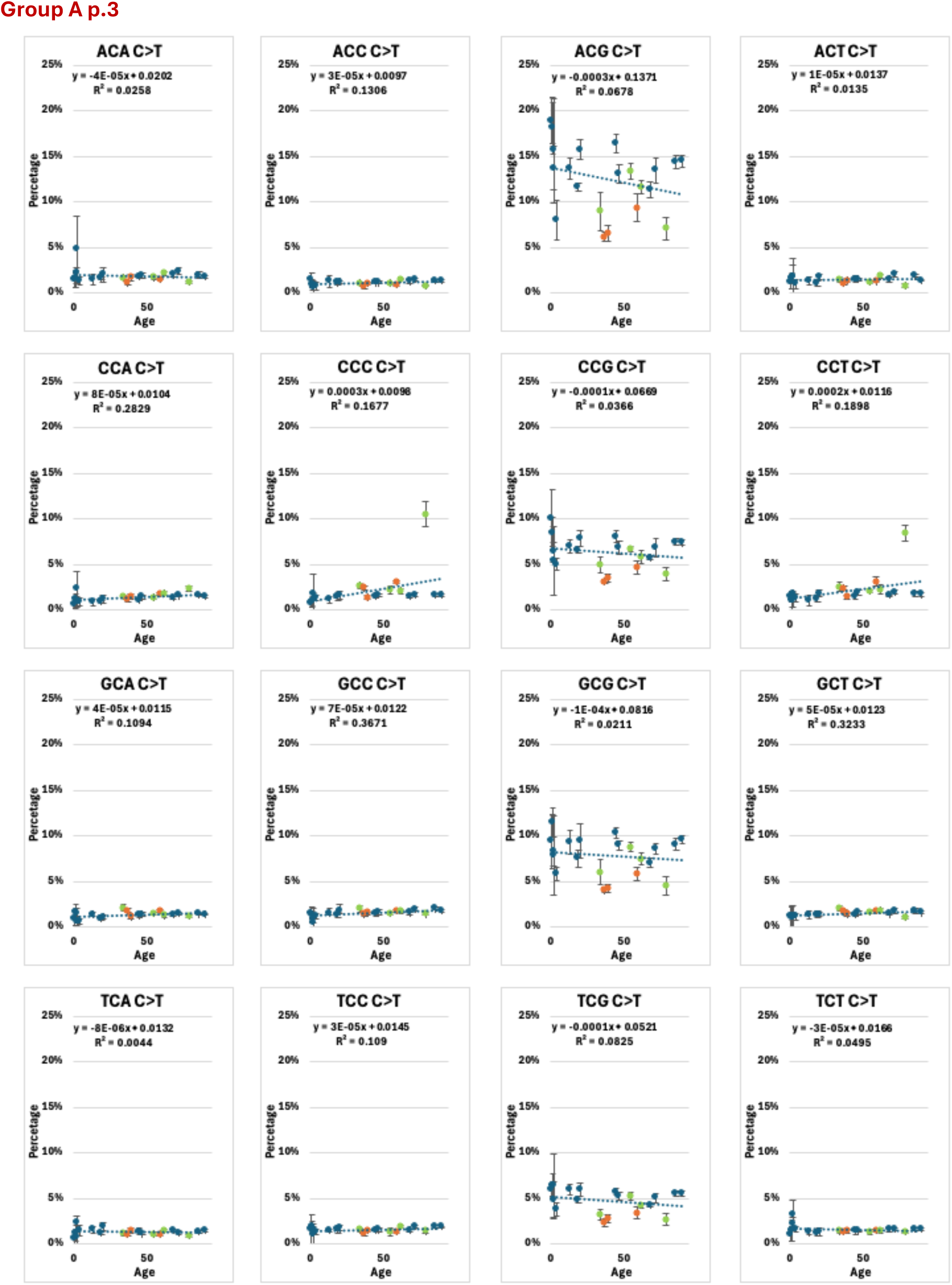
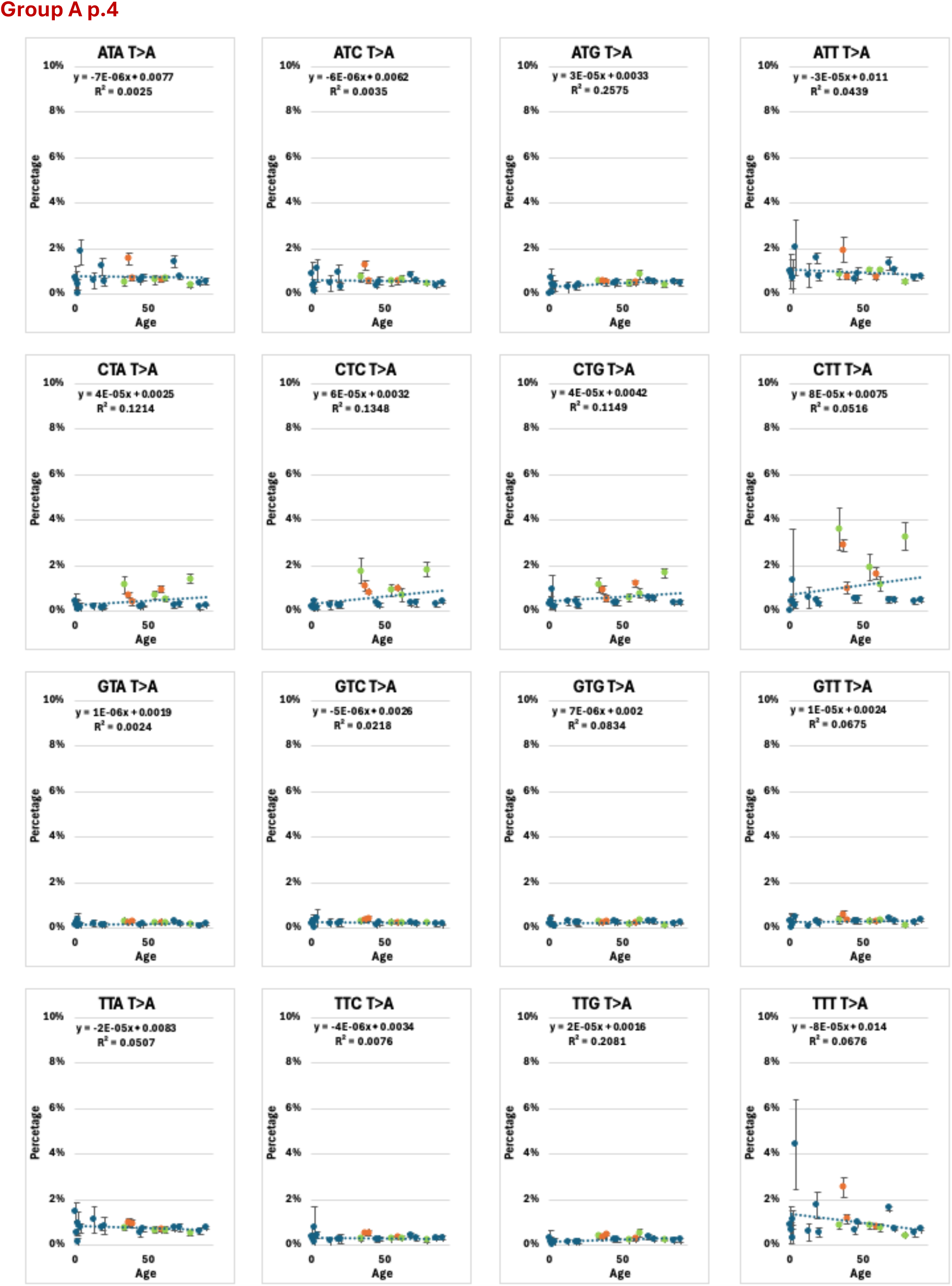
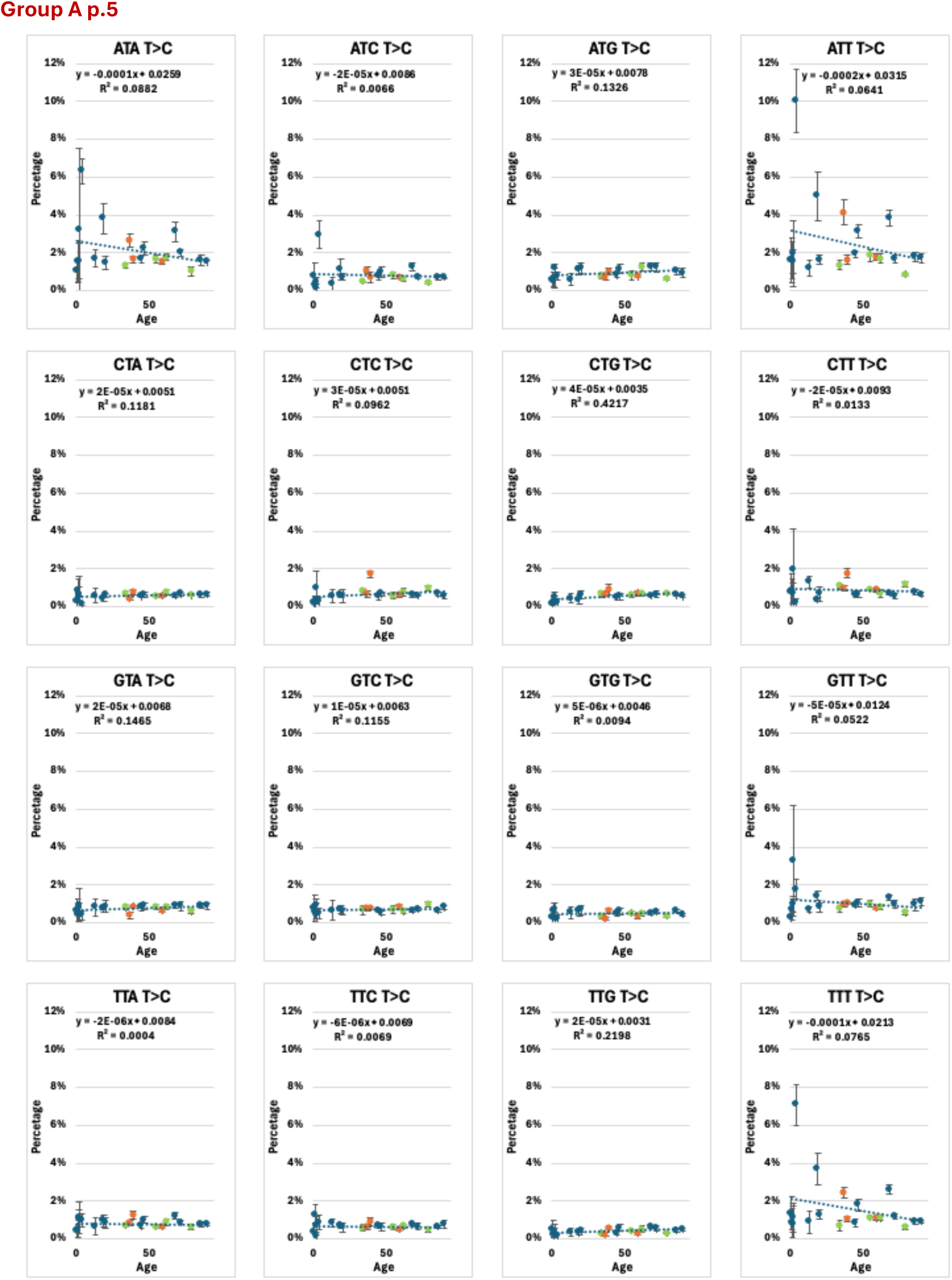
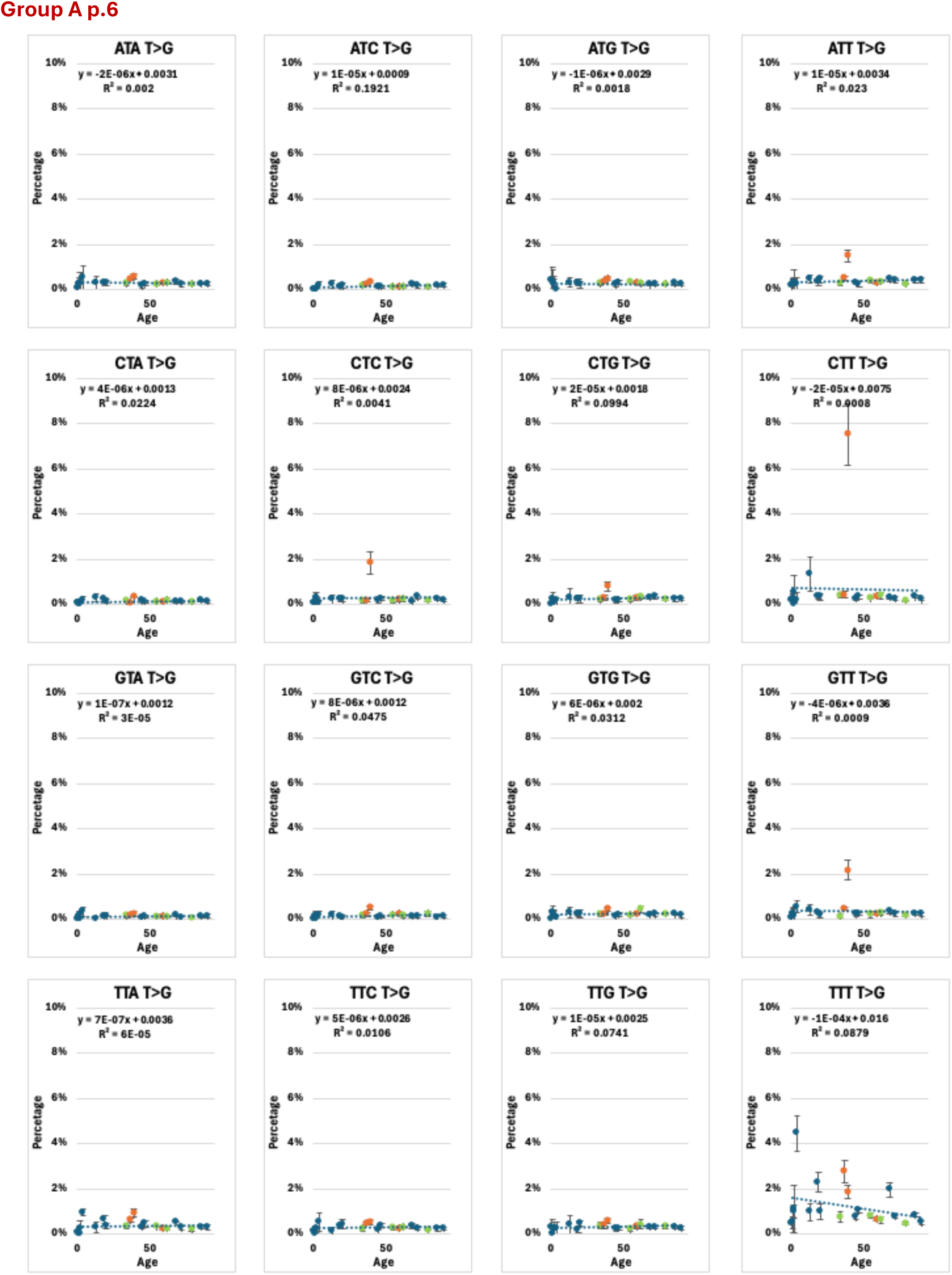
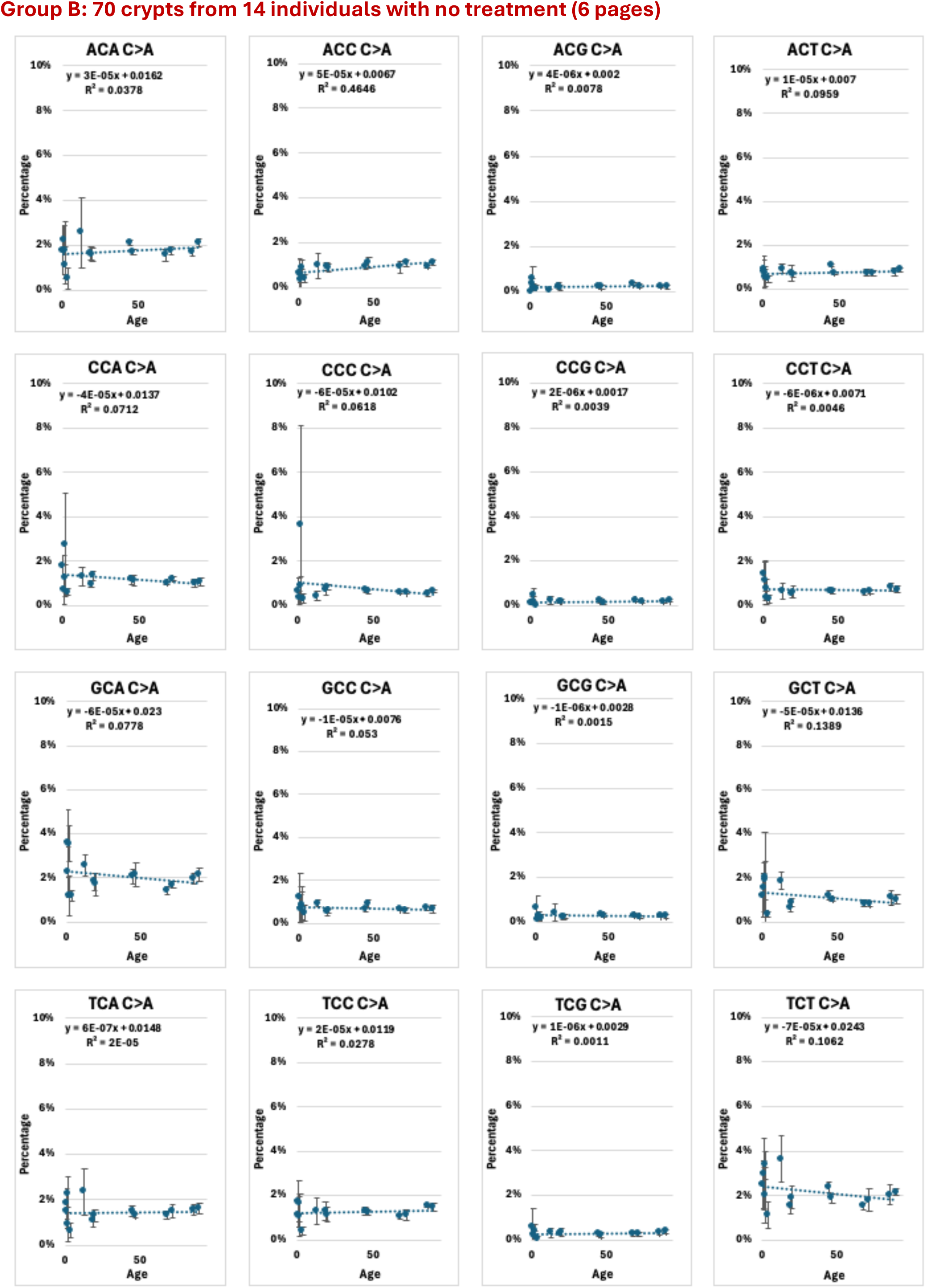
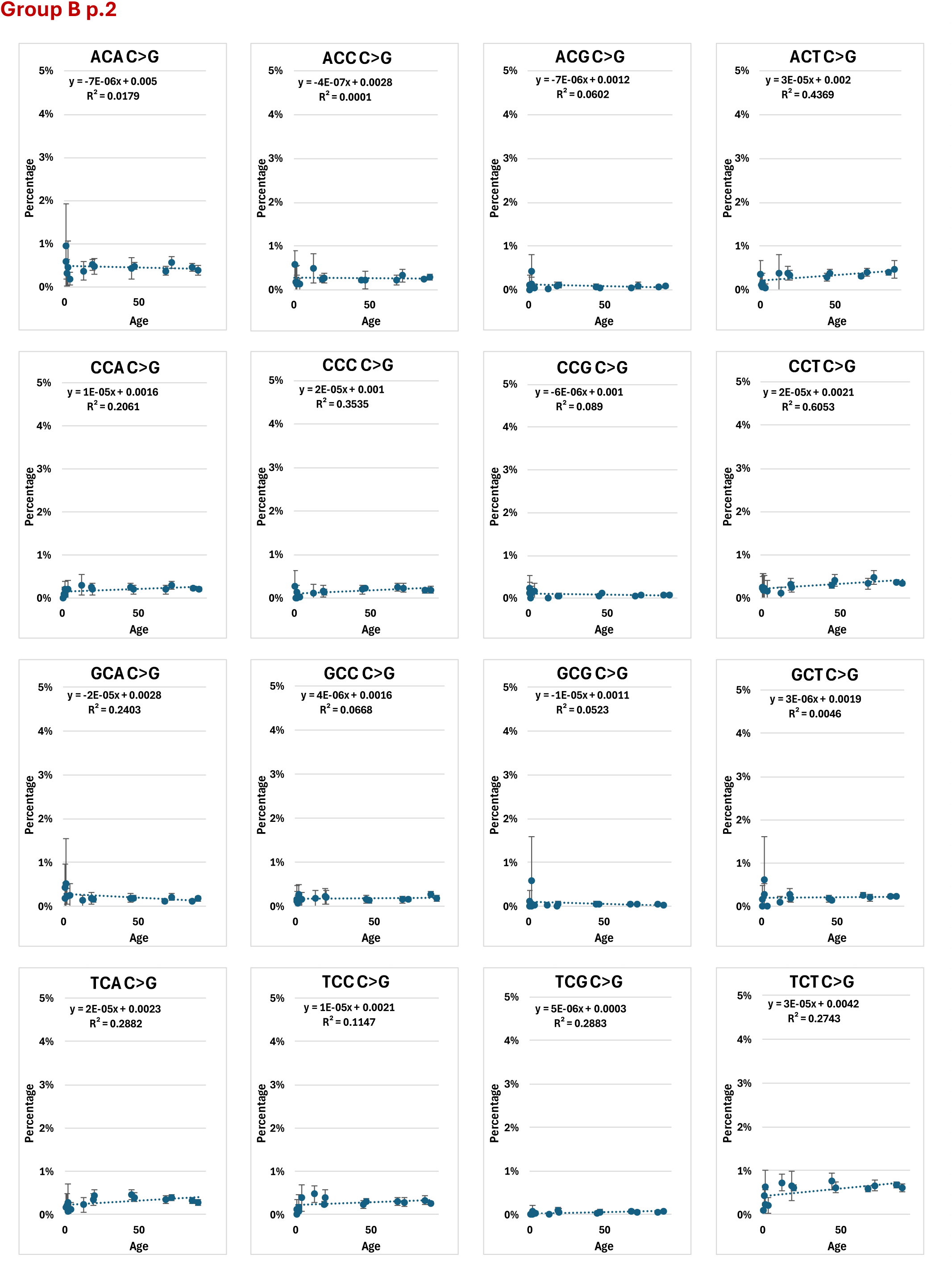
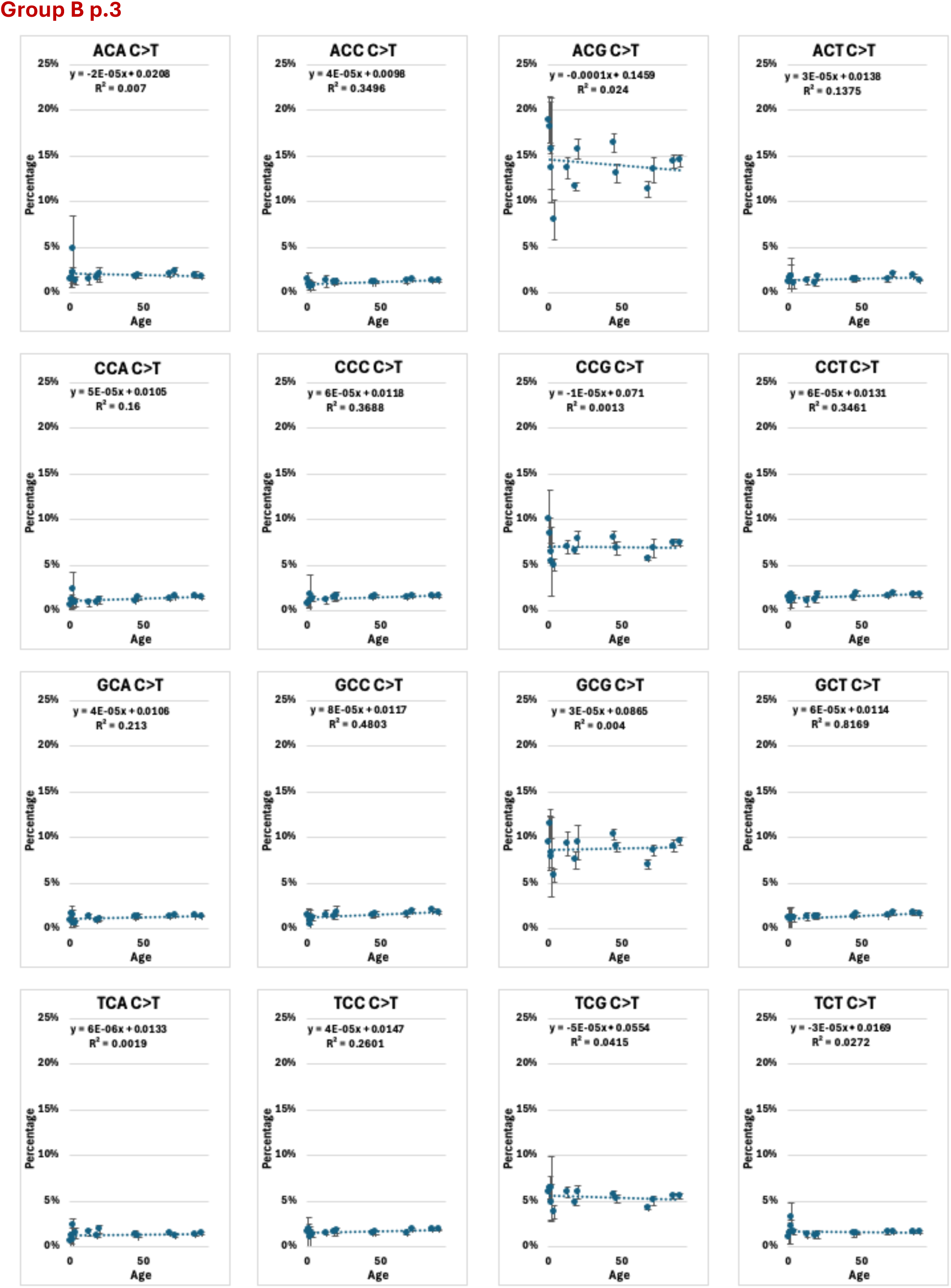
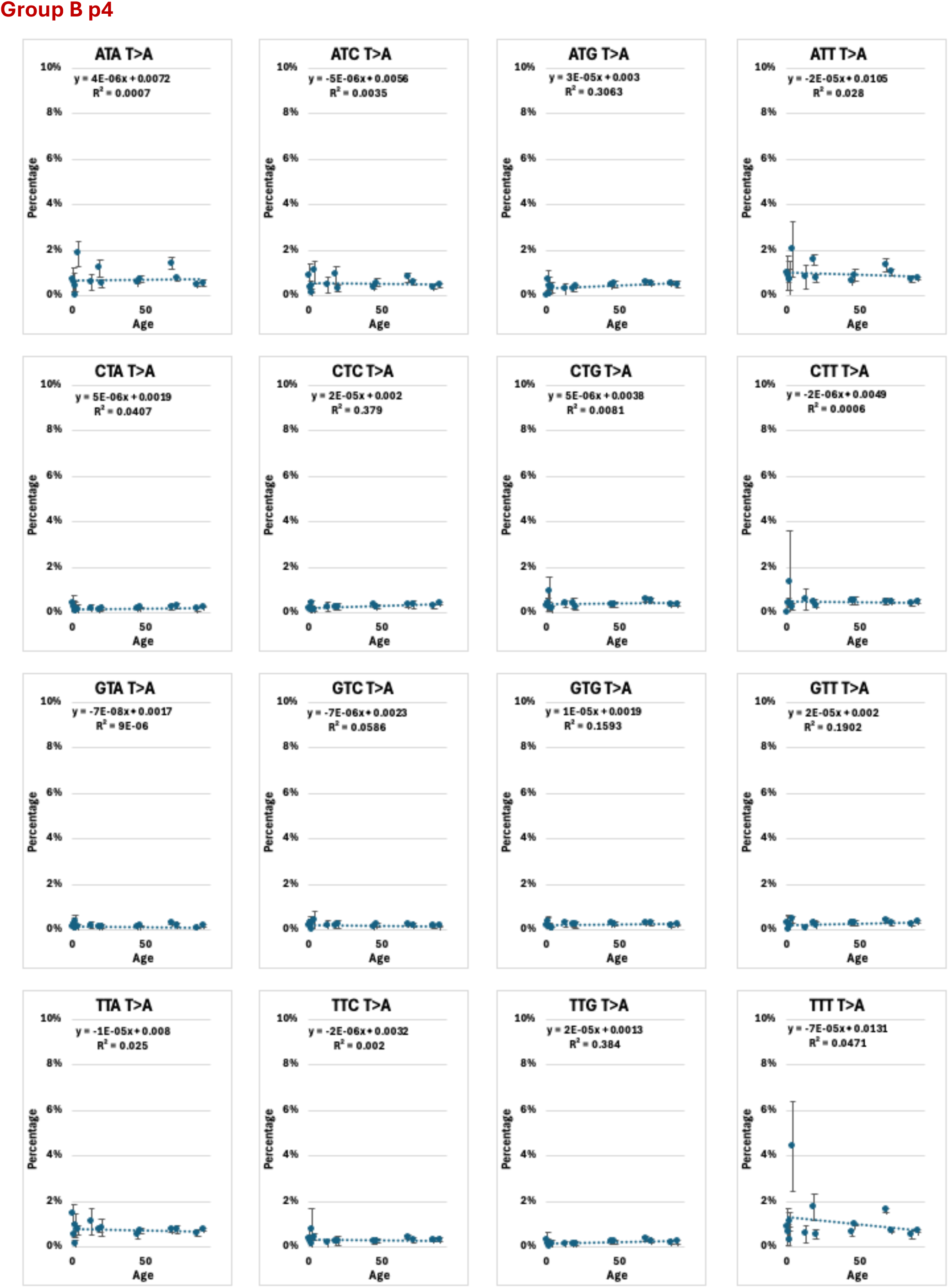
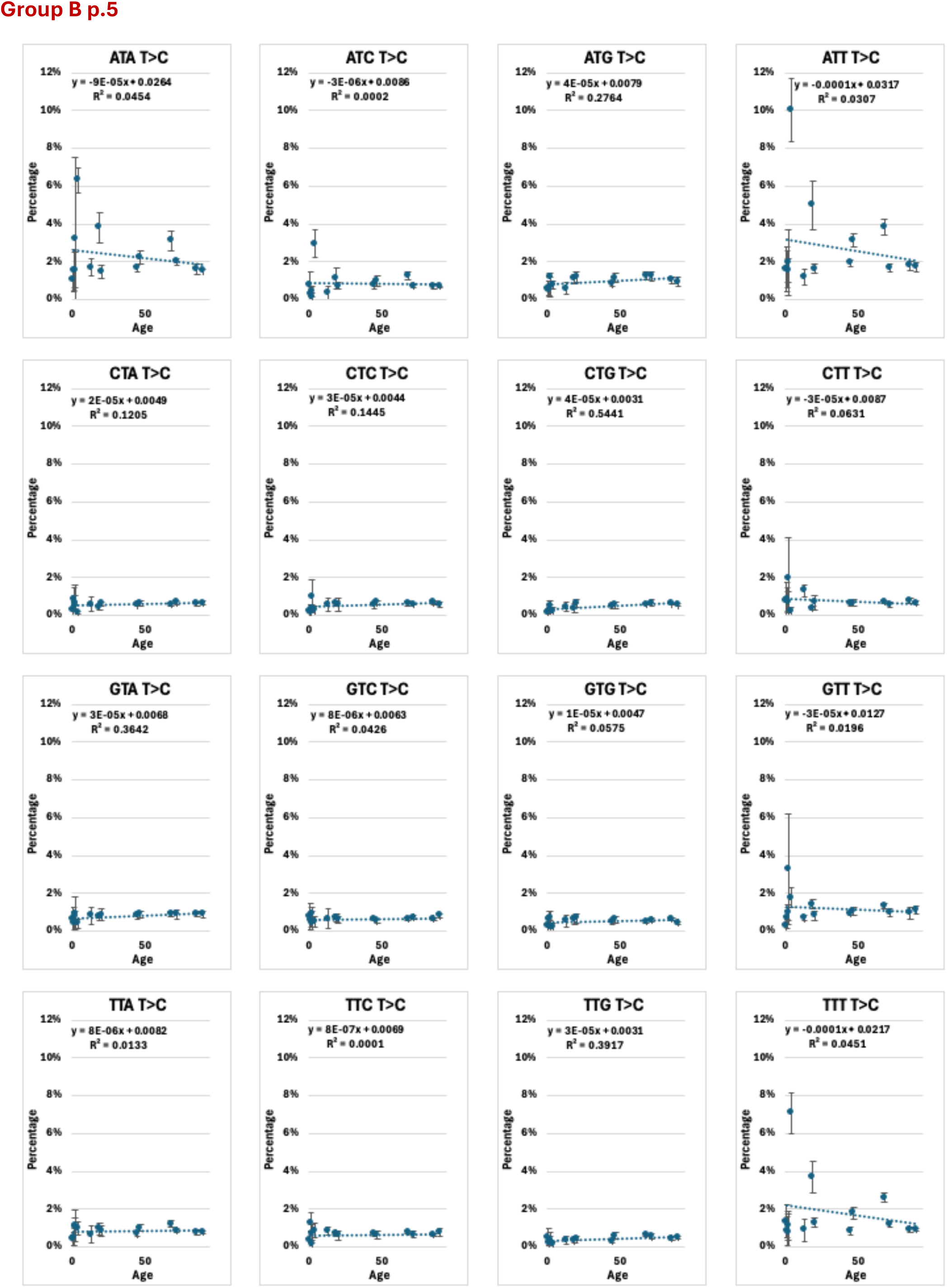
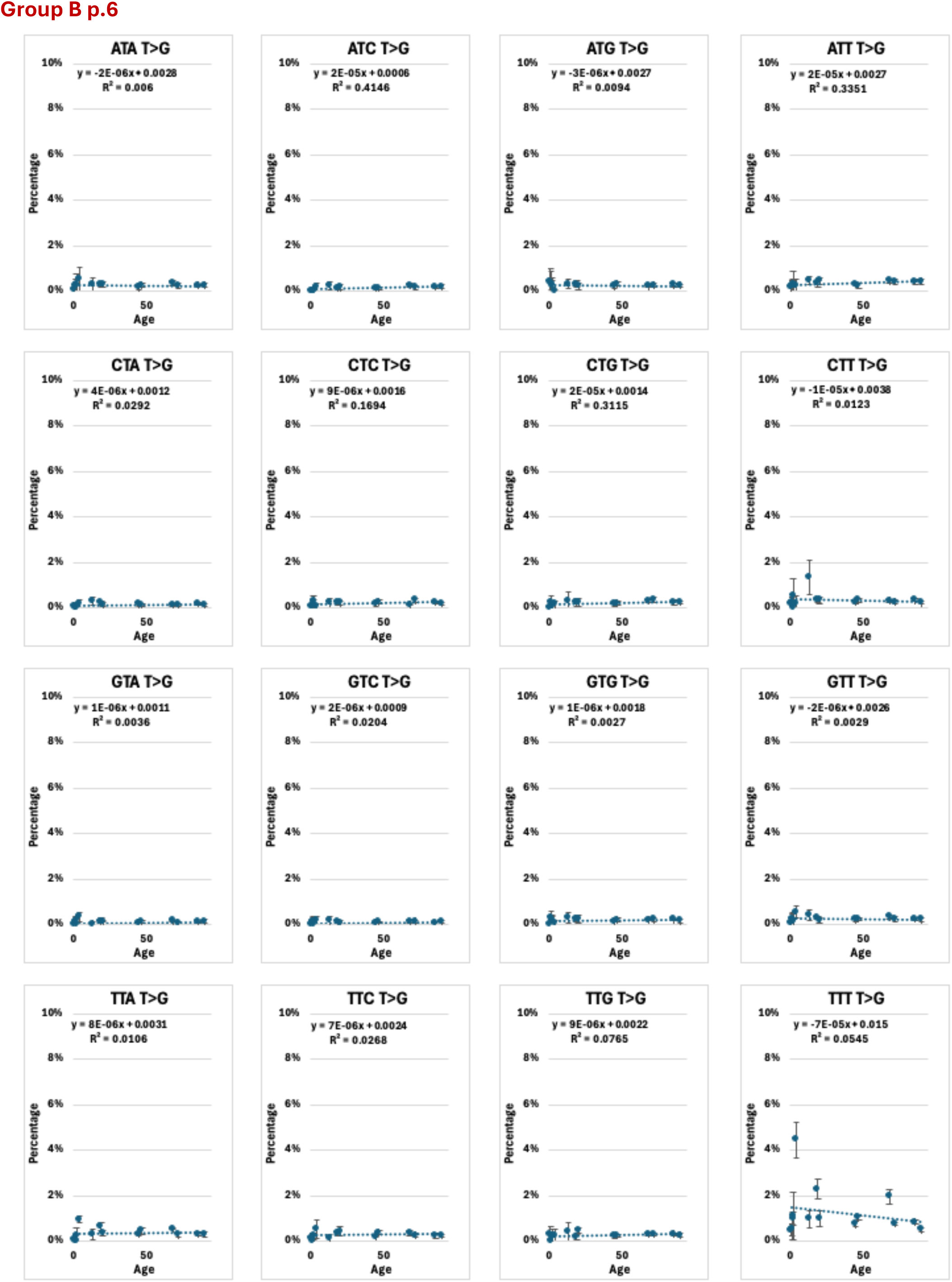
Mutation frequency of each mutation categories by age.

